# Tetrapod terrestrialisation: a weight-bearing potential already present in the humerus of the stem-tetrapod fish *Eusthenopteron foordi*

**DOI:** 10.1101/2024.02.09.579723

**Authors:** François Clarac, Alexis Cornille, Sifra Bijl, Sophie Sanchez

## Abstract

Our study shows that the von Mises stress, induced by external load on the humerus of Eusthenopteron, dissipates through the cortex, trabeculae and the muscles of the pectoral appendage involved in elevation and protraction. As Eusthenopteron’s microanatomy is similar to that of Devonian tetrapods, we expect them to share the same process of load dissipation and energy absorption through 1) cortical stress distribution; and 2) longitudinal trabecular conduction. Our FE simulations in hypothetical terrestrial conditions demonstrate that this type of microanatomical architecture could withstand the weight of Tiktaalik proportionally to the size of Eusthenopteron in standing posture. This tubular arrangement, including marrow processes originally involved in long-bone elongation, would have acquired a key secondary biomechanical function to increase the resistance and strength of the cancellous bone to external compressive load. As an exaptation, this specific trabecular architecture may have played a major role in the tetrapod land exploration about 400 million years ago.

## INTRODUCTION

Tetrapods have displayed a remarkable ability to adapt any types of environments. They have evolved a great diversity of specialised morphologies to perform fast motion in water, on land, or in the air. The fin-to-limb transition (Fig 1), that occurred about 400 million years ago, has played a major role in this radiation. Despite the high interest of the biology community for this crucial evolutionary event, it has been a challenge to date and characterise the terrestrial locomotory behavior of early tetrapods. In the last decades, the development of state-of-the-art biomechanical techniques/simulations (1–6) and the discovery of new tetrapod trackways (7–9) has relaunched this interest. This started with the publication of Niedzwiedzki et al. (7) proposing an origin for tetrapods deeper than previously suggested, into the early Middle Devonian. Trackways in a shallow-water environment were assigned to tetrapods in the Middle Devonian locality of Zachelmie in Poland. The fossil evidence revealed a gait symmetry for these tetrapods, as in terrestrial locomotion. This discovery drastically shifted back tetrapod “parasagittal” locomotion to the Middle Devonian. Using a computer-aided assessment of limb joint mobility, Pierce et al. (1) casted doubt on *Ichthyostega*-like tetrapods as the trackway makers in the locality of Zachelmie. They demonstrated that the early tetrapod *Ichthyostega* had a restricted shoulder and hip join mobility, unlikely to be able to push the body off the substrate and perform alternating limb movements to produce symmetrical gait prints. Dickson et al. (5) used functional adaptive landscapes on the tetrapod humerus to predict terrestrial ability of the tetrapod forelimb and suggested that early tetrapods may have used “transitional gaits” (such as crutching based on belly dragging) to explore land. Kawano and Blob collected Ground Reaction Forces (GRF) using a multi-axial force plateform on various modern tetrapods, to characterise these “transitional gaits and postures” in early tetrapods (2). They highlighted the mosaic of fish-like and tetrapod-like kinematic features in these taxa based on the inference of peak GRF magnitudes (10). Nyakatura et al. used close techniques to quantify the 3D kinematic components of limbs in dragging-belly conditions (4). On this basis they could propose an evolutionary scenario where limb propulsion preceded body weight bearing in tetrapods. The strength of the pectoral appendages and shoulder of finned stem tetrapods, such as *Tiktaalik*, was confirmed and featured by Hohn-Schultze et al. (3) by applying Finite Element Analysis (FEA) to map compressional stress peaks during hypothetical standing, crawling and walking on land. Even if these various investigations do not always agree on all aspects of the early locomotion of tetrapods, they all converge toward the hypothesis of an early ability of the pectoral appendages to a certain form of terrestriality (at least crutching) in stem tetrapods.

**Fig 1.**
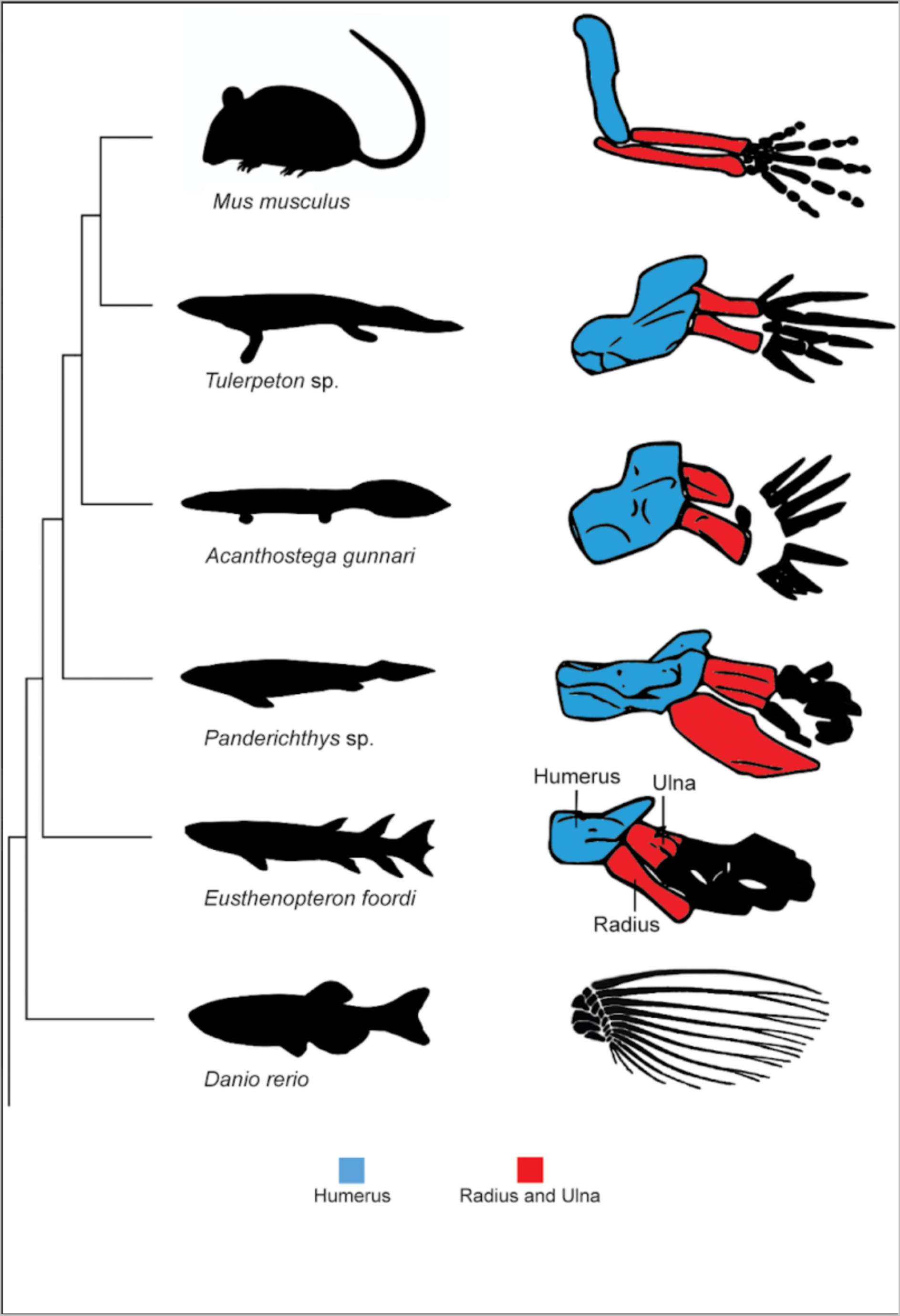
Simplified phylogeny of the tetrapodomorphs around the “fin-to-limb” transition. The homology between the appendicular bones is highlighted with specific colours. The representation of the zebrafish fin bones comes from Schneider and Shubin ((115); *Danio rerio*). The representation of the appendicular appendage in *Eusthenopteron foordi*, *Panderichthys* sp., *Acanthostega gunnari* and *Tulerpeton* sp. are modified after Clack (116). The representation of the mouse limb bones is modified after Onimaru *et al.* ((117); *Mus musculus*).

Surprisingly, none of them has considered the microanatomical architecture of the pectoral elements of stem tetrapods in their simulations although they are known to be very spongy. Microanatomical investigations conducted by several authors (6,11,12) concluded that the great sponginess of these elements was probably mechanically too weak to enable body weight support on land and this became commonly accepted (e.g., (5)).

Faced with these contradictions between morphological and microanatomical data, we propose here to test the biomechanical resistance of the trabecular mesh of a stem tetrapod to terrestrial conditions. Even though we know that *Eusthenopteron* was a fully aquatic stem tetrapod fish, we have decided to focus our study on this taxon as it has an exquisite 3D-preserved fin skeleton with intact trabeculae (13–15). *Eusthenopteron* belongs to the sister group of tetrapods, the tristichopterids. Its pectoral fin comprises a well differenciated humerus with a microanatomy similar to that of limbed stem tetrapods (e.g., (16,17)), i.e. it is a dense trabecular mesh greatly extended and surrounded by a thin cortex (13–15). It has a strong longitudinal directionality and it comprises tubular structures called marrow processes, involved in the elongation of the bone (15,18). The humerus has played a crucial role in the tetrapod land exploration (e.g., (1,19,20)). As the skeletal element connecting the shoulder girdle to the pectoral appendage, it distributes and absorbs many various stresses (3,21) and therefore is most likely a great fin/limb bone to highlight different stress patterns over the water-to-land transition. For this reason, we have focused our investigation on this element.

## MATERIALS AND METHODS

### Material

*Eusthenopteron foordi* is a sarcopterygian fish from the Frasnian (Late Devonian, i.e. 380 million years ago). The current study focuses on the humerus of an articulated adult specimen (NRM P248a) from the Escuminac Formation, Miguasha, Canada (22). The specimen is housed in the collections of the Naturhistorika Riksmuseet (NRM), Stockholm, Sweden (Fig 2).

**Fig 2.**
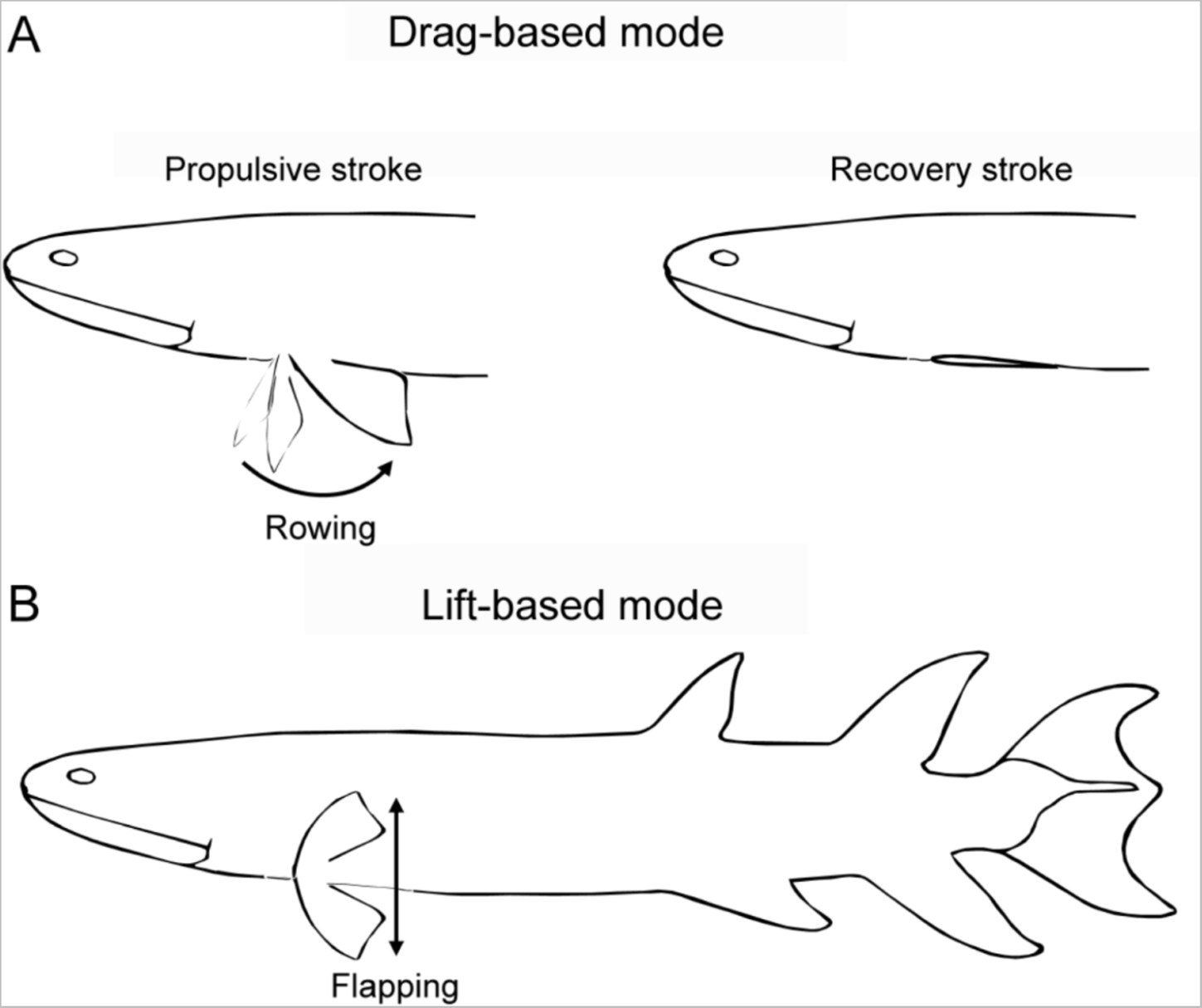
Representation of *Eusthenopteron foordi*. (A) Body reconstruction of *E. foordi*. The humerus is highlighted in red; (B) 3D rendering of the humerus (NRM P248a) with anatomical orientation axes (x, y, z). Note that the dorsal and ventral orientations are specific to either the animal or the bone.

### Method

#### 1. Characterisation of the forces applied on the pectoral fin of *Eusthenopteron foordi*

We want to test the resistance of the humerus of *Eusthenopteron foordi* to forces applied a) while swimming and b) in virtual standing terrestrial conditions. To do so, we need to identify the intensity of these forces before performing a biomechanical simulation using Finite-Element Analyses (FEA).

##### Swimming condition

*Swimming analog to* Eusthenopteron foordi *among extant actinopterygians*

###### Northern pike

We refer to the extant northern pike (*Esox lucius*) as a functional model for *E. foordi.* Both species have similar morphologies, characterised by a slender body shape and a posterior position of the dorsal and anal fins, able to give powerful thrusts when needed. *E. lucius* is an ambush predator that captures preys thanks to a fast-start strategy, which is defined by high-powered bursts of acceleration from a resting position (23). Based on morphological similarities, *E. foordi*’s lifestyle and behavior have been assumed to be close to those of *E. lucius* (24–27). *E. lucius* has been classified as a Body and Caudal Fin (BCF) transient swimmer, *i.e.* a swimming style that is characterised by fast-starts and powerful turns (28,29) during which the pectoral fins are held close to the body for hydrodynamic purposes (30). However, when they swim at low speed or when they hold position in one spot, the BCF transient swimmers turn out to perform a Median and Paired Fin locomotion (MPF), *i.e.* a behavior relying on the use of the pectoral fins to maneuver and stabilise the body at one spot while being exposed to the water flow (30,31). We expect this behavior to sum up the main uses of *E. foordi*’s pectoral fins in a near-shore complex turbid environment (22).

###### Catfish

*P. hemioliopterus* is a catfish that lives in turbid shallow waters (32) while showing a lower body length-to-weight ratio than the northern pike (*Esox Lucius*, (33)), which would result in increasing the ground reaction force at the elbow joint.

*Calculation of forces applied onto the humerus of* Eusthenopteron foordi *in labriform swimming mode*

In order to estimate the forces applied on the pectoral fin of *E. foordi* when performing a MPF locomotion, we chose to rely on the forces that were measured with a robotic model of a fish pectoral fin (34). This model consists of a three-motor driven pectoral fin that simulates a simplified version of the labriform swimming mode by resuming the three basic movements of the pectoral fin: 1) rowing and 2) feathering during the drag-based mode; 3) flapping during the lift-based mode. Since the arm of the robotic fin integrates a force sensor, it provides a good proxy to estimate the resultant of the environmental forces that were transmitted to the humerus of *E. foordi* within a similar set of movements (34), *i.e.* during the recovery stroke of the drag-based mode (Fig 3) or at any time when the fin is at rest in a horizontal position.

**Fig 3.**
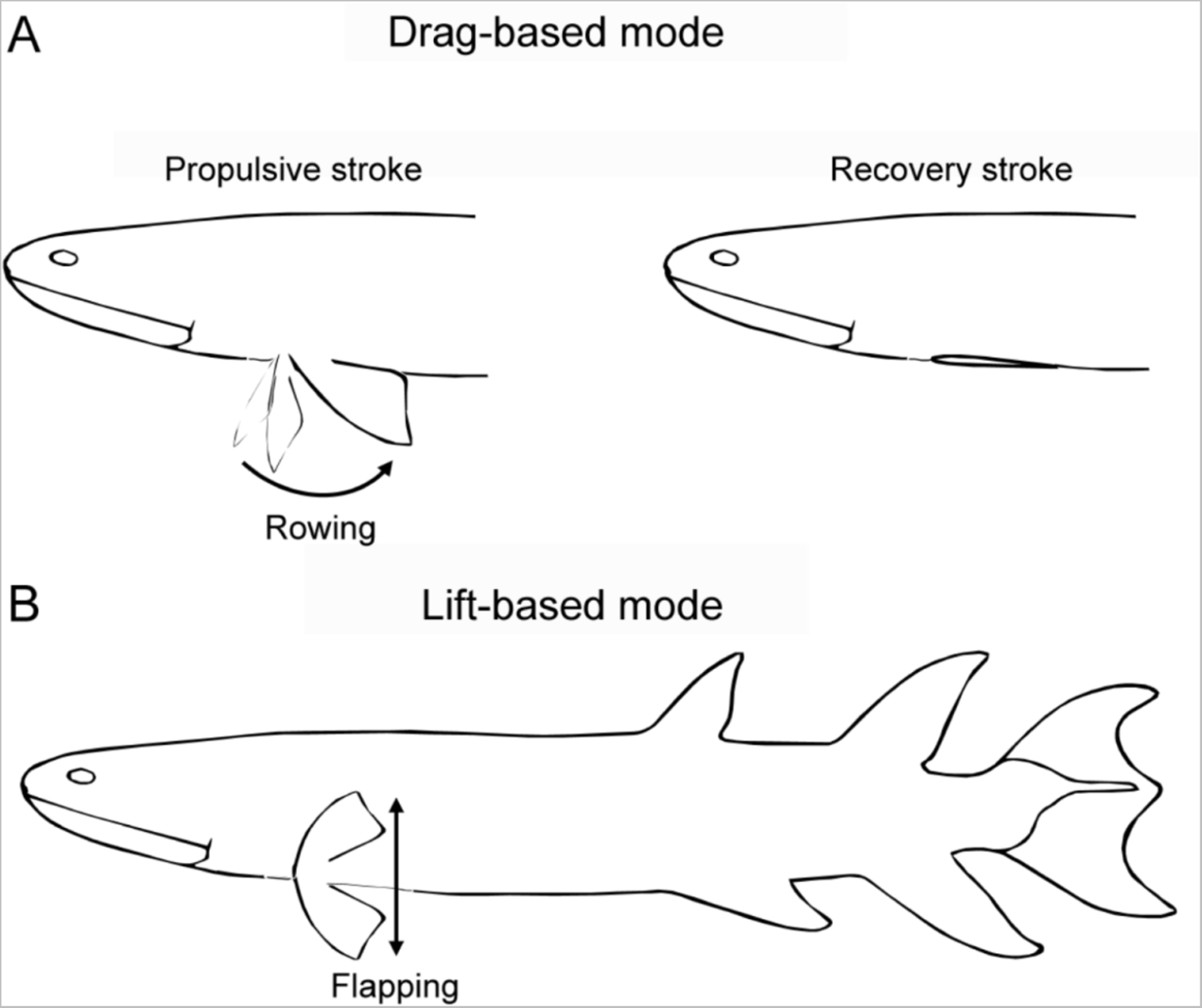
Labriform swimming mode. (A) Drag-based mode, showing both the rowing motion propulsive stroke, as well as the aligned recovery stroke; (B) Lift-based mode.

The sum of the external forces 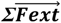 applying on the pectoral fin results from: 1) the viscous drag force 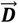 that is due to the resistance of the fin to the water flow; 2) the weight 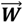 and the “resting” lift 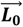 which compensate each other; 3) either a drag or a lift traction (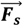; thrust) depending on the movement and orientation of the fin (Fig 4):

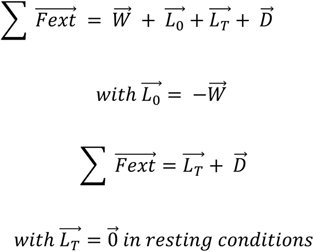

**Fig 4.**
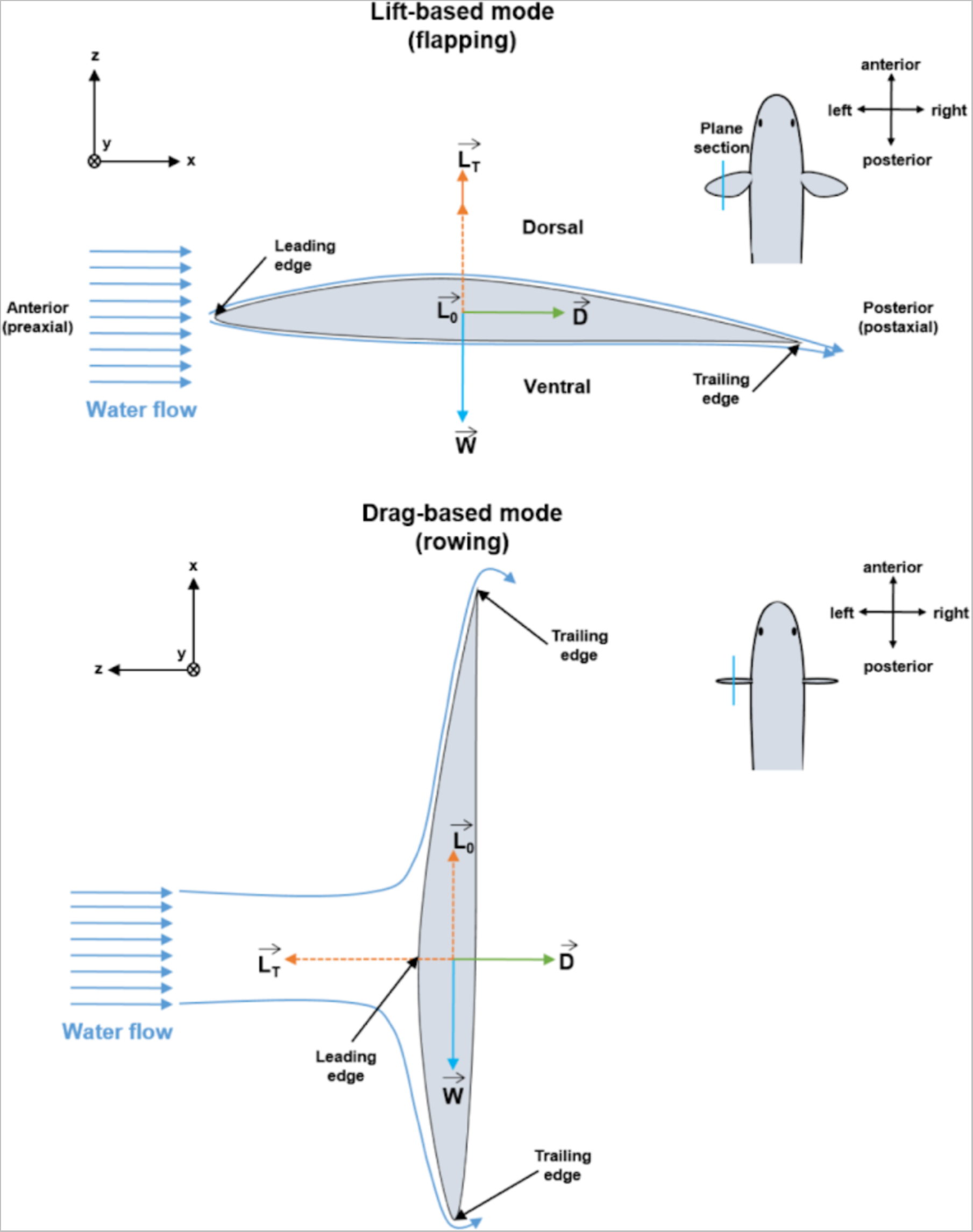
Resultant of the external forces that are applied on a pectoral fin during the labriform swimming. (A) Lift-based mode; (B) Drag-based mode; 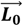 is the component of the lift that compensates the weight 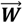; 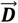 is the viscous drag forces of the water flow and 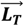 is the component of the lift that produces the thrust during the motion of the fin. The (x, y, z) axes represent the orientation of the humerus of *E. foordi* in this study.

We introduce 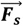 as the resultant of the external forces 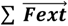 during swimming:

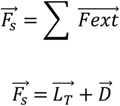

*Calculation of the drag and lift forces (**D** and **L_T_**)*

The swimming viscous drag forces ***D*** and lift forces ***L_T_*** that apply on an immerged body can be calculated by the following Newtonian equations:

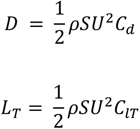

where ***ρ*** is the water density, ***S*** is the wetted surface area of the body, ***U*** is its velocity relative to the fluid (water), ***C_d_*** is the drag coefficient, and ***C_lT_***, the lift coefficient. These two coefficients are dimensionless quantities which are dependent on: the geometry of the object, its angle of attack and its surface texture (*e.g.* a scaled surface).

###### Values of the drag and lift coefficients (***C_d_*** and ***C_lT_***)

We assume that both the pectoral fin of the robot of (34) and the pectoral fin of *E. foordi* have the same drag coefficients (respectively *C_dr_* and *C_de_*) and lift coefficients (respectively *C_lTr_* and *C_lTe_*) because they share: 1) the same overall morphology (*i.e.* a streamline body, (35); 2) the same variation of both shape and angle of attack during the labriform swimming movements. These two parameters should vary similarly both along the passive flexible deformation of the rubber fin in the robot (34) and the coordinated movement of the fin rays in *E. foordi* (36).

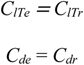

###### Value of the water density (***ρ***)

We consider that the water density ***ρ*** is a constant value. It only varies within a range of 3% in the current nature (*i.e.* from 1000 g/m^3^ in fresh waters up to 1030 g/m^3^ in the most salted and cold oceans;(37)). During the Devonian, the mean salinity of the ocean reached about 40 ‰ (38). As we consider that the sea surface temperature variations fluctuated between 20° and 30°C at the Miguasha locality during the Late Devonian (39), we then deduce that the resulting sea water density must have scored between 1025 and 1031 kg/m^3^. Since the Late Devonian Escuminac formation has been estimated to host an estuarine ecosystem (40), we therefore estimate that the water density must have remained below 1031 kg/m^3^ (*i.e.* at most 3% higher than the freshwater density).

###### Value of the pectoral fin surface of the robot (***S_r_***) and *Eusthenopteron foordi* (***S_e_***)

The variation in ***F_s_*** between the robot model ***F_Sr_*** and *E. foordi **F******_Se_*** only depends on the fin area ***S,*** and the velocity ***U*** that is relative to the water flow.

The surface area of the robot fin ***S_r_*** is provided by Suzuki *et al*. (34):

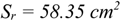

Consequently, we need to obtain a value for the surface area of the pectoral fin of *E. foordi **S******_e_***. The full pectoral fin is not preserved around the (left) humerus of the specimen which we study here (NRM P248a); nevertheless, all the axial bones are preserved along the right fin within the same specimen (NRM P248d and NRM P248b; Fig 5). We therefore used the right pectoral fin to calculate the specimen pectoral fin surface ***S_e_***. We have treated a photograph of a fully preserved pectoral fin of *E. foordi* (GN.787 UMZC from the University Museum of Zoology, Cambridge; Fig 3.18 in (27); Fig 5) using ImageJ (IJ 1.46r: Wayne Rasband, National Institute of Mental Health, Bethesda, Maryland, USA). We modelled the surface of the pectoral fin as an ellipse that is defined by a semi-major axis (***l***, *i.e.* the length from the proximal part of the humerus to the distal end of the preaxial radial III) and a semi-minor axis (***l_m_***, *i.e.* the transversal radius of the fin).

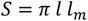

**Fig 5.**
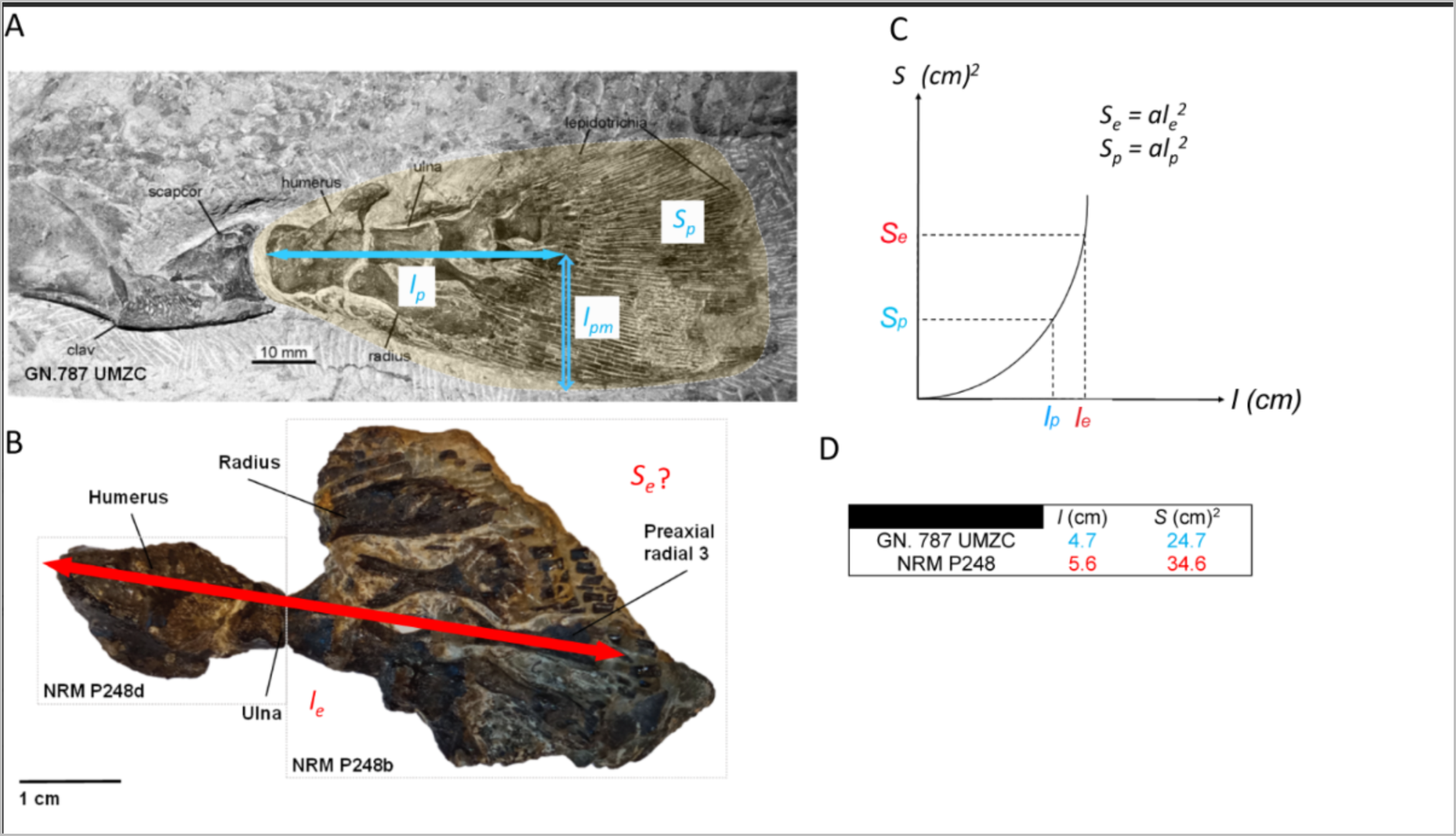
Estimation of the pectoral fin area in *Eusthenopteron foordi*. (A) Right pectoral fin of the GN.787 UMZC specimen: the surface area is ***S_p_*** plotted in shaded yellow and delimited with a white dashed outline. The picture is modified after Clack 2012 ((27); fig. 3,18). ***l_p_*** is the semi-major axis of the pectoral fin, considered as an ellipse. ***l_p_*** extends from the proximal part of the humerus to the distal end of the preaxial radial III; (B) Right pectoral fin of the NRM P248 specimen: the proximal half of the fin (NRM P248d) and the distal elements (NRM P248b) are connected along the red arrow which indicates the length ***l_e_*** of the semi-major axis of the pectoral fin; (C) Square function relation between the surface area (***S_p_***, GN.787 UMZC; ***S_e_***, NRM P248a) and the length (***l_p_*** GN.787 UMZC, ***l_e_*** NRM P248a) of the pectoral fin of *E. foordi*; (D) Calculation of the pectoral fin area ***S_e_*** in the specimen of interest based on the square function *S = al^2^*.

The specimen of reference (GN.787 UMZC) has therefore a surface ***S_p_*** defined as follows:

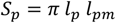

The specimen of interest (NRM P248a) has a surface ***S_e_*** defined as follows:

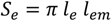

We consider that the axis length-ratio 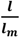 is equal between both specimens (NRM P248a and GN.787 UMZC). Indeed, they are both adults (15,27) and should therefore present no morphological difference in term of allometric growth. We consequently define a proportional relationship between ***l*** and ***l_m_*** that is identical to both specimens.

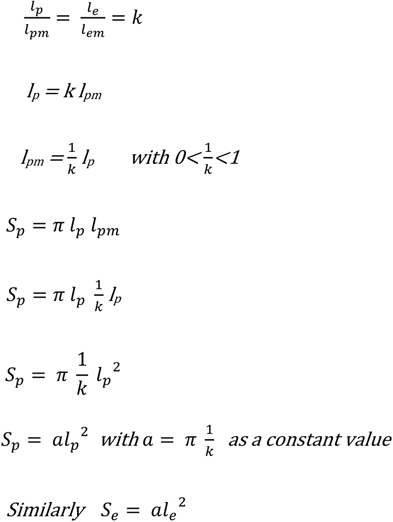

The length ***l_p_*** measures 4.7 cm. The surface ***S_p_*** was therefore calculated to 24.7 cm^2^. The length ***l_e_*** was measured to 5.6 cm (Fig 5B) on the right pectoral fin of NRM P248 (NRM P248d and NRM P248b; Fig 5B).

Based on this, we could calculate the surface area of the right pectoral fin of NRM P248 and therefore deduce the surface of the left fin NRM P248a (***S_e_***; Fig 5D):

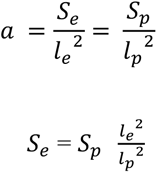

We found a surface ***S_e_*** of 34.6 cm^2^.

###### Value of the velocity of the robot (***U_r_***) and *Eusthenopteron foordi* (***U_e_***)

The velocity of the robot fin ***U_r_***, relative to the surrounding aquatic environment, is given by Suzuki *et al*. (34):

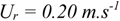

We have estimated that the mean velocity of *E. foordi **U******_e_*** was probably similar to that of the Northern pike, *Esox lucius*, whose Body Length by second was measured to 0.45BL/s (41). This measurement translates into a mean relative velocity:

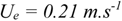

This assessment relies on the body length of the studied specimen of *E. foordi*. The fin NRM P248a is associated in the collection with other skeletal elements including an 89-mm long lower jaw, which is representative of the standard body length. Thomson and Hahn (42) demonstrated a linear relationship between the length of the lower jaw and the overall standard body length. The standard length extends from the tip of the snout to the posterior end of the most caudal vertebra (42). Because the vertebral column almost reaches the tip of the caudal fin, we can here consider that the standard length greatly represents the full body length. *E. foordi*’s standard body length is estimated to reach 478 mm.

*Resultant between the external forces of* Eusthenopteron foordi *and the robot during labriform swimming*

Finally, we can calculate the resultant of the external forces applied on the humerus of *E. foordi **Fs******_e_*** during labriform swimming, based on the drag and lift equations mentioned above, the calculated values of ***S_e_*** (34.6 cm^2^) and ***U_e_*** (0.21 m.s^-1^) and the robot pectoral fin characteristics that were designed by Suzuki *et al*. 2008 (34) (*i.e. **S******_r_*** = 58.35 cm^2^; ***U_r_*** = 0.20 m.s^-1^):

To simplify the following equation, let’s introduce *C_Tot_* as the resultant of the lift coefficient (*C_lt_*) and the drag coefficient (*C_d_*), considered equal in both models at any moment during labriform swimming (cf. values of the drag and lift coefficients mentioned above).

The ratio 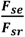 can then be summed up as the coefficient of variation ***K_var_*** in the resultant of the external forces:

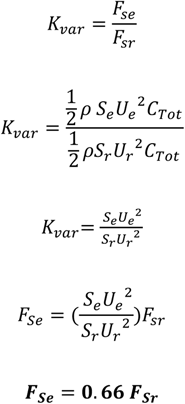

*Values of the external forces applied on the humerus of* Eusthenopteron foordi *during labriform swimming*

The hydrodynamic loads shown by Suzuki *et al*. (34) represent the resultant of the external hydrodynamic forces which are applied on the robot pectoral fin during the drag-based mode and the lift-based mode (fig. 16a & 16b respectively in Suzuki *et al*. (34)). The loads are represented by two curves displaying the variation of the force intensity in each direction of the 3D space (x, y, z) within each type of movement (drag-based and lift-based). In order to assess the greatest constraints the humerus can endure, we have first selected the highest value in whichever axis and then picked up the corresponding values in the other axes to set loads on the Finite-Element (FE) model (see details below). Eventually we inverted the coordinate system used by Suzuki *et al*. (see Fig. 4 in (34)) to suit to the anatomical coordinated system we have displayed in Fig 2. The values for ***F_se_*** and ***F_sr_*** are given in table 1.

**Table 1.** Summary of labriform swimming forces, drag-based and lift-based respectively, applied at the elbow joint of *Eusthenopteron foordi*, along each anatomical axis of the pectoral fin (x,y,z), *i.e*. x: preaxial to postaxial; y: distal to proximal; z: ventral to dorsal. *F_sr_* is the force computed by the arm of the robot pectoral fin (34). *F_se_* is the force that we will apply at the elbow joint of *E. foordi*’s humerus, deduced from the relation *F_se_ = 0.66 F_sr_*

|  |  | Drag-based mode (N) | Lift-based mode (N) |
| --- | --- | --- | --- |
| **Robot model ($F_{sr}$)** | -x | 4.2 | 1.3 |
|  | -y | 1.1 | -0.8 |
|  | -z | 0.1 | 2.4 |
| ***E. foordi* model ($F_{se}$)** | x | -2.77 | -0.86 |
|  | y | -0.73 | 0.53 |
|  | z | -0.07 | -1.58 |

##### Testing a hypothetical terrestrial model

We want to check the resistance of the humerus of *Eusthenopteron foordi* to terrestrial stresses to understand when and how this resistance to land constraints evolved over the fin-to-limb transition. We want to rule out here any assumption that we are testing *E. foordi*’s ability to walk on the ground.

*Characterisation of the ground reaction force*

To test whether the microarchitecture and morphology of the humerus of *E. foordi* would have been able to support the body weight on land, we have assessed the ground reaction force according to Newton’s third law of motion (43,44), i.e. the ground reaction force 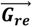 results from a mutual and simultaneous interaction between the animal and the ground. In standing position, 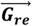 corresponds to the animal weight:

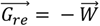

where 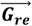 stands for the total ground reaction force that applies on the specimen in response to the full body weight 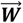.

We have assessed the ground reaction force transmitted to the humerus 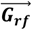 as if the fish was standing on the four paired fins on land. In that situation, we assume that the specimen weight is equally distributed on the four paired fins.

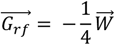

where 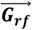 stands for the ground reaction force component that applies on one pectoral fin.

Here we suppose that the ground reaction force applied on the pectoral fin should be fully transmitted to the humerus via the distal-proximal axis of the pectoral fin. This scheme is biased and leads to extrapolate the force transmission to the elbow joint. In reality, it would be partially absorbed and compensated by the distal elements of the fins (made of bone, cartilage and muscles). However, this systematic error is not detrimental for the current study because we do not aim at testing whether *E. foordi* could walk on land but we rather aim at testing the resistance of the humerus of *E. foordi* to high terrestrial stresses. We thus propose the following equations to build up our Finite-Element (FE) model:

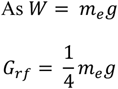

***m_e_*** is the full body mass and 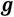 is the gravitational force equivalent (acceleration).

###### Value of the mass

We have estimated the mass of the studied specimen of *E. foordi **m******_e_*** based on the body length-to-mass relation of the Northern pike established by Willis (33). Following their proposed equation, we have:

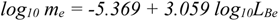

where ***m_e_*** is the mass of *E. foordi* in grams and ***L_Be_*** is the standard body length of *E. foordi* in millimetres.

As estimated above, based on the lower jaw/body standard length relationship established by Thomson and Hahn (42):

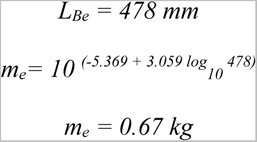

###### Value of the gravitational force equivalent (acceleration)

The paleolatitude of Miguasha during the early part of the Late Devonian was at θ = 17° N (45) based on the international gravity formula (Geodetic Reference System 1967):

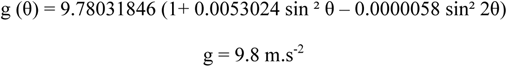

*Calculation of the ground reaction force*

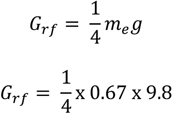

The ground reaction force applied on each paired fin of *E. foordi **G_rf_*** therefore scores 1.64 N.

Since this calculation is very dependent on the mass of the specimen, we decided to repeat the analysis for the analogy with the catfish, *Phractocephalus hemioliopterus* (32). This species shows a lower body length-to-mass ratio (especially in female which are significantly larger than males; (32)). We used the following equation to determine the mass of *E. foordi* as if it had the same body length-to-weight as a female *P. hemioliopterus* specimen (32):

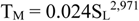

where (**T_M_**) is the total mass in grams and (**S_L_**) is the standard length in centimetres. We thus determined a body mass of 2.35 kg and a consequent weight of 23.03 N for *E. foordi*. This last value represents an increase by a factor of 3.5 in comparison with the scenario in which we used northern pike body length-to-weight ratio. In this new configuration, the ground reaction force that is applied on one pectoral fin reaches 5.75 N (table 2).

**Table 2.**
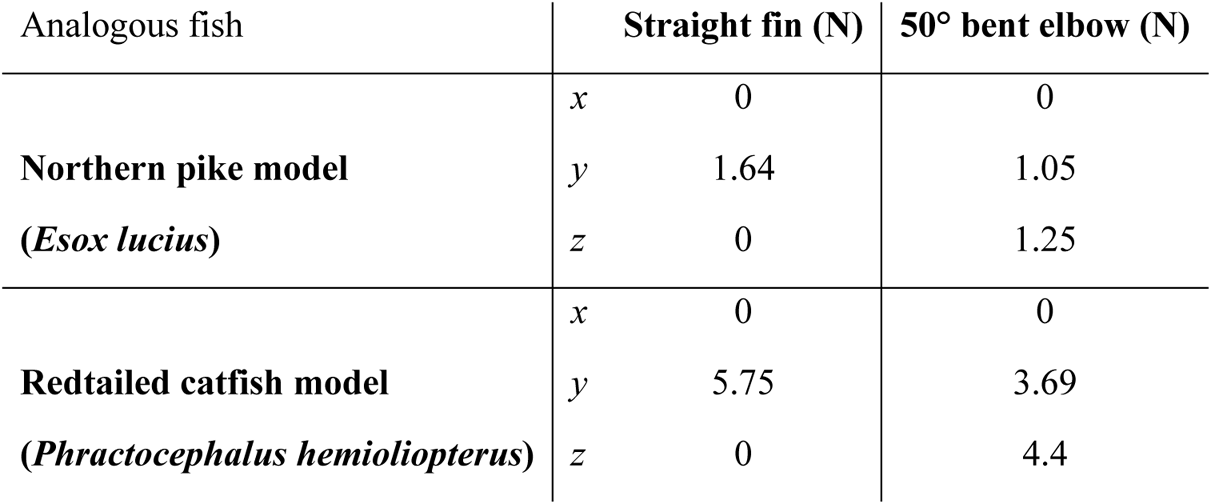
The Ground reaction forces (G_rf_) applied to the elbow joint of *Eusthenopteron foordi* in the simulated terrestrial models, straight-finned and elbow bent at 50° respectively, along each anatomical axis of the pectoral fin (x, y, z), *i.e.* x: preaxial to postaxial; y: distal to proximal; z ventral to dorsal.

| Analogous fish |  | Straight fin (N) | 50° bent elbow (N) |
| --- | --- | --- | --- |
| **Northern pike model**<br>**(*Esox lucius*)** | x | 0 | 0 |
|  | y | 1.64 | 1.05 |
|  | z | 0 | 1.25 |
| **Redtailed catfish model**<br>**(*Phractocephalus hemioliopterus*)** | x | 0 | 0 |
|  | y | 5.75 | 3.69 |
|  | z | 0 | 4.4 |

#### 2) Finite Element Analysis (FEA)

The aim of the study is to provide both quantitative results and graphic representations of mechanical stresses supported by the humerus of *Eusthenopteron foordi* under the different loading situations characterised above. Three models will be produced:

- a “plain” model. In this case, the inner microanatomy of the humerus is not treated but instead the resistance of the overall bone is based on a theoretical cortical bone value.
- an “isotropic cancellous” model, commonly used in FEA, whose resistance value is based on a mechanical test of a referenced cancellous bone. Here we have taken a value in the literature reflecting a very sparse organization of the spongiosa in the humerus of *E. foordi* (Young’s modulus of 13.5 GPa; (46)). In this configuration, the trabecular organization is considered isotropic (i.e. with no preferential orientation).
- an “actual trabecular” model, for which the trabecular arrangement in the medullary cavity has been modeled and meshed in 3D.

Comparison between the different models will highlight the role the orientation of the trabecular mesh plays in absorbing stresses during swimming or while standing. We will refer to the von Mises criterion to solve the Finite-Element (FE) models and predict their resistance to environmental stresses. This von Mises criterion is commonly used to define a strength criterion in terms of maximum distortion of a ductile material (yield strength; (47)).

##### Scanning parameters

The humerus of *Eusthenopteron foordi* (NRM P248a) was imaged at the European Synchrotron Radiation Facility (ESRF, France) using Propagation Phase-Contrast Synchrotron Radiation Microtomography (PPC-SRµCT) at beamline ID19, with a voxel size of 20.24 µm. The experiment was done in monochromatic conditions, using a double Si111 Bragg monochromator and a FreLON 2k14 CCD detector (48). The optics system was coupled to a 20 μm-thick Gadox scintillator. The propagation distance – between the sample and detector – was fixed to 950 mm. The scans were performed with an energy of 60 keV.

##### Data reconstruction and segmentation

The raw data were filtered and converted into TIF images at the ESRF. The segmentation of the humerus NRM P248a was made with the software VGStudio MAX version 2.1 (Volume Graphics Inc., Germany) at Uppsala University (Sweden).

##### Creation of 2D triangular surface FE meshes

The first step of FEA is to create a FE mesh out of a 3D segmented model (Fig 6A). To do so, we use the “opening-closing” tool in VGStudio MAX in order to clean and minimise surface connection problems creating small and sharp artefacts. We produce a surface mesh englobing the entire bone; this mesh is composed of 16 million triangles that we export as a STL file. To homogenise and reduce the number of triangles, we use Geomagic wrap 2017 (3D systems Inc.; Fig 6B), and more specifically the automated repairing tool “mesh doctor” to remove the mesh self-intersections. We then use the “quick smooth” tool to homogenise the size and the shape of the triangles. The latter stage consists in applying a filter to smoother the mesh surface by averaging the position of the triangles and removing sharp edges. At this stage, the remaining kinks can be manually corrected thanks to a full set of dedicated tools in Geomagic. After this first remeshing process, we obtain a mesh composed of 12 million triangles. Due to software and workstation calculation limitations, we need to reduce this number to 9 million triangles to be able to run a FEA on a 3D volume mesh model. By increasing the edge length of the triangles, via the “remesh” tool, while keeping the self-intersection removing process on, we obtain a final 2D triangular surface mesh, composed of about 8.7 million triangles. The “plain” and “isotropic cancellous” models are created from a 2D surface mesh by deleting the microanatomical structures inside the bone (i.e. the trabeculae). We thus built two models:

- one 2D triangular surface mesh that reconstructs both the external morphology of the bone and the inner trabeculae (for the “actual trabecular” configuration);
- one model that consists of an empty external envelop that represents the bone external shape only (for the “plain” and “isotropic cancellous” configurations).

**Figure 6.**
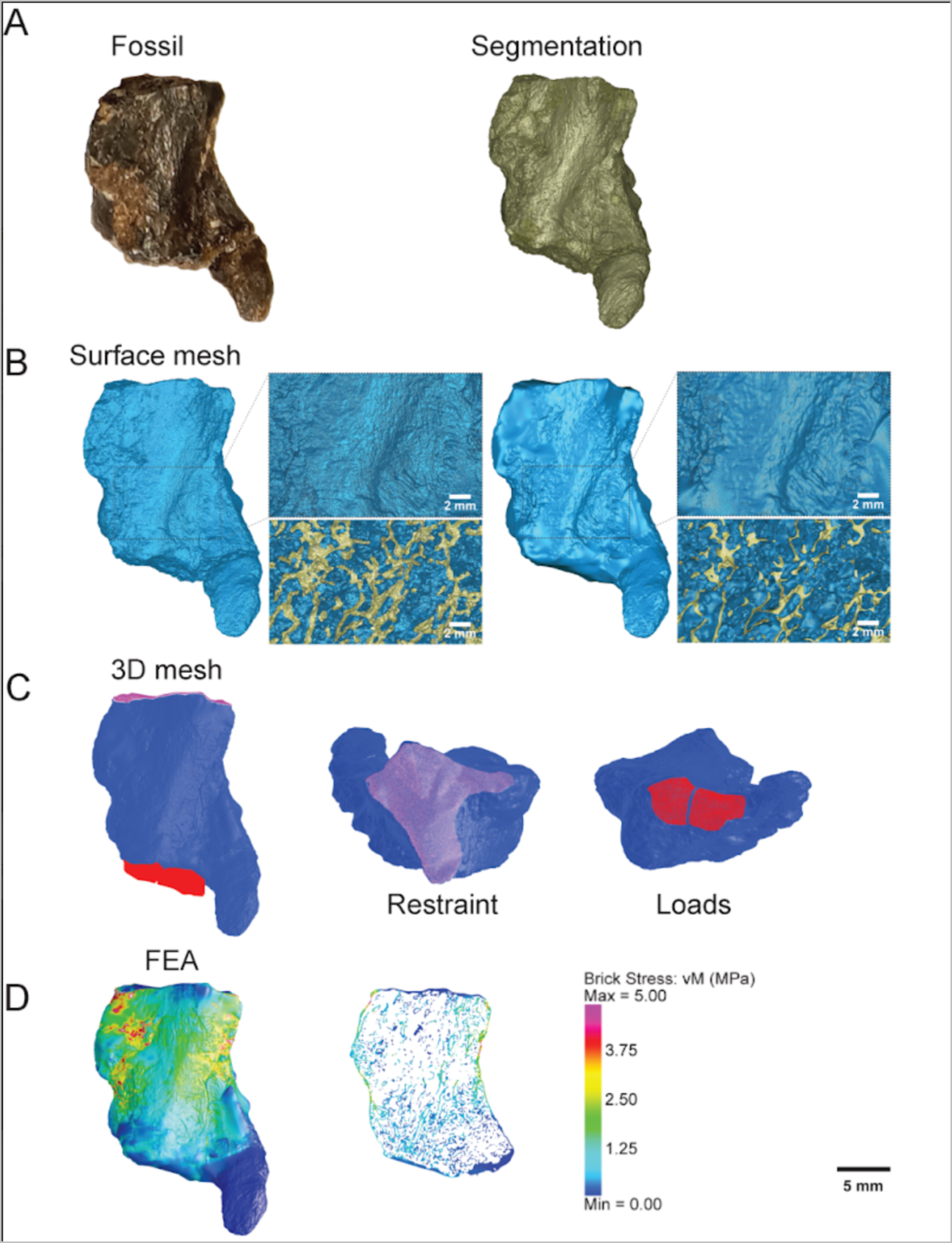
Summary of the finite element processing on the humerus of *Eusthenopteron foordi*. (A) synchrotron scan and segmentation of the specimen; on the left: a picture of the actual fossil; on the right: the 3D-segmented model; (B) On the left: the original 2D-triangular mesh; on the right: the obtained 2D-triangular mesh after the full remeshing process; (C) Selection of the regions of interest and attribution of the properties at the nodes; restraining at the shoulder articulation, loading at the ulnar and radial articulations (D) Finite Element Analysis (FEA), from left to right: stresses shown on the surface model, a thin section showing the stresses on the trabecular mesh.

##### Generation of 3D volume mesh models

We import both meshes in 3matic 13.0 (Materialize inc., Belgium) to produce the volume mesh, *i.e.* to transform the 2D surface triangles into a 3D volume mesh of tetrahedrons. To do so, we use the “create volume mesh” tool. The plain model finally consists of 9 133 335 tetrahedrons and the trabecular model of 12 804 546 tetrahedrons. We then need to check the quality of the surface mesh of both 3D volume models again and clean it, if necessary, using the “fix wizards” tool. The remaining “shells” (*i.e.* non-connected elements) and overlapping triangles can now be completely removed.

##### Finite Element Analyses (FEA)

We export both models as NASTRAN files in order to import them in Strand7 3.0.0 Preview 37 (Pty Ltd., Australia) (Fig 6C) to perform the FEA. We first set the material properties to:

- an elasticity of the bone in tension or compression (measured by the parameter called Younǵs modulus):

`o` of 18 GPa for the “plain” and “actual trabecular” models (Strand 7 default value for the compact bone);
`o` of 13.5 GPa for the “isotropic cancellous” model (46);
- a Poissońs ratio of 0.3. This is a value that reflects the expansion or contraction of a material in the directions perpendicular to the load direction (49);
- a bone density of 2 g/cm^3^ (50).
We then select the bone external area connecting to either the elbow or the shoulder joint by manually selecting the nodes on the surface of the mesh to fix the articulations. According to Andrews and Westoll (24), the shoulder joint of *E. foordi* was immobile besides one possible rotating upward movement with an angle of 20° (along the *x* axis). This has been taken into consideration when restraining the nodes at the shoulder joint of the humerus NRM P248a (Fig 6C). Regarding the elbow joint, we have selected the nodes in two separate areas interpreted as the regions of contact between the humerus and the radius or ulna (through cartilage). We then apply the previously calculated forces on the nodes of these selected areas in the framework of a homogenous repartition. We finally run the analyses (Fig 6D) using the “solver” tool (window). We have selected the following parameters:
- a Linear Static Analysis (LSA, which involves a small deformation with no change of loading with time);
- a sparse storage scheme;
- an AMD (Approximate Minimum Degree ordering algorithm) sorting method;
- an Iterative Preconditioned Conjugate Gradient (PCG) solution type. The number of iterations has been set to 10,000 for the plain model, and to 50,000 for the trabecular model in order to ensure the convergence of the solution for each model;
- the convergence tolerance to 1.0 x 10^-5^.

We export a table gathering all the calculated mean stress values for each finite single element (tetrahedron) in a .txt format (*i.e.* n_t_ = 12 804 546 for the “actual trabecular” model; n_p_ = 9 133 335 for the “plain” and “isotropic cancellous” models). We open this file with R 4.0.0 (CRAN; R Core team) to run a non-parametric statistical test (Wilcoxon signed-rank test as n_t_ and n_p_ are not distributed normally) to compare the median stress between these two models. This comparison is crucial for highlighting the role of the trabecular orientation.

##### Simulations

###### Swimming

Values are taken from the pike’s (*Esox lucius*) pectoral fin and body properties for the swimming simulation in both drag-based and lift-based modes (Fig 7).

**Figure 7.**
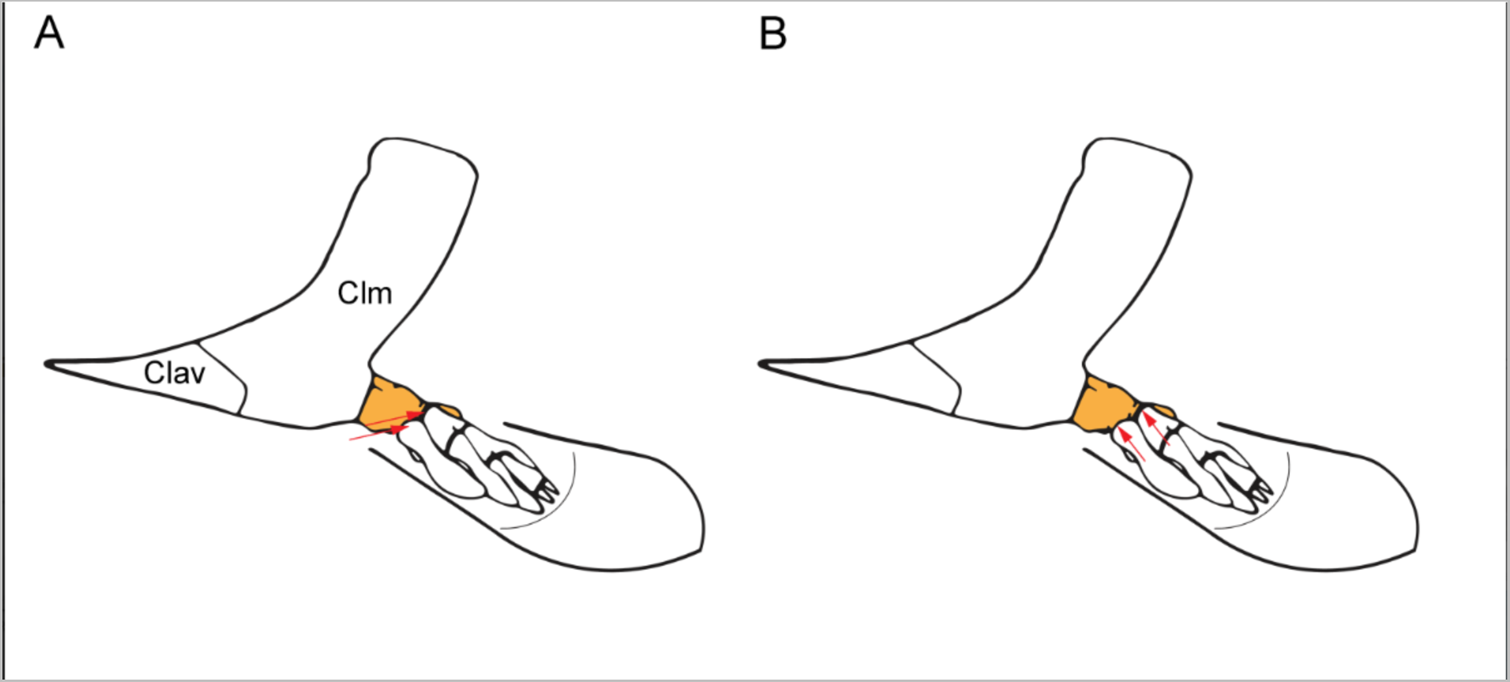
Application of the swimming stress forces on the elbow joint of the humerus in *Eusthenopteron foordi* in labriform swimming models. (A) Lift-based mode with the main direction of loading on the leading edge of the fin due to water flow. (B) Drag-based mode with the main direction of loading from an alternate orientation due to motion and rotation of the fin. See Fig 4 for details. The humerus is highlighted in orange. Abbreviations: (Clv) clavicle; (Clm) cleithrum.

###### Terrestrial standing

In standing terrestrial conditions, values are taken from both the pike (*E. lucius*) and the catfish (*Phractocephalus hemioliopterus*). In order to test if the humerus of *E. foordi* was sensitive to the direction the ground force reaction transmitted to the elbow joint, we have considered the different configurations of the pectoral fin. Andrews and Westoll (24) suggested two potential configurations: either 1) the whole pectoral fin lays straight along the proximo-distal axis of the humerus (Fig 8A), or 2) the pectoral fin is bent down vertically by an angle of 50° at the elbow joint (Fig 8B). In the first case, 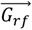 follows the *y* axis (Fig 8A); in the second case, 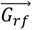 is decomposed into two components through the *y* axis and the *z* axis 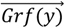 and 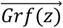 (Fig 8C). The forces are calculated

**Fig 8.**
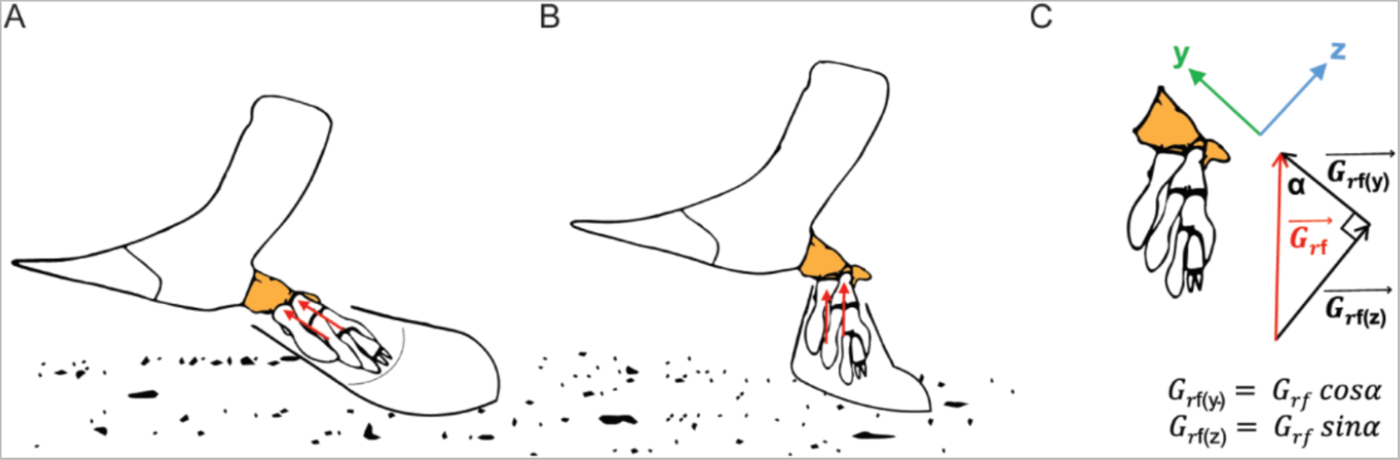
Application of the ground reaction force (G_rf_) on the elbow joint of the humerus in *Eusthenopteron foordi* in the simulated terrestrial models. (A) “straight fin” configuration: the whole pectoral fin lays straight along the proximal-distal axis of the humerus; (B) “50° bent elbow” configuration: the pectoral fin is bent down by an angle of 50° at the elbow joint; (C) Trigonometric method to calculate the components of the ground forces on proximal-to-distal axis (G_rf(y)_) and on the ventral-to-dorsal axis (G_rf(z)_) in the “50° bent elbow” configuration. The humerus is highlighted in orange. Abbreviations: (Clv) clavicle; (Clm) cleithrum.

(Table 2) according to the laws of trigonometry (Fig 8C) as follows:

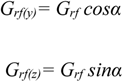

## RESULTS

Finite Element Analyses (FEA) have produced different patterns that we describe and compare here. We will consecutively focus on the data modelled a) while swimming and b) in virtual standing terrestrial conditions.

### Swimming models: qualitative comparisons

Figures 9 and 10 display mechanical stress distribution (von Mises criterion) in the humerus of *Eusthenopteron foordi* during two active phases of the labriform swimming (34): 1) the drag-based mode – referring to the horizontal paddling movement (Fig 9, Fig 3A); 2) the lift-based mode – referring to the vertical paddling movement (Fig 10, Fig 3B). Each simulation was conducted three times: 1) on the “plain model” (Young’s modulus of 18 GPa; Fig 9A and 10A); 2) on the “isotropic cancellous” model (Young’s modulus of 13.5 GPa; Fig 9B and 10B); and 3) on the “actual trabecular” model (based on the segmentation of the trabeculae considered with a Young’s modulus of 18 GPa; Fig 9C and 10C).

**Fig 9.**
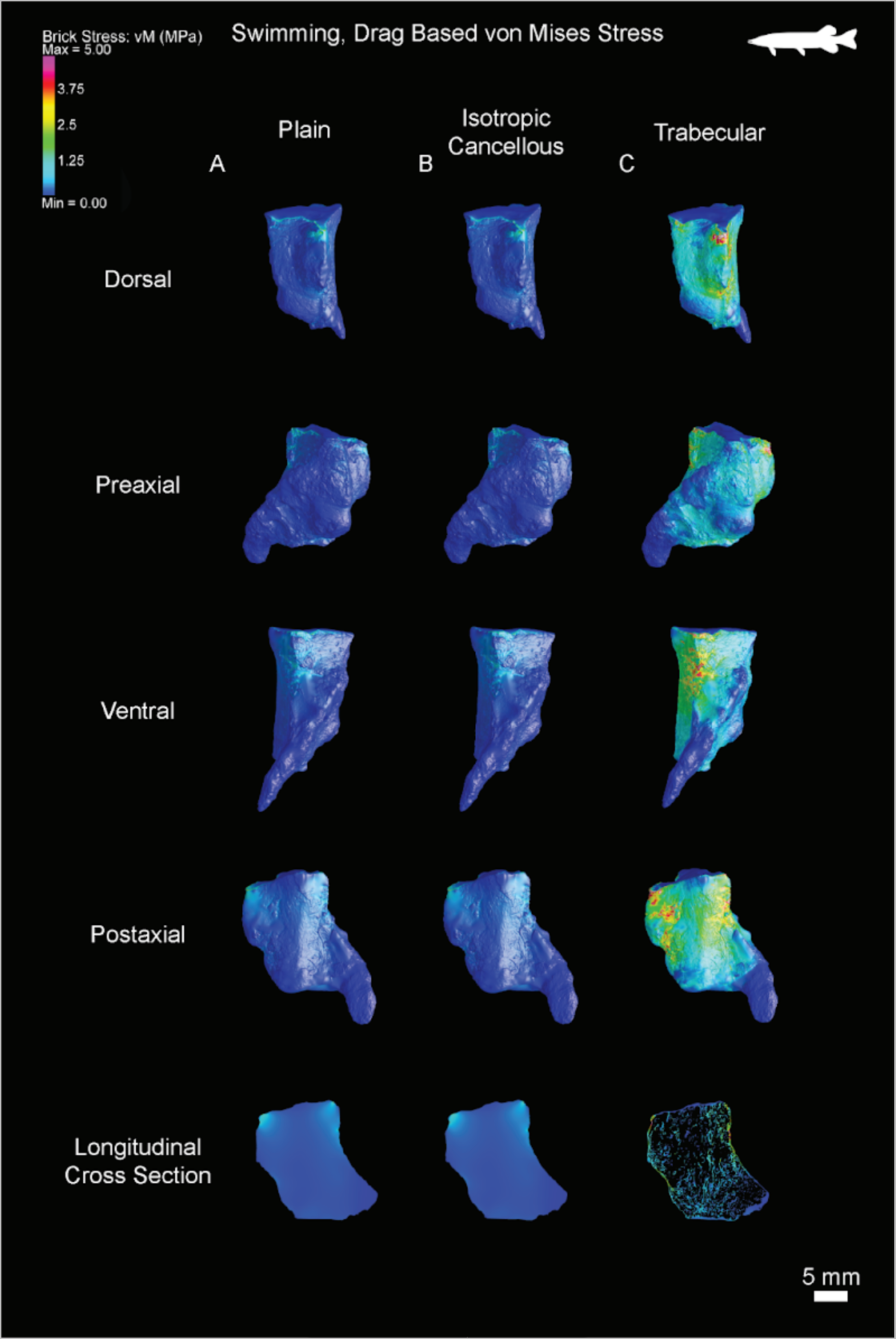
Distribution of the von Mises stress in the drag-based swimming model based on the pectoral fin properties of the northern pike (*Esox lucius*), applied to the *Eusthenopteron foordi* humerus; Stress distribution from 0-5 MPa. Four views and a longitudinal cross section are shown. (A) ‘’Plain’’ mode; (B) Isotropic cancellous model; (C) Trabecular mesh model.

**Fig 10.**
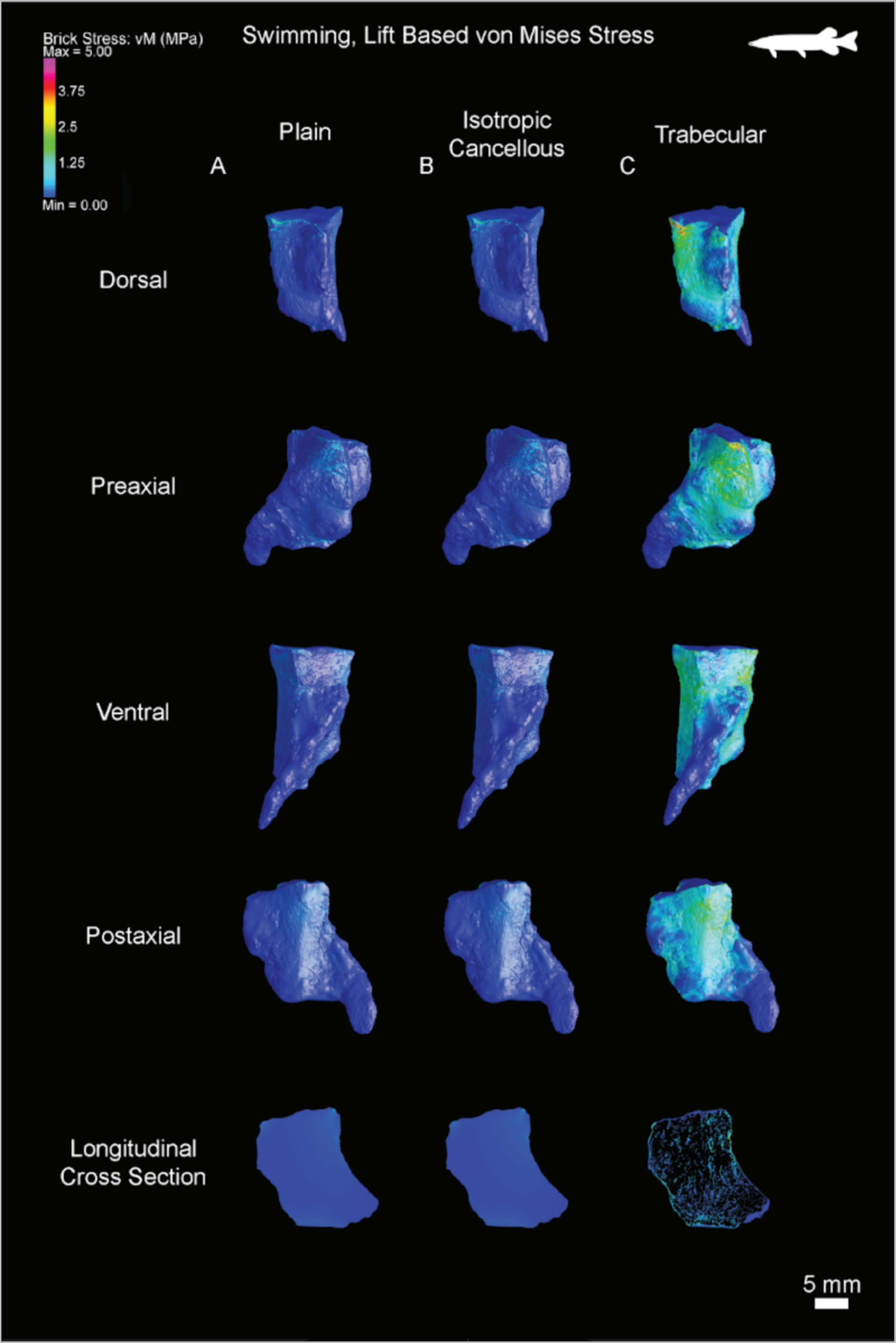
Distribution of the von Mises stress in the lift-based swimming model based on the pectoral fin properties of the northern pike (*Esox lucius*), applied to the *Eusthenopteron foordi* humerus; Stress distribution from 0-5 MPa. Four views and a longitudinal cross section are shown. (A) ‘’Plain’’ mode; (B) Isotropic cancellous model; (C) Trabecular mesh model.

#### 1) Drag-based mode (based on the pike’s body properties – *Esox lucius*)

The FEA results mostly reveal a “plain” model in dark blue (Fig 9A). This is the colour code for a stress value lower than 1MPa (Fig 9A). This demonstrates that the humerus in drag-based swimming conditions is not very stressed. A local stress can be visualised in light blue on the radial and ulnar facet of the distal articulations (in pre- and post-axial views on Fig 9A) but this should be considered artificial as due to the application of the ground reaction force at these points. Slightly higher stress values (of 1-2 MPa) can be visualised in light blue in the proximal region of the deltoid process and coracobrachialis fossa (Fig 11; Fig 9A).

**Fig 11.**
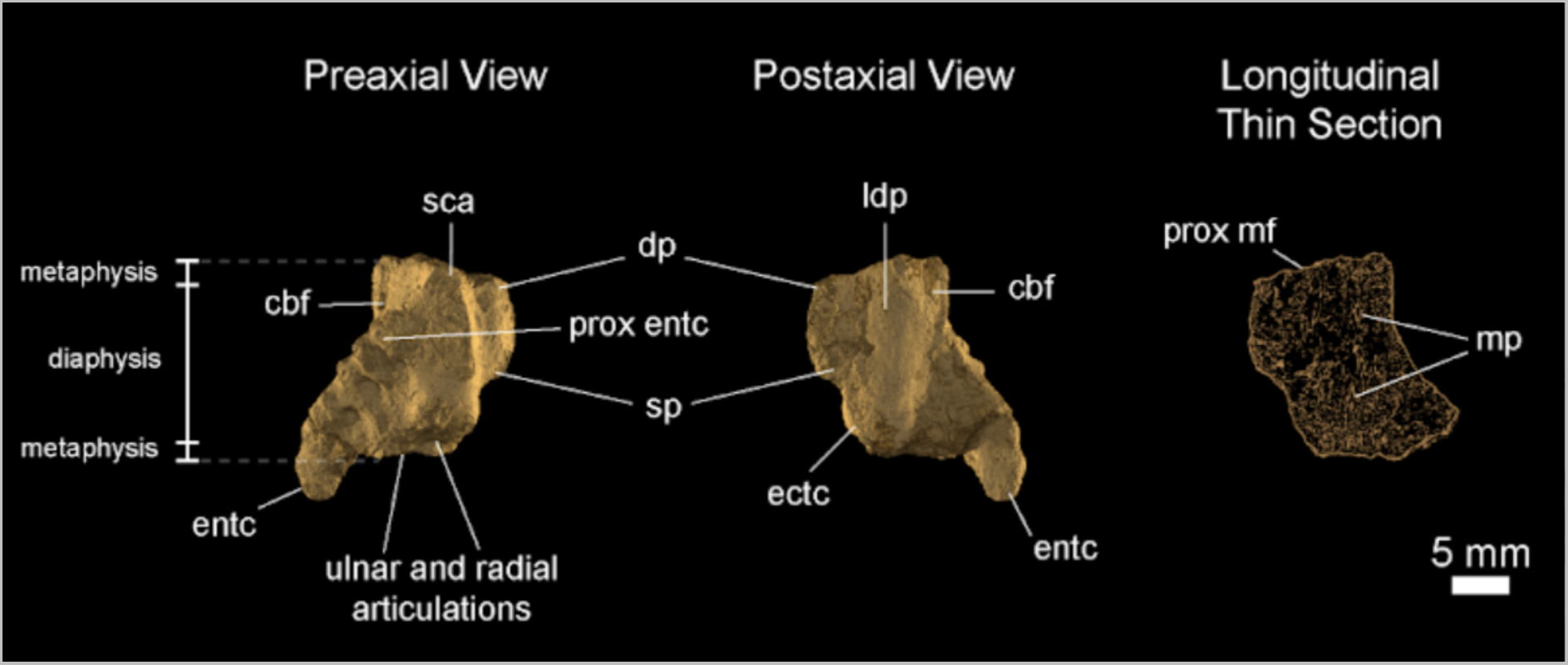
3D model and thin section of the humerus of *Eusthenopteron foordi*, in preaxial and postaxial views. Abbreviations: (cbf) coracobrachialis fossa; (dp) deltoid process; (ectc) ectepicondyle; (entc) entepicondyle; (ldp) latissimus dorsi process; (mp) marrow process; (prox entc) proximal tip of the entepicondyle; (prox mf) proximal mineralisation front; (sca) supracoracoidus attachment; (sp) supinator process.

In cross sections, Supplementary data 1 shows that higher von Mises stress are located in the periphery of the humerus in the regions of the deltoid process and coracobrachialis fossa. In “plain” condition, the centre of the humerus remains unstressed.

The “isotropic cancellous” model exhibits the same stress distribution, with the same intensity, as the “plain” model (Fig 9B). As in “plain” conditions, the humerus is slightly more stressed in the proximal region of the humerus, i.e. in the most proximal part of the deltoid process and coracobrachialis fossa (Fig 11; Figs 9B and 9A).

Cross sections (Suppl. data 2) exhibit the same stress distribution as in “plain” conditions (Suppl. data 1). The stress is concentrated in the proximal part of the humerus, in periphery of the deltoid process and coracobrachialis fossa. It reaches at most 1.5 MPa (Suppl. data 2). The rest of the inner core seems unstressed.

The “actual trabecular” model (Fig 9C) also shows von Mises stress in the same regions as the corresponding “plain” and “isotropic cancellous” models. However, the stress, there, is overall higher (locally appearing in red, i.e. reaching about 4 MPa) and far more extended than in the “plain” and “isotropic cancellous” models. Peaks of stress are located ventrally in the coracobrachialis fossa, dorsally on the proximal tip of the deltoid process, mesially on the postaxial facet of the deltoid process towards the latissimus dorsi process, as well as more distally, along the supinator process (Fig 11; Fig 9C). The cortical bone of the supracoracoideus attachment is also very locally stressed (up to 4 MPa). These stress peak decrease from 4 MPa (in red) to 2.5 MPa (in green) towards the distal region of the humerus. In the distal metaphysis of the humerus, the stress even reaches about 1 MPa (light blue). The ridge and the distal tip of the entepicondyle remain barely (if not) stressed (Fig 11; Fig 9C).

The stress not only spreads at the surface of the bone, but it penetrates the inner cancellous bone (Fig 9C, Suppl. data 3). Contrary to the surface of the bone, the stress in the cancellous bone is not restricted to the location of the processes and muscle attachment but it is distributed instead along the trabeculae. In longitudinal cross section (Fig 9C, Suppl. data 3), the trabecular network displays higher peaks of (1-2 MPa) stress along the longitudinal trabeculae (including the marrow processes, Fig 11), crossing the humerus from the distal to the proximal epiphysis (Fig 9C, Suppl. data 3).

#### 2) Lift-based mode (based on the pike’s body properties – *Esox lucius*)

As in drag-based conditions, the “plain model” in lift-based swimming mode (Fig 10A) shows a very low von Mises stress distribution (revealed by a blue colour-coded surface, reflecting a stress less than 1 MPa). Once again, in pre- and post-axial views, a local stress can be visualised in light blue on the radial and ulnar facet of the distal articulations due to the application of the ground reaction force at these points. Locally slightly higher-stress regions in the proximal part of the bone appear in light blue. These regions include the supracoracoideus attachment, and to a lesser extent, the coracobrachialis fossa (Fig 11; Fig 10A).

Multiple cross sections in the “plain” model of the humerus of *Eusthenopteron foordi* show that the stress stays in periphery in lift-based conditions (Fig 10A; Suppl. Data 4). No stress diffuses to the heart of the bone.

The “isotropic cancellous” model displays the same von Mises stress distribution as the “plain” model (Fig 10B and 10A). The stress intensity is very low (likewise the “plain” model) and localised in the proximal metaphysis of the bone (Fig 10B).

Longitudinal sections in this model (Suppl. Data 5) show that the stress does not penetrate the inner core of the bone. The distribution is strictly the same as for the “plain” model (Suppl. Data 5). As for the “plain” model, this is the least stress configuration in swimming locomotion.

In the “actual trabecular” model, the most stressed regions concentrate on the supracoracoideus attachment (with a peak of stress of 4 MPa on the tip), along the postaxial margin (with a higher intensity up to 2.5 MPa on the latissimus dorsi process), and to a lesser extent, on the shaft (Fig 10C; Suppl. Data 6). An average stress of 1 MPa disperses to the shaft from the proximal to the distal metaphysis (with local maxima at 2 MPa). The stress distribution extends up dorsally to the proximal tip of the deltoid process, as well as along the most proximal and ventral part of the entepicondyle ridge (Fig 10C). The entepicondyle ridge seems less stressed in lift-based than in drag-based mode. In both cases, the tip of the entepicondyle remains unstressed by the applied forces (Fig 10C and 9C).

In longitudinal section, the stress, once more, essentially circulates along the longitudinal trabeculae (including the marrow processes, Fig 11) (Fig 10C; Suppl. Data 6).

### Swimming models: quantitative comparisons

All von Mises-stress models have been configured with a scale ranging from 0 to 5 MPa (Fig 9,10; Suppl. data 1,2,3,4,5,6). Beyond this threshold, very local peaks of stress are displayed in white (e.g. in the “actual trabecular” model in drag-based mode; Fig 9C). The maximum values (table 3) can reach remarkably high levels of von Mises stress in swimming conditions (37.129 in drag-based mode for the trabecular model), but because they result from very limited local peaks restricted to few isolated finite elements (tetrahedrons; see material and methods for details), they are not relevant to the whole pattern of stress distribution. We deliberately chose to calculate the median stress for each model (i.e. from the 9 133 335 FE values of the “plain” model, and the 12 804 546 FE values of the “actual trabecular” model) to compare the different models through statistical analyses. Contrary to the mean stress, the median stress is not influenced by outlier values. After exporting the FE stress values for each model, we have compared all our model median stress values using a Wilcoxon signed-rank test. This comparison shows significant p-values when comparing the “plain” models with their respective “actual trabecular” models, as well as between both “actual trabecular” models in lift- and drag-modes (table 4). This means that the difference between these pairs is significant. However, there is no difference between the “plain” model and the “isotropic cancellous” model in either mode (Fig 12). Overall, the “actual trabecular” models display higher median von Mises stress than the “plain” models (Fig 12), with an increase by a factor of 3.63 for the drag-based mode and by a factor of 3.21 for the lift-based mode (Fig 12). It is also important to mention that the median stress is much higher in drag-based mode compared to the lift-based mode (in any configuration: “plain”, “isotropic cancellous” or “actual trabecular” version; table 3).

**Fig 12.**
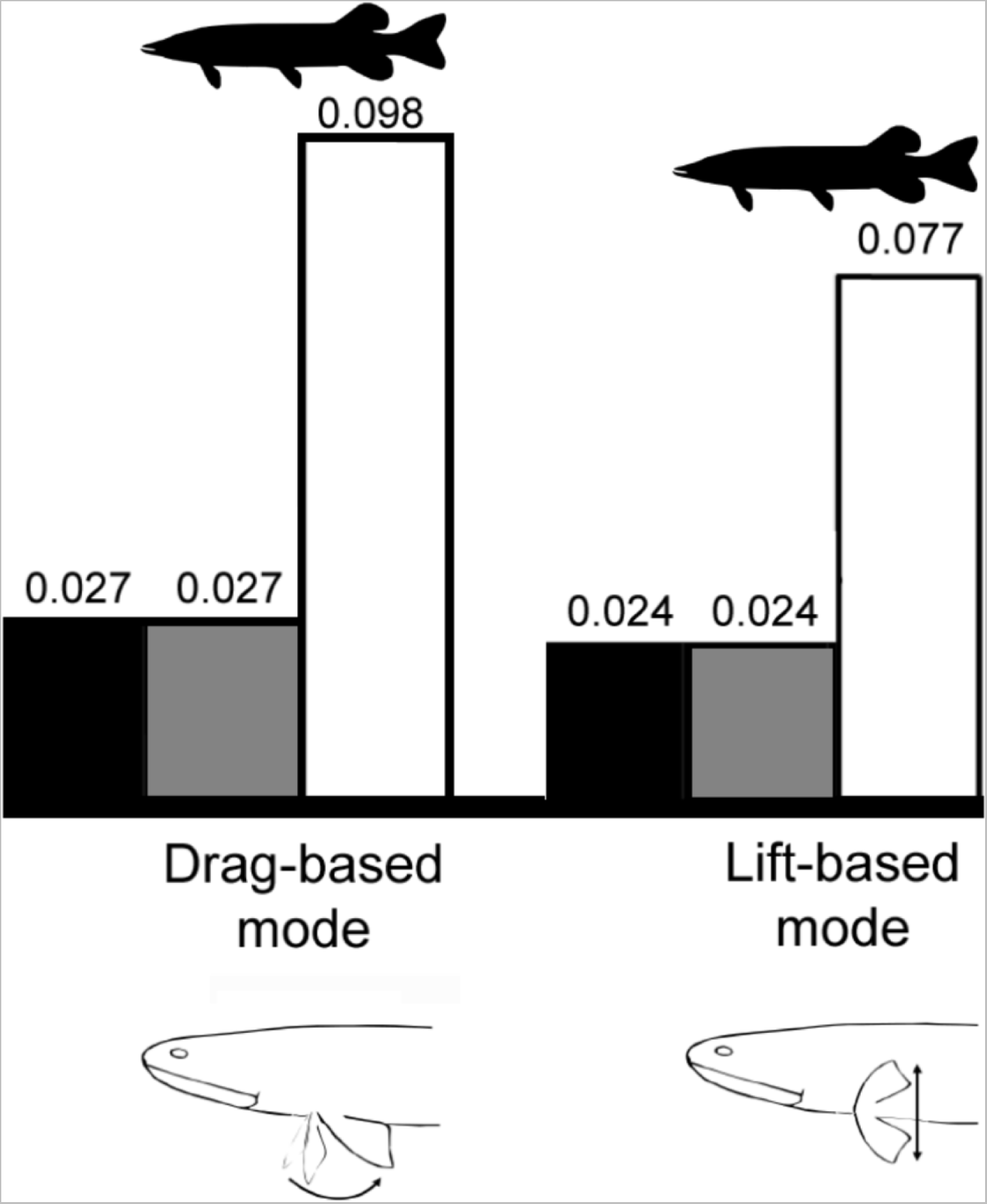
Median von Mises stress (MPa). Each graph plots the median stress for both the drag-based and lift-based swimming conditions based on the pectoral fin properties of the northern pike (*Esox lucius*), applied to the *Eusthenopteron foordi* humerus. The values for the “plain” models are illustrated as black columns, the “isotropic cancellous” models as gray columns, and the “actual trabecular” models as white columns.

**Table 3.** Statistical distribution of the von Mises stress in each finite-element model (MPa). Abbreviations: (P) “plain” model, (IsoC) “isotropic cancellous” model, (Tr) “actual trabecular” model, (*El*) for *Esox lucius* for the “Pike-based” models and (*Ph*) for *Phractocephalus hemioliopterus referring to the* “Catfish-based” model.

| Models | von Mises stress (MPa) |  |  |  |  |  |
| --- | --- | --- | --- | --- | --- | --- |
|  | Min | 1st.<br>Quartile | Median | Mean | 3rd.<br>Quartile | Max |
| **<u>Swimming conditions</u>** |  |  |  |  |  |  |
| *El* - (P) Drag based mode | 0.000 | 0.009 | 0.027 | 0.043 | 0.054 | 11.334 |
| *El* - (IsoC) Drag based mode | 0.000 | 0.009 | 0.027 | 0.043 | 0.054 | 11.334 |
| *El* - (Tr) Drag based mode | 0.000 | 0.034 | 0.098 | 0.175 | 0.222 | 37.129 |
| *El* - (P) Lift based mode | 0.000 | 0.008 | 0.024 | 0.034 | 0.049 | 4.142 |
| *El* - (IsoC) Lift based mode | 0.000 | 0.008 | 0.024 | 0.034 | 0.049 | 4.142 |
| *El* - (Tr) Lift based mode | 0.000 | 0.026 | 0.077 | 0.137 | 0.177 | 13.470 |
| **<u>Terrestrial conditions</u>** |  |  |  |  |  |  |
| *El* - (P) Straight fin | 0.000 | 0.010 | 0.006 | 0.009 | 0.015 | 1.238 |
| *Ph* - (P) Straight fin | 0.000 | 0.007 | 0.020 | 0.033 | 0.051 | 4.340 |
| *El* - (IsoC) Straight fin | 0.000 | 0.010 | 0.006 | 0.009 | 0.015 | 1.238 |
| *Ph* - (IsoC) Straight fin | 0.000 | 0.007 | 0.020 | 0.033 | 0.051 | 4.340 |
| *El* - (Tr) Straight fin | 0.000 | 0.010 | 0.016 | 0.036 | 0.044 | 7.992 |
| *Ph* - (Tr) Straight fin | 0.000 | 0.018 | 0.057 | 0.126 | 0.153 | 28.021 |
| *El* - (P) 50° bent elbow | 0.000 | 0.008 | 0.023 | 0.031 | 0.046 | 3.918 |
| *Ph* - (P) 50° bent elbow | 0.000 | 0.027 | 0.079 | 0.108 | 0.160 | 13.736 |
| *El* - (IsoC) 50° bent elbow | 0.000 | 0.008 | 0.023 | 0.031 | 0.046 | 3.918 |
| *Ph* - (IsoC) 50° bent elbow | 0.000 | 0.027 | 0.079 | 0.108 | 0.160 | 13.736 |
| *El* - (Tr) 50° bent elbow | 0.000 | 0.020 | 0.064 | 0.113 | 0.148 | 9.495 |
| *Ph* - (Tr) 50° bent elbow | 0.000 | 0.075 | 0.220 | 0.387 | 0.506 | 32.515 |

**Table 4.**
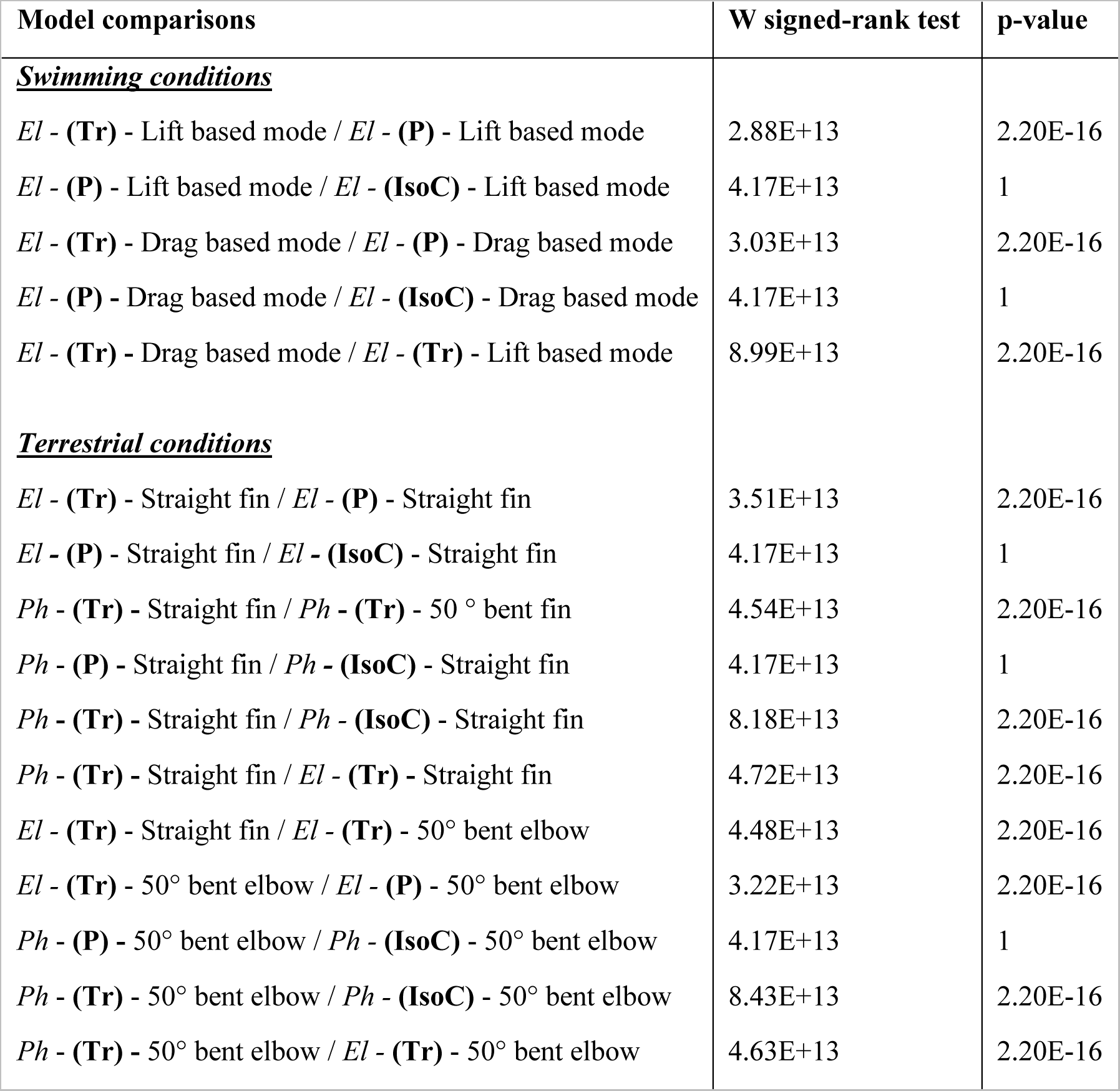
Wilcoxon signed-rank test comparison between Finite Element models (MPa). Abbreviations: (P) “plain” model, (IsoC) “isotropic cancellous” model, (Tr) “actual trabecular” model, (*El*) for *Esox lucius* for the “Pike-based” models and (*Ph*) for *Phractocephalus hemioliopterus referring to the* “Catfish-based” model.

| Model comparisons | W signed-rank test | p-value |
| --- | --- | --- |
| **<u>Swimming conditions</u>** |  |  |
| *El* - (Tr) - Lift based mode / *El* - (P) - Lift based mode | 2.88E+13 | 2.20E-16 |
| *El* - (P) - Lift based mode / *El* - (IsoC) - Lift based mode | 4.17E+13 | 1 |
| *El* - (Tr) - Drag based mode / *El* - (P) - Drag based mode | 3.03E+13 | 2.20E-16 |
| *El* - (P) - Drag based mode / *El* - (IsoC) - Drag based mode | 4.17E+13 | 1 |
| *El* - (Tr) - Drag based mode / *El* - (Tr) - Lift based mode | 8.99E+13 | 2.20E-16 |
| **<u>Terrestrial conditions</u>** |  |  |
| *El* - (Tr) - Straight fin / *El* - (P) - Straight fin | 3.51E+13 | 2.20E-16 |
| *El* - (P) - Straight fin / *El* - (IsoC) - Straight fin | 4.17E+13 | 1 |
| *Ph* - (Tr) - Straight fin / *Ph* - (Tr) - 50 ° bent fin | 4.54E+13 | 2.20E-16 |
| *Ph* - (P) - Straight fin / *Ph* - (IsoC) - Straight fin | 4.17E+13 | 1 |
| *Ph* - (Tr) - Straight fin / *Ph* - (IsoC) - Straight fin | 8.18E+13 | 2.20E-16 |
| *Ph* - (Tr) - Straight fin / *El* - (Tr) - Straight fin | 4.72E+13 | 2.20E-16 |
| *El* - (Tr) - Straight fin / *El* - (Tr) - 50° bent elbow | 4.48E+13 | 2.20E-16 |
| *El* - (Tr) - 50° bent elbow / *El* - (P) - 50° bent elbow | 3.22E+13 | 2.20E-16 |
| *Ph* - (P) - 50° bent elbow / *Ph* - (IsoC) - 50° bent elbow | 4.17E+13 | 1 |
| *Ph* - (Tr) - 50° bent elbow / *Ph* - (IsoC) - 50° bent elbow | 8.43E+13 | 2.20E-16 |
| *Ph* - (Tr) - 50° bent elbow / *El* - (Tr) - 50° bent elbow | 4.63E+13 | 2.20E-16 |

### Terrestrial models: qualitative comparisons

Figures 13 to 18 show the von Mises stress distribution through the humerus resulting from the virtual application of the ground reaction force at the elbow joint. This test aims at simulating hypothetical terrestrial constraints on the humerus of *Eusthenopteron foordi*. As mentioned earlier, we have considered two hypothetical scenarios on the basis of Andrew and Westoll’s (24) suggestions: 1) the pectoral fin lays straight along the proximo-distal axis of the humerus (Fig 8A); 2) the pectoral fin is vertically bent down by an angle of 50° at the elbow joint (Fig 8B). Both cases were illustrated by three simulations: a “plain” model represented as a full homogeneous volume of compact bone matrix (Young’s modulus of 18 GPa), an “isotropic cancellous” model (based on a Young’s modulus of 13.5 GPa) and an “actual trabecular” model (based on segmented trabeculae). Because the animal’s weight plays a major role in the terrestrial configuration, two analogous fishes were considered here for estimating the weight of the specimen of *E. foordi*: 1) the northern pike (*Esox lucius*) (Fig 13,14; Suppl. Data 7-12) whose body length/weight ratio is probably the most similar to that of *E. foordi*, and 2) the catfish (*Phractocephalus hemioliopterus*) (Fig 15,16,17,18; Suppl. data 13-24) whose weight is much larger and would therefore represent the maximum weight a fish with the length of *E. foordi* could reach (see material and methods for details). Because stress peaks exceeded the 5-MPa limit in the catfish configuration, the scale was increased to 15 MPa in figures 16 and 18.

**Fig 13.**
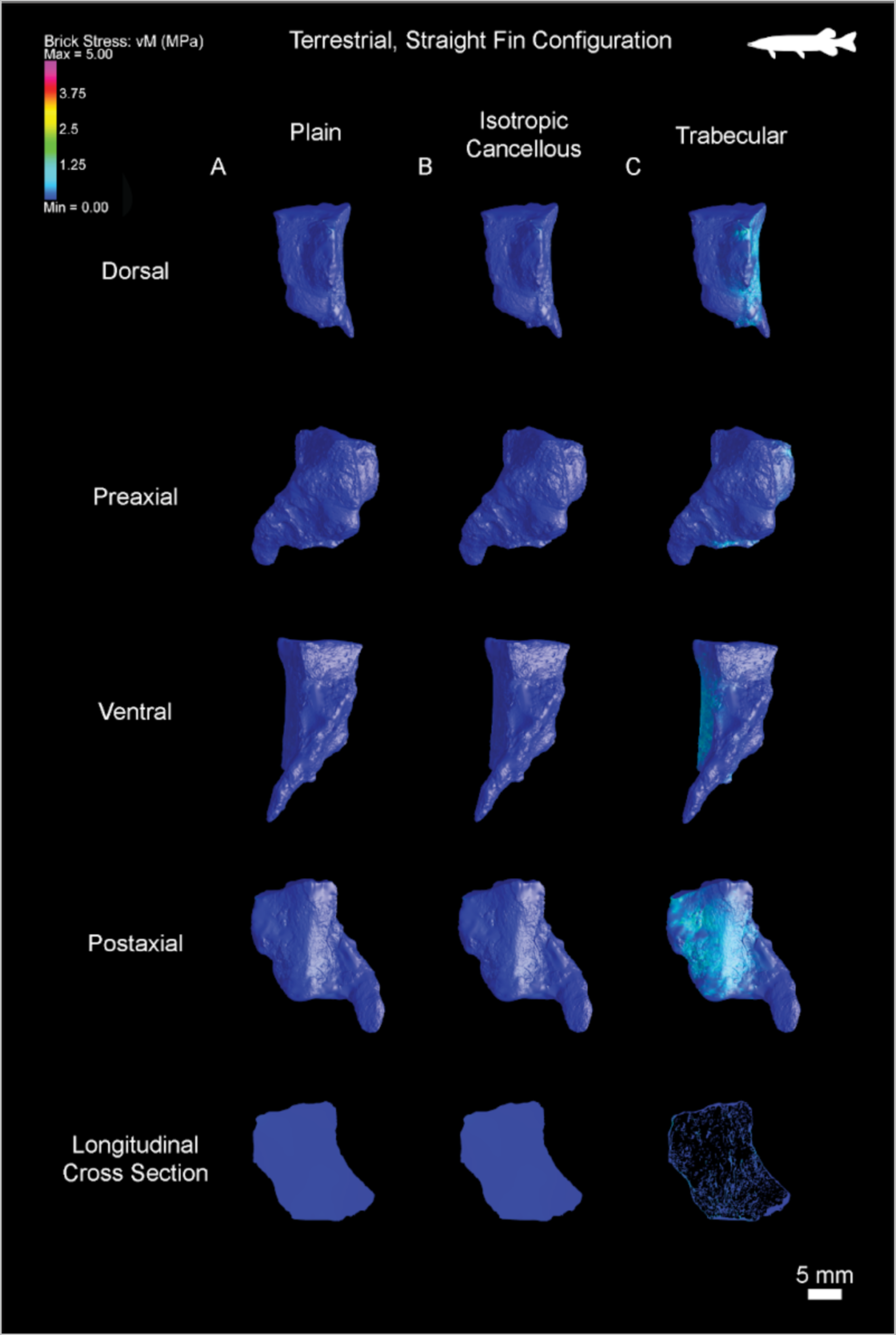
Distribution of the von Mises stress in the simulated terrestrial, straight-finned, model based on the pectoral fin properties of the northern pike (*Esox lucius*), applied to the *Eusthenopteron foordi* humerus. Stress distribution from 0-5 MPa. Four views and a longitudinal cross section are shown. (A) ‘’Plain’’ mode; (B) Isotropic cancellous model; (C) Trabecular mesh model.

**Fig 14.**
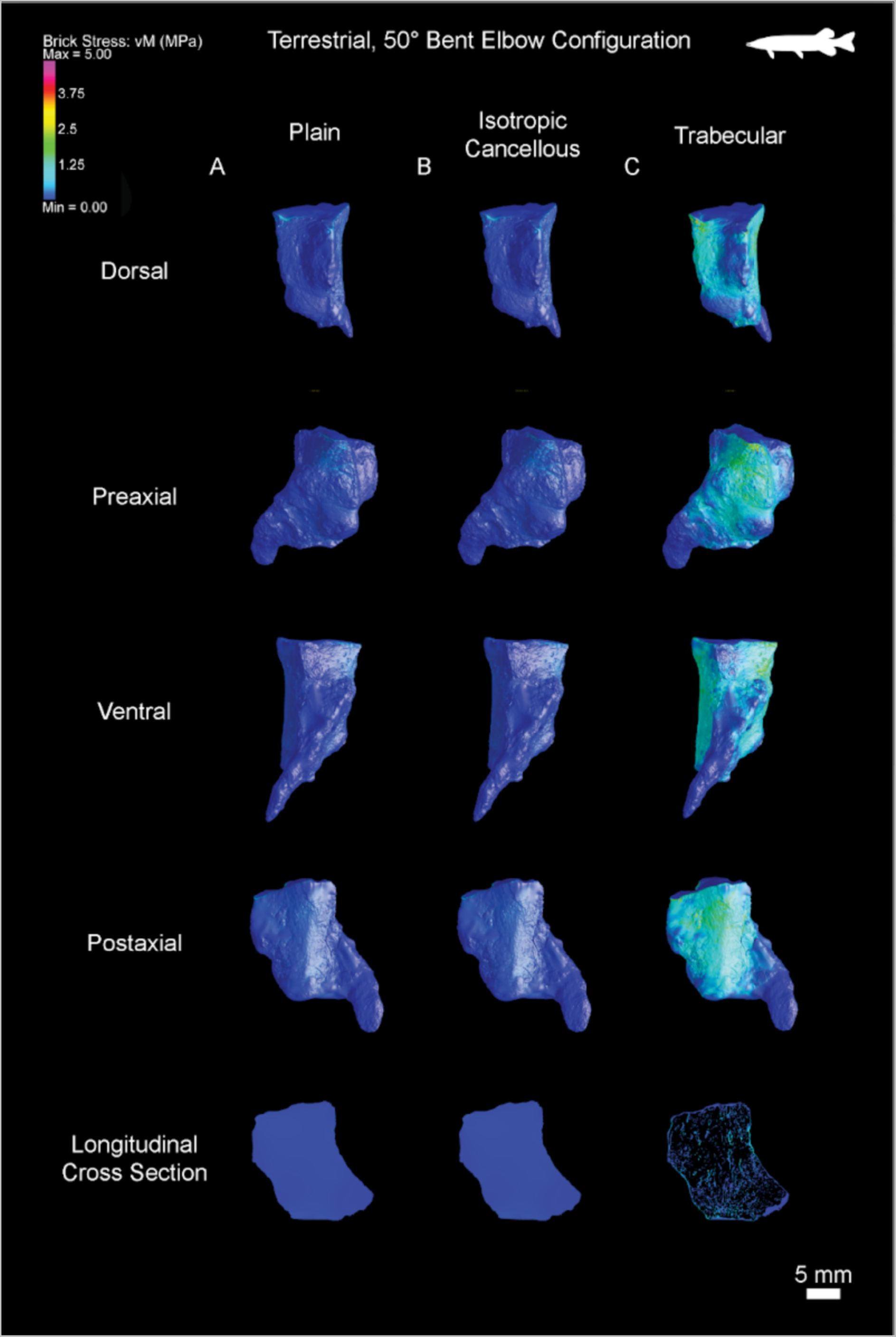
Distribution of the von Mises stress in the simulated terrestrial, elbow bent at 50°, model based on the pectoral fin properties of the northern pike (*Esox lucius*), applied to the *Eusthenopteron foordi* humerus. Stress distribution from 0-5 MPa. Four views and a longitudinal cross section are shown. (A) ‘’Plain’’ mode; (B) Isotropic cancellous model; (C) Trabecular mesh model.

**Fig 15.**
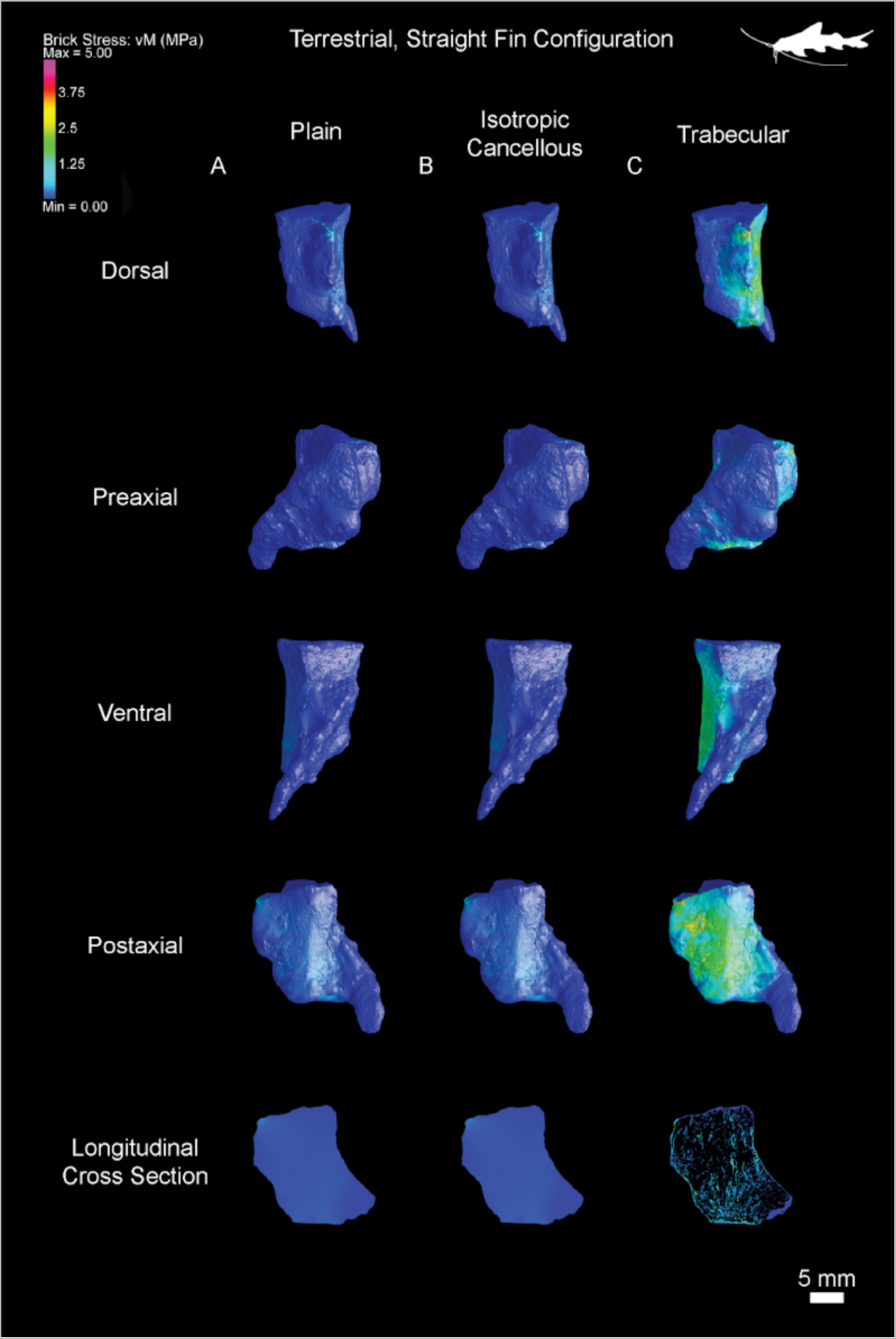
Distribution of the von Mises stress in the simulated terrestrial, straight-finned, model based on the pectoral fin properties of the redtail catfish (*Phractocephalus hemioliopterus*), applied to the *Eusthenopteron foordi* humerus. Stress distribution from 0-5 MPa. Four views and a longitudinal cross section are shown. (A) ‘’Plain’’ mode; (B) Isotropic cancellous model; (C) Trabecular mesh model.

**Fig 16.**
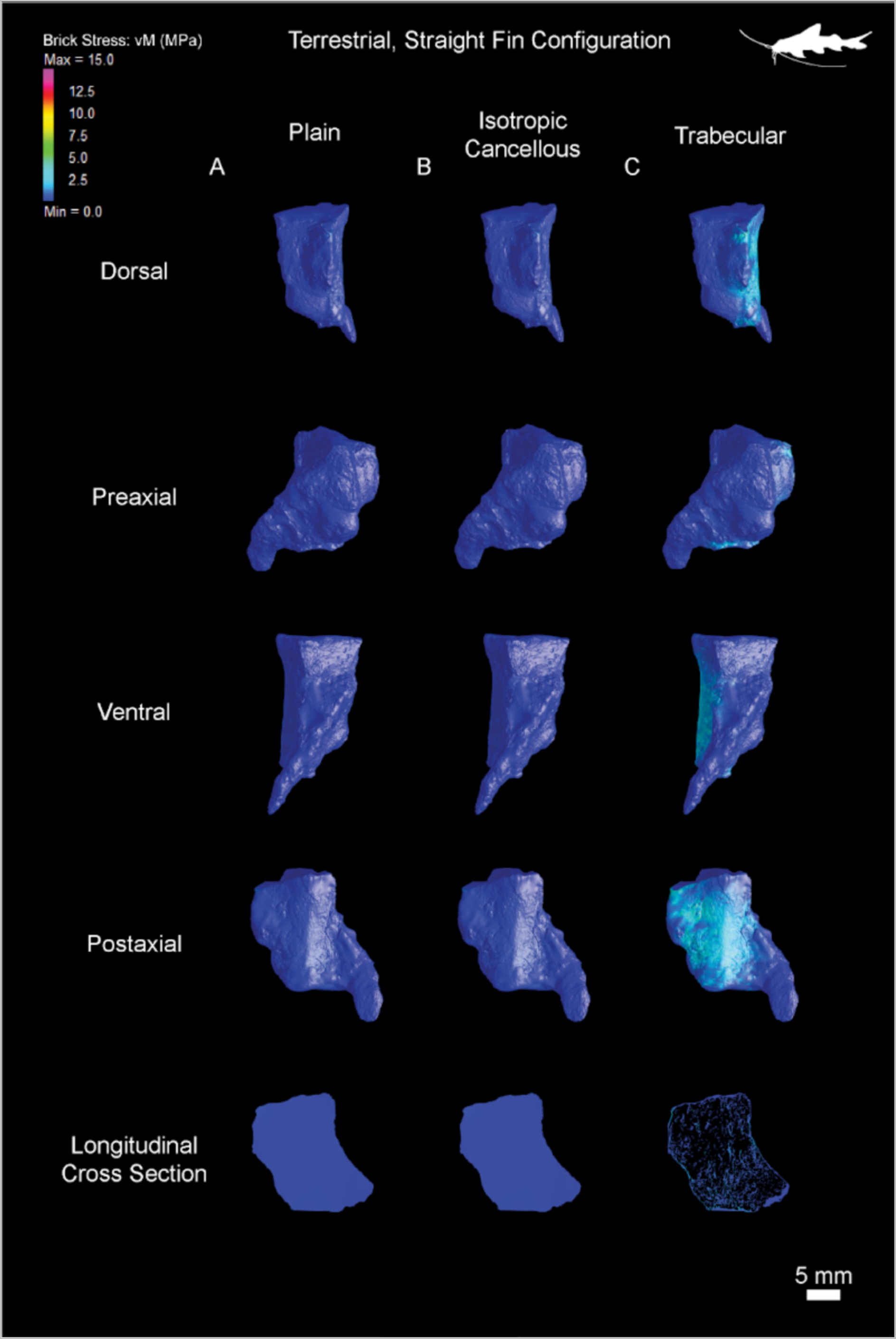
Distribution of the von Mises stress in the simulated terrestrial, straight-finned, model based on the pectoral fin properties of the redtail catfish (*Phractocephalus hemioliopterus*), applied to the *Eusthenopteron foordi* humerus. Stress distribution from 0-15 MPa. Four views and a longitudinal cross section are shown. (A) ‘’Plain’’ mode; (B) Isotropic cancellous model; (C) Trabecular mesh model.

**Fig 17.**
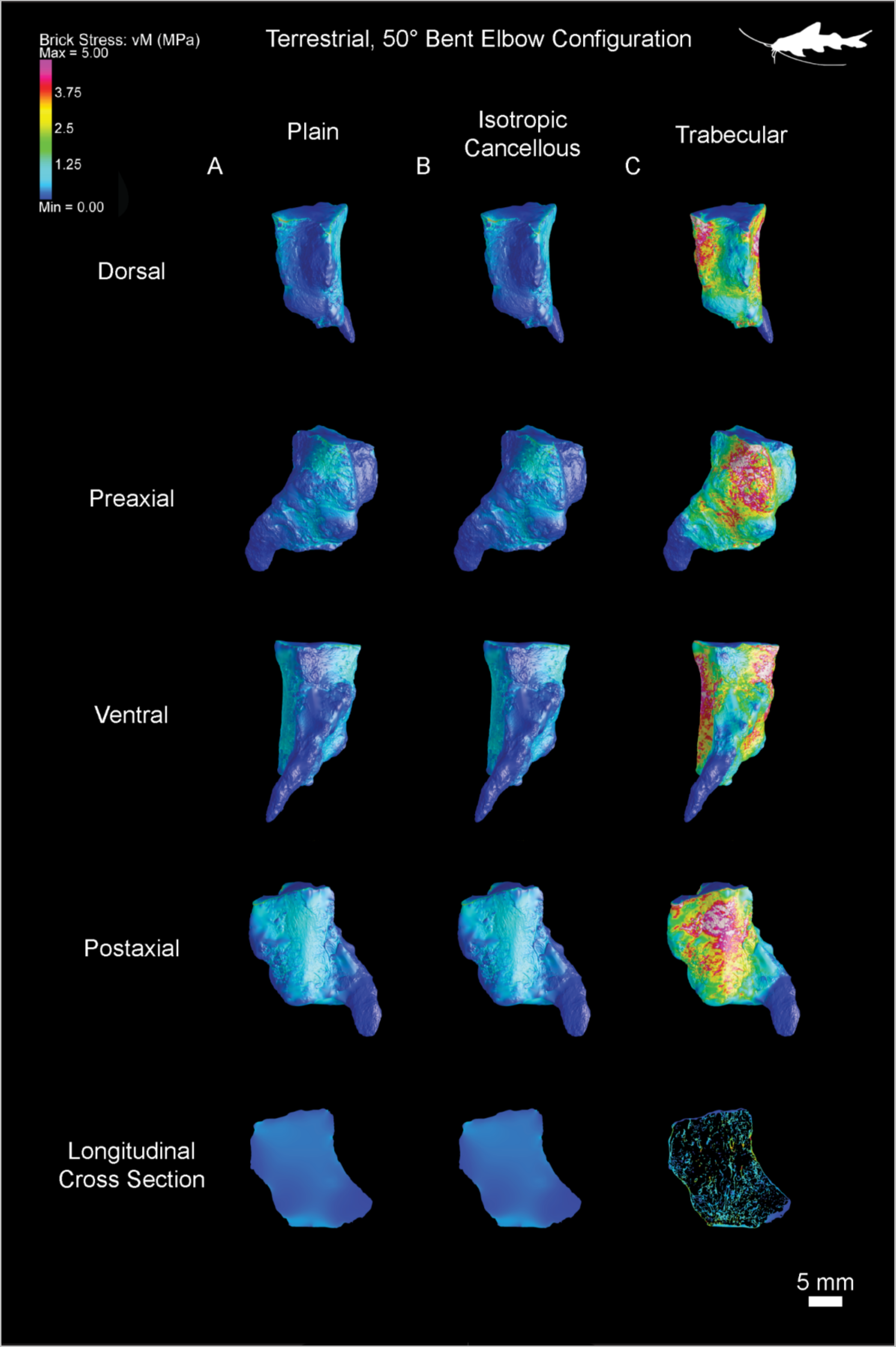
Distribution of the von Mises stress in the simulated terrestrial, elbow bent at 50°, model based on the pectoral fin properties of the redtail catfish (*Phractocephalus hemioliopterus*), applied to the *Eusthenopteron foordi* humerus. Stress distribution from 0-5 MPa. Four views and a longitudinal cross section are shown. (A) ‘’Plain’’ mode; (B) Isotropic cancellous model; (C) Trabecular mesh model.

**Fig 18.**
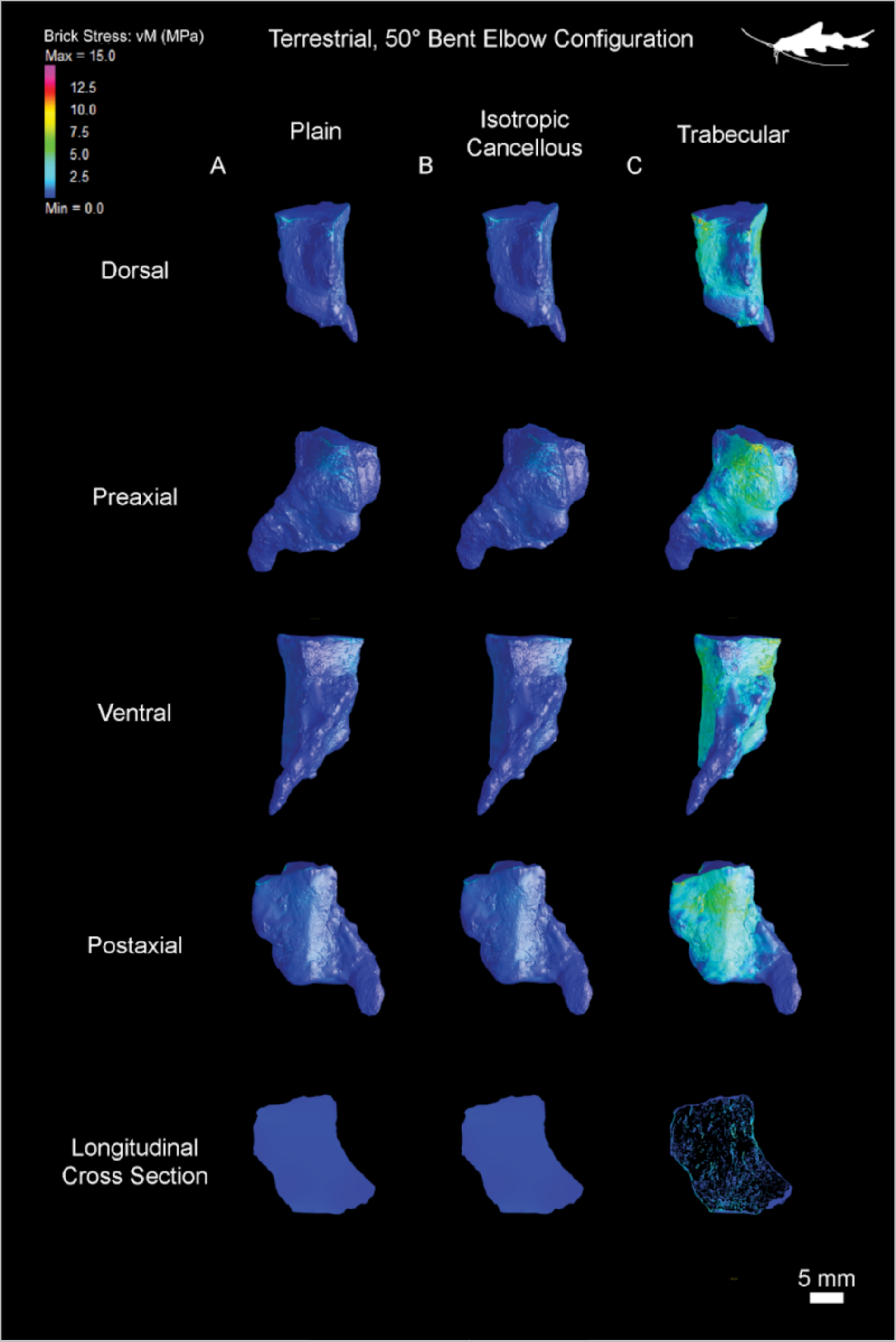
Distribution of the von Mises stress in the simulated terrestrial, elbow bent at 50°, model based on the pectoral fin properties of the redtail catfish (*Phractocephalus hemioliopterus*), applied to the *Eusthenopteron foordi* humerus. Stress distribution from 0-15 MPa. Four views and a longitudinal cross section are shown. (A) ‘’Plain’’ mode; (B) Isotropic cancellous model; (C) Trabecular mesh model.

#### 1) Straight-fin configuration based on the pike’s body properties (*Esox lucius*)

The “plain” model (Fig 13A) globally appears in dark blue, i.e. it displays a very low level of stress. There is no distinguishable pattern of higher stress distribution on the morphological view of the humerus (Fig 13A) nor on the cross sections (Suppl. data 7). There is a very limited local point of low stress appearing in light blue (1 MPa) on the proximal tip of the deltoid process (Fig 11; Suppl. data 7).

The “isotropic cancellous” model exhibits exactly the same stress distribution as the “plain” model in morphological view, as well as through longitudinal sections (Fig 13B; Suppl. data 8).

The “actual trabecular” model (Fig 13C; Suppl. data 9) shows a clear increase in the von Mises stress on the postaxial ridge of the humerus (in light blue – reaching a maximum of about 1.25 MPa). The stress distribution extends to the distal region of the ectepicondyle, dorsally to the deltoid process, and stops ventrally at the location of the coracobrachialis fossa (Fig 11; Fig 13C). The preaxial and ventral facets of the humerus are almost not stressed. Surprisingly, the most proximal region of the humerus metaphysis seems much less stressed (except for the deltoid process) than the distal metaphysis.

The longitudinal sections illustrate little stress spreading in the trabecular mesh through the longitudinal trabeculae (Suppl. data 9). The stress remains essentially on the surface of the bone, in the cortex, at the location of the postaxial ridge (Suppl. data 9). A local stress can be visualised in light blue in preaxial view on the radial and ulnar facet of the distal articulations due to the application of the ground reaction force at these points.

Supplementary data Figure S1 is provided for the simulation with the analogous fish *Esox lucius* for comparison purposes with the stress distribution on the humerus of *Eusthenopteron foordi* constrained with the weight of *Phractocephalus hemioliopterus*. Because Supplementary data Figure S1 does not provide more details than Figure 13, Supplementary data Figure S1 won’t be described here.

#### 2) 50° bent-elbow fin configuration based on the pike’s body properties (*Esox lucius*)

The “plain” model (Fig 14A, Suppl. data 10) appears once more very dark blue. It exhibits some little stress (around 1MPa) in light blue in the proximal region of the supracoracoideus attachment, as well as at the mineralisation front above the deltoid process (Fig 11; Fig 14A).

The longitudinal sections (Suppl. data 10) display no stress at all in the core of the bone.

The “isotropic cancellous” model exhibits the exact same stress pattern as the “plain” model (Fig 14B). The stress is essentially restricted to the supracoracoideus attachment.

The longitudinal sections display no stress diffusion in the inner region of the humerus (Suppl. data 11).

The “actual trabecular” model (Fig 14C; Suppl. data 12) shows a global increase of stress when the fin is bent at 50°. This mainly comes out in light blue, thereby suggesting a 1-to-2.5 MPa stress (Fig 14C). It is distributed on the entire surface of the humerus, except at the tip of the entepicondyle. Surprisingly, the deltoid process, the proximal region of the entepicondyle ridge, as well as the proximal region to the coracobrachialis fossa are less stressed (with dark-blue patches; Fig 14C). On the contrary, the supracoracoideus attachment and the latissimus dorsi process are more stressed (in green-yellow: 1.5-to-3 MPa; Fig 14C).

The longitudinal virtual thin sections show that the stress dissipates along the longitudinal trabeculae, and more specifically along the marrow processes through the entire mesh (Suppl. data 12; Fig 11). Within the trabecular mesh, the stress does not exceed 1.25 MPa.

Supplementary data Figure S2 is provided for comparison purposes with the next sections based on the catfish’s body properties (*Phractocephalus hemioliopterus*). Because Supplementary data Figure S2 does not provide more details than Figure 14, Supplementary data Figure S2 won’t be described here.

#### 3) Straight-fin configuration based on the catfish’s body properties (*Phractocephalus hemioliopterus*)

Due to extremely local stress peaks higher than 5 MPa in the trabecular models in these conditions, some stress saturates in white peaks at the surface of the humerus (Fig 15C). Because the colour saturation limits the observation of stress distribution, we have decided to increase the scale range up to 15 MPa (Fig 16). This will allow us to visualise the stress distribution in the less stressed regions, as well as in the areas of high-peak stress (Fig 16). As a control we ran cross-sectional movies in all three configurations at both 5MPa (Suppl. data 13,14,15) and 15MPa (Suppl. data 16,17,18).

The “plain” model (Fig 15A, Suppl. data 13) appears slightly more stressed on the postaxial ridge than in any other parts of the humerus (Fig 15A). The maximum peaks of stress are restrained to tiny area located on the proximal tip of the deltoid process, the postaxial facet of the deltoid process, along the postaxial ridge and the ulnar facet (with an intensity not higher than 1 MPa; Fig 15A). The preaxial region and the entepicondylar ridge are not stressed.

Longitudinal virtual thin sections in the humerus show that the stress diffusion into the core of the bone (in light blue) essentially comes from the tip of the deltoid process with an intensity of about 1-1.5 MPa (Suppl. data 13). This peak of stress only slightly propagates in periphery to this area (Suppl. data 13).

The stress distribution in the “isotropic cancellous” model is equally displayed as in the “plain” model (Fig 15B). The stress intensity is the same as well.

The through-out movie in the “isotropic cancellous” model Suppl. data 14 shows that the inner diffusion of von Mises stress in the core of the bone is similar to the pattern obtained in “plain” conditions.

It is important to note that the “trabecular” model based on the morphology and weight of *Phractocephalus hemioliopterus* (Fig 15C; Suppl. data 15 and 18) shows the same pattern of stress distribution as the “trabecular” model based on the morphological settings of *Esox lucius* in terrestrial conditions in the configuration of a straight fin (Fig 13C). Nonetheless, an increase in stress intensity is clearly noticeable (Fig 13C and 15C, this is even more obvious when compared to Supplementary data Figure S1C; Suppl. data 9,15,18). The stress along the postaxial ridge averages 2 MPa with spots of higher values mostly located on the postaxial facet of the deltoid process, on the proximal tip of the deltoid process, and to a lesser extent on the postaxial ridge (Fig 15C). The ulnar articular facet is also slightly stressed (around 1 MPa; Fig 16C).

Longitudinal thin sections made in the “trabecular” model of the humerus show a diffusion of the stress in the vicinity of the postaxial border (Fig 15C and 16C; Suppl. data 15 and 18). Even though the longitudinal trabeculae (including the marrow processes) absorb most of the stress (1.5 MPa, Suppl. data 18), the rest of the trabeculae also participate in the distribution of the mechanical constraints (Fig 15C).

#### 4) 50° bent configuration based on the catfish’s body properties (*Phractocephalus hemioliopterus*)

This is the most stressed configuration; therefore, we have also produced figures and cross-sectional movies using 5MPa and 15MPa scales due to extremely local stress peaks (Fig 17 and Fig 18; Suppl. data 19,20,21,22,23,24).

The “plain” model shows the most stressed configuration of all “plain” models investigated in this study (Fig 17A). The stress applied in the preaxial and postaxial views reaches an average value of 1-1.5 MPa (Fig 17A). Locally in the proximal region of the humerus (especially at the location of the mineralisation front above the deltoid process and the supracoracoideus attachment), spots of stress reach about 3 MPa (Fig 17A). Surprisingly, the deltoid process itself, as well as the entepicondyle ridge and the coracobrachialis fossa are the least stressed regions in these conditions (less than 1 MPa, Fig 17A).

Longitudinal thin sections made in the inner core of the humerus show that the stress radiates inside the bone except in the entepicondyle (Fig 17A; Suppl. data 19 and 22). It is more intense in the vicinity of the the pre- and post-axial margins, as well as at the location of the growth plates associated with the ulnar articular facet (1-1.5 MPa, Fig 17A; Suppl. data 19 and 22).

Once again the stress distribution in the “isotropic cancellous” model is equal to that of the “plain” model in both distribution and intensity at the surface, as well as in the inner core of the humerus (Fig 17B;18B; Suppl. data 20 and 23).

The “trabecular” model of the humerus in this configuration is the most constrained (Fig 17C and 18C). Because the stress at the surface outstrips the scale bar limit (Fig 17C), a 15-MPa scale bar has been produced (Fig 18C). The major peaks of stress are displayed on the pre- and post-axial margins of the humerus, with an intensity surpassing 7 MPa in the proximal region of the bone at the location of the latissimus dorsi process and 10 MPa in the vicinity of the supracoracoideus attachment (Fig 18C).

As for the other configurations, von Mises stress distributes internally with less intensity than at the surface. The maximum stress attains a value of 4-5 MPa with local peaks of 7 MPa in the trabecular mesh and concentrate on the longitudinal trabeculae, including the marrow processes (Suppl. data 23 and 24).

### Terrestrial models: quantitative comparisons

Von Mises stress values were collected in all configurations and summed up in quartiles in table 3. The latter shows that the highest stress values are encountered in the trabecular configurations with the catfish as a weight-to-length ratio analog: 28.021 for the mode with a straight fin; 32.515 for the mode with the bent fin (table 3). The other stress values are all lower than 15 MPa (table 3). These extremely high values probably only represent isolated voxels as all median values range under 1 MPa (table 3). The highest median value by far is that of the 50° bent-elbow configuration (with the catfish as analog for the weight) with a number of 0.387. In the straight-fin configuration, only the “trabecular” model – with the catfish as a weight analog-exceeds the value of 0.1 MPa. In the bent configuration, all models with the catfish as a weight analog have median stress values higher than 0.1 MPa. In the case of the northern pike as a weight analog, only the trabecular one exceeds 0.1 MPa.

Statistical comparisons were made between these models using a Wilcoxon signed-rank test (table 4). All comparisons were significantly different except the comparisons between “plain” and “isoC” models (table 4), thereby meaning that only these pairs were not significantly different from each other (Fig 19), as observed in swimming conditions too (Fig 12). When the pike is taken as a weight analog, the stress increases by a factor of 2.75 ± 0.5 in the “actual trabecular” model in any configuration. When the catfish is taken as a weight analog, the stress increases by a factor of 2.8 ± 0.5 in the “actual trabecular” model in any configuration (Fig 19).

**Fig 19.**
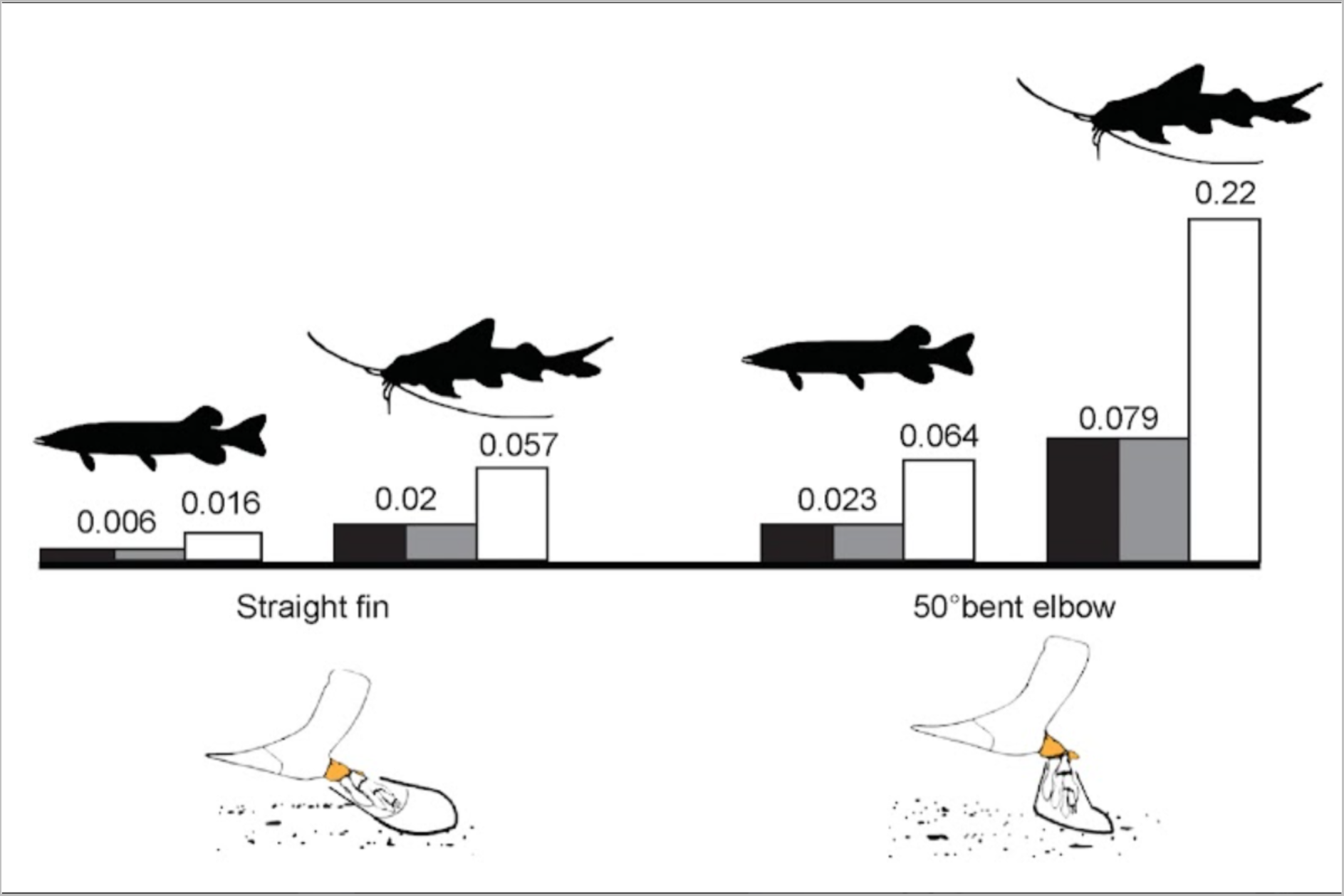
Median von Mises stress (MPa). Each graph plots the median stress for the simulated terrestrial conditions, straight-finned and with a bent elbow at 50°, based on the pectoral fin properties of the northern pike (*Esox lucius*) and the redtail catfish (*Phractocephalus hemioliopterus*) respectively, applied to the *Eusthenopteron foordi* humerus. The values for the “plain” models are illustrated as black columns, the “isotropic cancellous” models as gray columns, and the “actual trabecular” models as white columns.

## DISCUSSION

### Finite Element Analysis – considering the trabecular arrangement in long bones of stem tetrapods

Over about 400 million years, the evolution of the tetrapod limb has displayed a very large variety of morphologies and microanatomies adapted to various lifestyles and locomotory behaviours (5,51–57). Long bones are formed of compact and cancellous mineralised material (58,59). The cancellous bone displays a trabecular architecture, supposed to reflect the main external load orientation (49,60–68). However, because many other parameters such as body mass/size, long-bone regionalization, ageing and/or hormonal remodeling play a role in the trabecular arrangement (e.g., physiology (69); fatigue microfractures (70,71); phosphocalcic homeostasis and endocrine regulation, (72–75); ageing, (76–78); mass, (79)), the interpretation of the trabecular pattern has been very complex. Many different types of biomechanical studies – including Finite Element Analyses (FEA) – have been carried out on local trabecular regions of interest – especially in the biomedical field – to interpret the factors shaping the trabecular mesh (80–86), but very few FEA have considered combining microanatomy and morphology at a larger scale due to the heavy data it generates. In the few cases FEA was performed on an entire bone considering the 3D trabecular architecture (87,88), contradictory conclusions emerged on the importance of the trabecular pattern for understanding von Mises stress distribution. For these reasons, most FEA have been conducted on bone solid/plain models and this has been sufficient for understanding the skeletal stress patterns in these cases (89–92).

*Eusthenopteron* seems to be a perfect study case for checking the importance of the trabecular mesh in FEA: 1) *Eusthenopteron* is of medium size (with the adult reaching 1-to-1.5 m) (26,93), 2) the humeral micro-anatomy of *Eusthenopteron* is not yet really regionalised (15), and 3) the spongy core in the long bone of *Eusthenopteron* keeps a relatively similar orientation and arrangement during their development (15). Three models were therefore tested on an adult specimen of *Eusthenopteron* to check whether the microanatomical arrangement actually played a role in stress distribution and absorption. A “plain” model was considered with an overall Young modulus of 18 GPa. This is a common practice in FEA studies (94). An “isotropic cancellous” model was investigated with the Young modulus set to 13.5 GPa (46). This model assumed that the orientation and arrangement of the trabeculae was isotropic, i.e. had no preferential direction. Finally, the “actual trabecular” mesh with the exact trabecular arrangement was segmented and tested with a Young modulus of 18 GPa (standard value in Strand7 and (94)). The results show that the peaks of stress are always situated at the same locations whatever the model within a single configuration (Suppl. data Figure S3). This means that the morphology of the humerus would be the main structure conducting the stress at the surface of the cortex – whatever the Young modulus used (between at least 13.5 and 18 GPa) and the orientation of the trabecular arrangement in the humerus. It is worth mentioning though that the “plain” and “isotropic cancellous” models greatly undervalue the stress the bone endures: in swimming conditions, the stress in the “plain” and “isotropic cancellous” models was underestimated by 29% (± 2%) (Fig 12) and in terrestrial conditions, the stress in the “plain” and “isotropic cancellous” models was underestimated by 36% (± 1%) (Fig 19). The “trabecular” model therefore is the most reliable model to measure quantitatively the stress level the humerus actually undergoes. This proportional difference in median stress value (ΔMPa) between models remains moderate in the current study case of a medium-sized stem-tetrapod fish like *Eusthenopteron*, but this ΔMPa would likely have larger consequences if stress estimates were to be made in the skeletal elements of an animal of greater body mass (for which the yield strength of the matrix in tension could reach 114 MPa and the breaking stress in tension, 133 MPa; (49,95).

### Role of long-bone trabeculae in stress dissipation in the humerus of *Eusthenopteron foordi*

Longitudinal cross sections were made in the three models to understand this discrepancy and reveal the stress distribution inside the bone. The “plain” and “isotropic cancellous” models revealed a diffused but localised pattern around the high peaks of stress (e.g., Suppl. data 19, 22). The “trabecular” model however revealed higher stress values diffusing along the tubular walls of longitudinal canals (e.g., Suppl. data 24) identified in previous studies as marrow processes (15,18). The theory of “bone’s mechanostat” proposed by Frost (96,97) suggests that bone mass tends to increase in overloaded regions and decrease in low strained regions through remodelling to minimise the strain energy (98). Because the “plain” and “isotropic cancellous” models artificially comprise a homogeneous bone filled in with a large mass of mineralised material, they artificially absorb more energy, thereby restraining the diffusion of the stress within the bone. The “trabecular” model therefore is the only one able to reveal the real diffusion of von Mises stress within the trabecular mesh.

In swimming conditions, in drag-based mode, the stress diffuses deeply into the trabecular mesh and spreads longitudinally through multiple marrow processes running in the vicinity of the peaks of stress. In lift-based mode, however, the peaks of stress are of lesser intensity and the stress remains locally constrained at the surface of the humerus. Consequently, fewer marrow processes get involved in stress dissipation. The main environmental load in both swimming condition (drag- and lift-based modes) come from the water flow during swimming (Fig 4). The “trajectorial theory” (60,99) suggesting that the orientation of the trabecular mesh would reflect the main environmental load direction has received strong support from experimental and theoretical studies on terrestrial animals (49,65,80,100–102). Although the water-flow load in our simulation is more or less orthogonal to the proximo-distal axis of the humerus in both modes, the main arrangement of the trabecular mesh is not oriented in this direction but aligns instead with the proximo-distal axis of the humerus. The “trajectorial theory” (60,99) therefore does not seem to apply here in aquatic conditions. The literature also shows no preferential orientation in longitudinal thin sections of femora and humeri of marine mammals (such as cetaceans, (55,103)) – apart from longitudinal metaphyseal marrow processes involved in the elongation of the bones. This would suggest that the “trajectorial theory” may rely on a load intensity threshold.

In terrestrial conditions, in the straight configuration – whatever the weight-to-length ratio – the stress spreads in two ways: 1) peaks of stress can be observed at the surface of the bone on the postaxial ridge; 2) numerous separated marrow processes are stressed in the inner core of the trabecular mesh. The stress spreading at the surface of the cortex essentially dissipates locally. Almost no marrow processes are involved in absorbing the energy from the cortex. However, the stress induced in the direction of the main load (Fig 8) is directly conducted through the marrow processes running longitudinally from the distal epiphysis to the proximal one. As demonstrated by Fyhrie and Carter (104), the strain energy density is minimised when the architecture of the trabecular mesh is anisotropic such that the main orientation of the trabeculae parallels the maximum principal stress. The numerous longitudinal marrow processes in the humerus of *Eusthenopteron*, oriented in the same direction as the main external load (i.e. the Ground Reaction Force, Fig 8), therefore play a major role in reducing the overall stress on the bone in this configuration. This would explain why this configuration is the least stressed in our study (Fig 13;15;17;19). This FEA would support the hypothesis proposed by Andrews and Westoll (24) that *Eusthenopteron* could probably prop its body at the bottom of the water. This may be for basking or hunting as northern pikes do, or more commonly as lungfish benthic behaviour (105).

The hypothetical bent configuration is the most stressed configuration, probably because it is the least realistic regarding *Eusthenopteron*’s locomotory behaviour. Nevertheless, even though the fin in these conditions is the most constrained (with a median von Mises stress value of 0.22 MPa, and a highest value of 32.51 MPa, when the catfish is considered as a morphological reference), these values are far from the yield strength of the bone matrix in tension (114 MPa) and breaking stress in tension (133 MPa) (49,95). The peaks of stress are located in the humeral cortex of *Eusthenopteron* but the trabecular mesh is greatly stressed in these conditions too. The stress spreads deeper in the vicinity of the peaks and, once more, greatly dissipates through the internal marrow processes (Suppl. Movie 21, 24-25). The von Mises stress value in this configuration fits in the range of extant limb-bone values which resist to compressional loading under gravity (e.g., between 0.02 MPa in the femur of the flamingo (106) and 5 MPa in the limb of the crocodile (107). This means that the humerus of *E. foordi* would have been able to handle this stress intensity to resist to terrestrial standing posture.

Because most Devonian stem tetrapods have similar body size and long-bone microanatomy to *Eusthenopteron* (16,108), we suggest that FEA should be applied on 3D models of long bones considering their actual trabecular mesh. This would give 1) a more exact stress intensity, and 2) information on the internal stress dissipation within long bones exposed to external loads. This is crucial for a proper interpretation of the lifestyle and locomotory behaviour of these Devonian stem tetrapods for which we have no extant representative for applying strict comparative anatomy.

### Role of the muscles in stress dissipation at the surface of the humerus of *Eusthenopteron foordi*

In lift-based swimming conditions, the von Mises stress is mostly spread in the proximal region where the muscle attachments for shoulder elevation/depression are located (105). The highest peaks of stress are highlighted at the location of the supracoracoideus attachment and the postaxial margin process connecting to the latissimus dorsi. In drag-based swimming conditions, the von Mises stress is distributed on the entire surface of the humerus with peaks of stress in the coracobrachialis fossa, on the deltoid and supinator processes. These muscles are involved in the shoulder protraction/retraction and elevation/depression movements, as well as the radial/ulnar deviation (105).

Even though the muscle attachments have not been taken in consideration in this FEA, stresses increased at the location of the processes and fossae where muscles are supposed to attach. The musculoskeletal system plays a major role in absorbing the energy when impacted by forces such as the ground reaction force (109,110). The bone absorbs the stress by deforming (110) while the muscles absorb the stress by negative reaction (110–112). Muscles are known to play a double role in the maintenance of the skeletomuscular system and their role in absorbing the stress is as important as the generation of motions (113). Our FEA results would therefore suggest that the von Mises stress distributed in lift-based mode in the proximal region of the humerus would be absorbed by the supracoracoideus and latissimus dorsi, also involved in the elevation/depression movements of the fin during swimming (Fig 11). In drag-based swimming conditions, the von Mises stress is higher and therefore probably needs to be absorbed by both the muscles involved in the elevation/depression and the radial/ulnar deviation.

In terrestrial conditions, when the fin is kept straight, the stress is preferably distributed on the postaxial side of the humerus in lesser intensity, even when loaded with the weight of the catfish. The highest peak is located at the attachment of the latissimus dorsi. In this configuration, the fin would be locked like a crutch and held by the connection of the latissimus dorsi to the animal’s back as in lungfish (105). When the fin is bent with an angle of 50° though, the stress intensity drastically increases and the stress is distributed on both sides (Fig 16;18). It could therefore be absorbed by all the muscles involved in the motions of elevation/depression, protraction/retraction and radial/ulnar deviation. These results show that the von Mises stress distribution in the humerus of *Eusthenopteron* is not random and reflects the muscular activity in motion and energy absorption depending on the load applied in different hypothetical environmental conditions.

In conclusion, both the trabecular and muscular patterns play a major role in absorbing and dissipating the stress induced by the external main load on the humerus of *Eusthenopteron* in both swimming and standing conditions. This FEA shows that the energy would be absorbed by different muscles depending on the load. The bent configuration in terrestrial conditions would be the most demanding in terms of energy absorption and stress dissipation through the muscles and trabeculae though, but despite this, the von Mises stress value is in the range of extant tetrapod limb-bone values resisting to compressional loading under gravity (106,107). This shows that the combination of the musculature, morphology and microanatomy of the humerus of *Eusthenopteron* would be able to withstand the Ground Reaction Force in terrestrial standing posture. This suggests that the muscular component of the musculoskeletal system may have played a major role in both motion and energy absorption over the tetrapod terrestrialisation, especially as the girdle and limb musculature drastically enlarged over the fin-to-limb transition (27). The great sponginess in the long bones of stem tetrapods would therefore not have been an obstacle to terrestrial locomotion.

### Evolution of limb-bone marrow processes and their role in tetrapod terrestrialisation

Cancellous bone forms through endochondral ossification (58). During this, marrow processes invade the growing zone located in the long-bone metaphysis (18). These marrow processes provide enzymes secreted by the marrow cells to degrade the cartilage and leave room for the formation of bone trabecular tubules through endochondral ossification (114). In *Eusthenopteron*, these tubular marrow processes form an enclosed network running proximo-distally within the humerus (15). This gives a strong anisotropy to the trabecular mesh. Although primarily related to the growth of the humerus, the bony wall of these marrow processes also participates in conducting and/or dissipating the stress in swimming conditions, but most of all in standing posture in hypothetical terrestrial conditions (see above Discussion sections). This secondary function would therefore be an exaptation that probably played a major role in tetrapod terrestrialisation. Our FEA simulations on the (1.8-cm-long) humeral microanatomy of *Eusthenopteron* – based on the catfish body mass (i.e. 2.35 kg) – proportionally reflect the load imposed by the body weight of a stem tetrapod like *Titktaalik* on their humeri (with a humerus length = 7cm, (19); estimated body mass = 7 kg, (3)). This suggests that the numerous tubular marrow processes running longitudinally within the extended spongiosa in the limb bones of stem tetrapods (e.g., (11,16,17,46)) may have been key skeletal components to develop terrestrial locomotion rather than an obstacle to weight bearing, as previously thought (5,6,11). Of course, this trabecular configuration has a weight limit and this probably is a contributing factor for the humerus (and other long bones) to evolve into elongated bones with a narrow diaphysis, a thicker cortex and a drastically different morphology to ease terrestrial motion in larger tetrapods (5,57).

## CONCLUSION

Our study shows that the von Mises stress, induced by external load on the humerus of *Eusthenopteron*, dissipates through the cortex, trabeculae and the muscles of the pectoral appendage involved in elevation and protraction. As *Eusthenopteron*’s microanatomy is similar to that of Devonian tetrapods, we expect them to share the same process of load dissipation and energy absorption through 1) cortical stress distribution; and 2) longitudinal trabecular conduction. Our FE simulations in hypothetical terrestrial conditions demonstrate that this type of microanatomical architecture could withstand the weight of *Tiktaalik* proportionally to the size of *Eusthenopteron* in standing posture. This tubular arrangement, including marrow processes originally involved in long-bone elongation, would have acquired a key secondary biomechanical function to increase the resistance and strength of the cancellous bone to external compressive load. As an exaptation, this specific trabecular architecture may have played a major role in the tetrapod land exploration about 400 million years ago.

## ACKNOWLEDGEMENTS

The authors thank Paul Tafforeau from the European Synchrotron Radiation Facility (ESRF, Grenoble) for assisting with imaging the fossil (proposal EC203). We thank Thomas Mörs at the Naturhistoriske Riksmuseet, Stockholm for making the fossil material available. We are grateful to Mohammed Bazzi (Stanford University) and Jordi Estefa (Uppsala University) for their assistance and inputs with the Strand7 and R software packages. We thank Per Ahlberg (Uppsala University) for his assistance with useful discussions.

