## Supplementary data 1-25 for "Tetrapod terrestrialisation: a weight-bearing potential already present in the humerus of the stem-tetrapod fish *Eusthenopteron foordi*"

Supplementary Movie 1

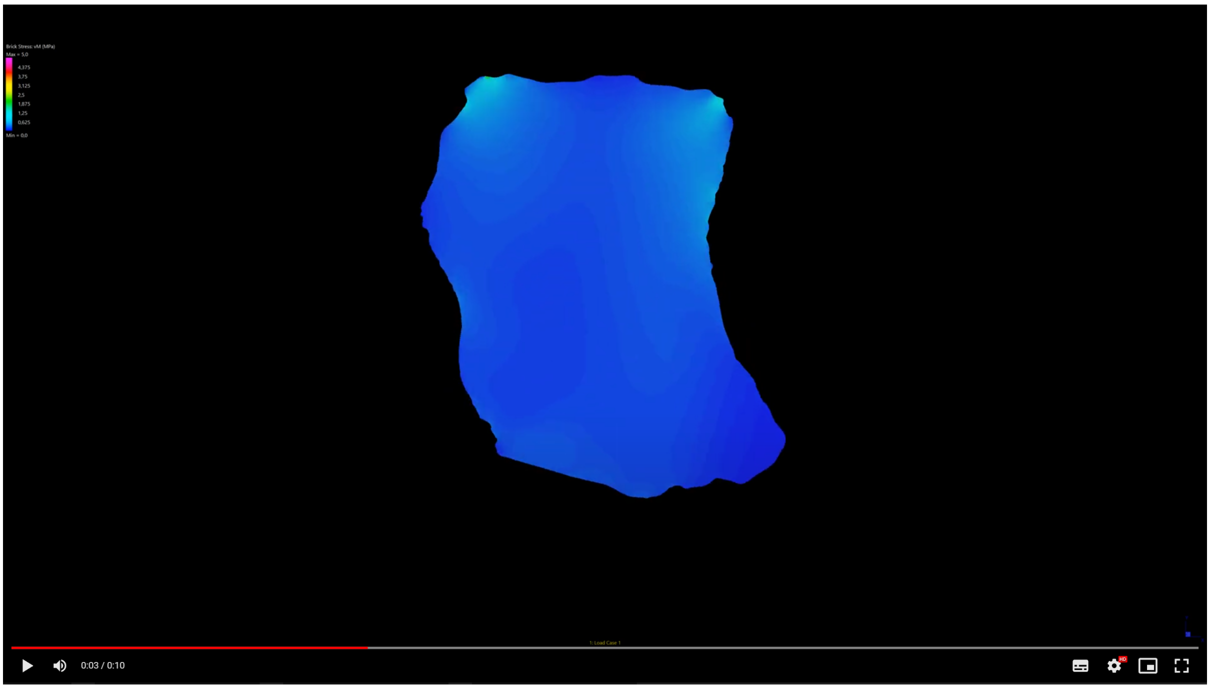


Supplementary Movie 2

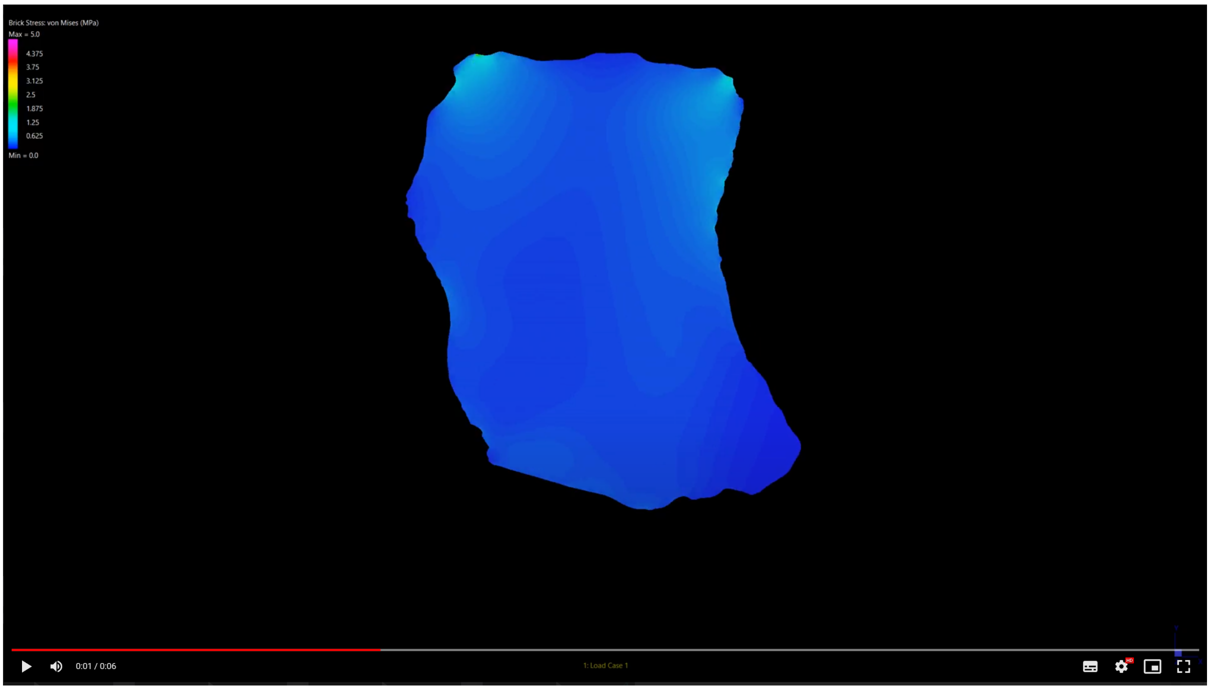

Supplementary Movie 3

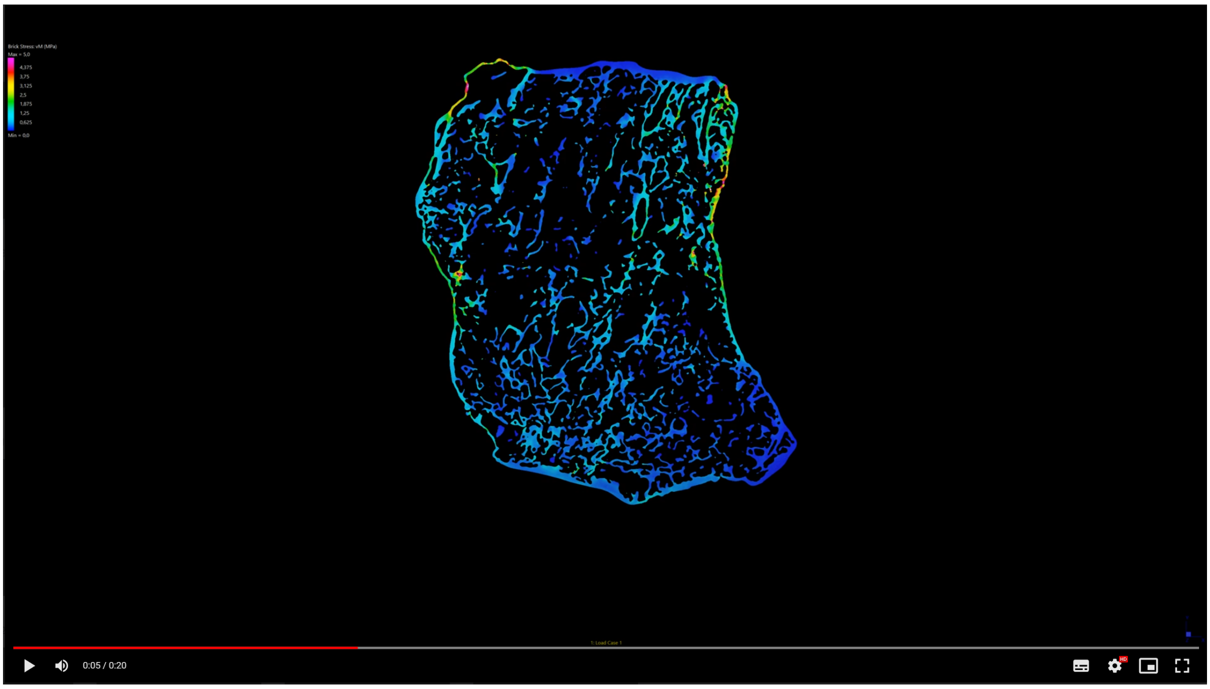


Supplementary Movie 4

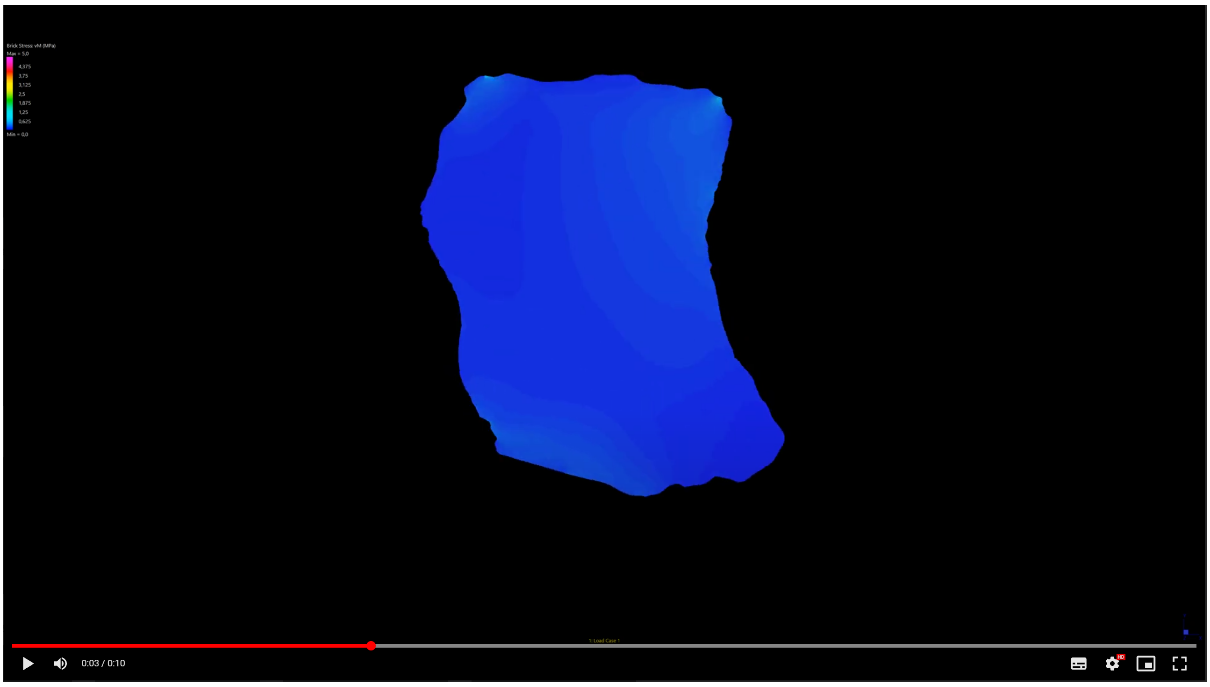

Supplementary Movie 5

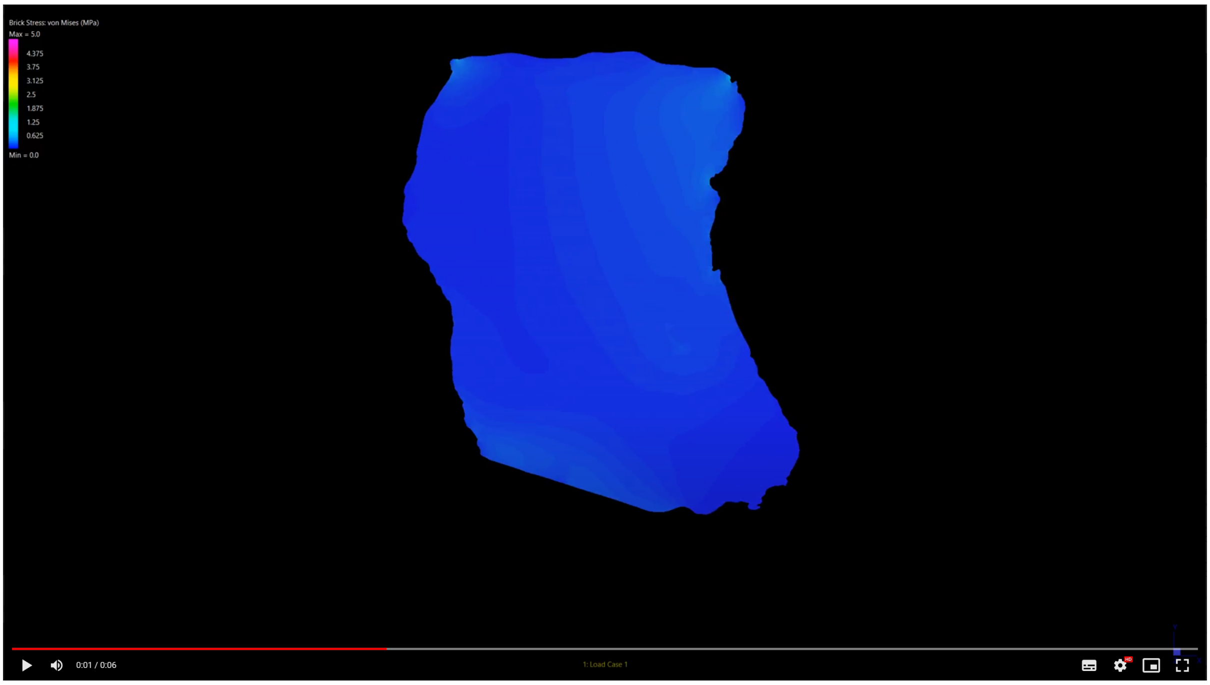


Supplementary Movie 6

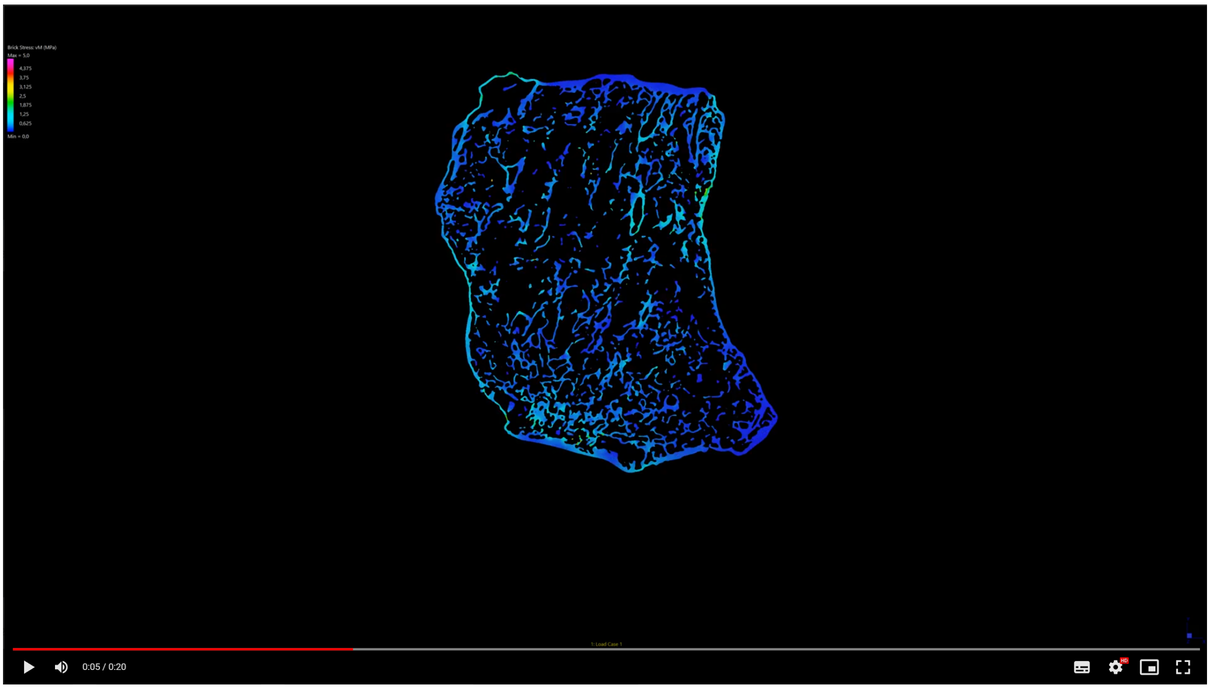

Supplementary Movie 7

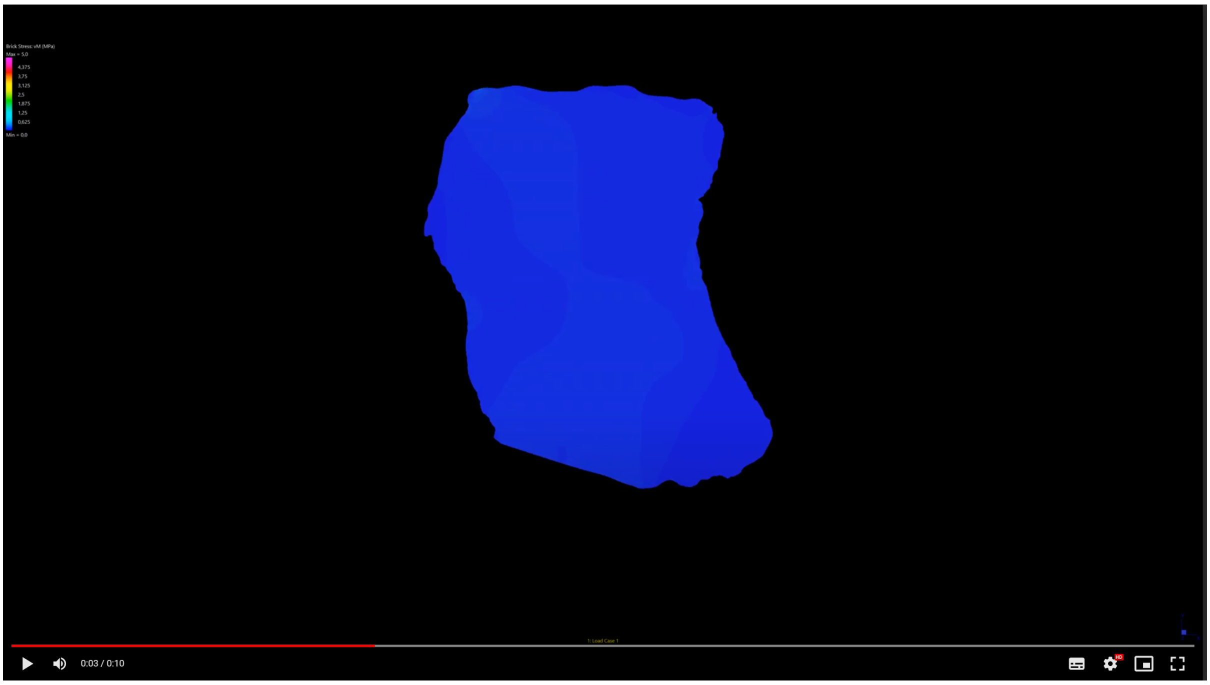


Supplementary Movie 8

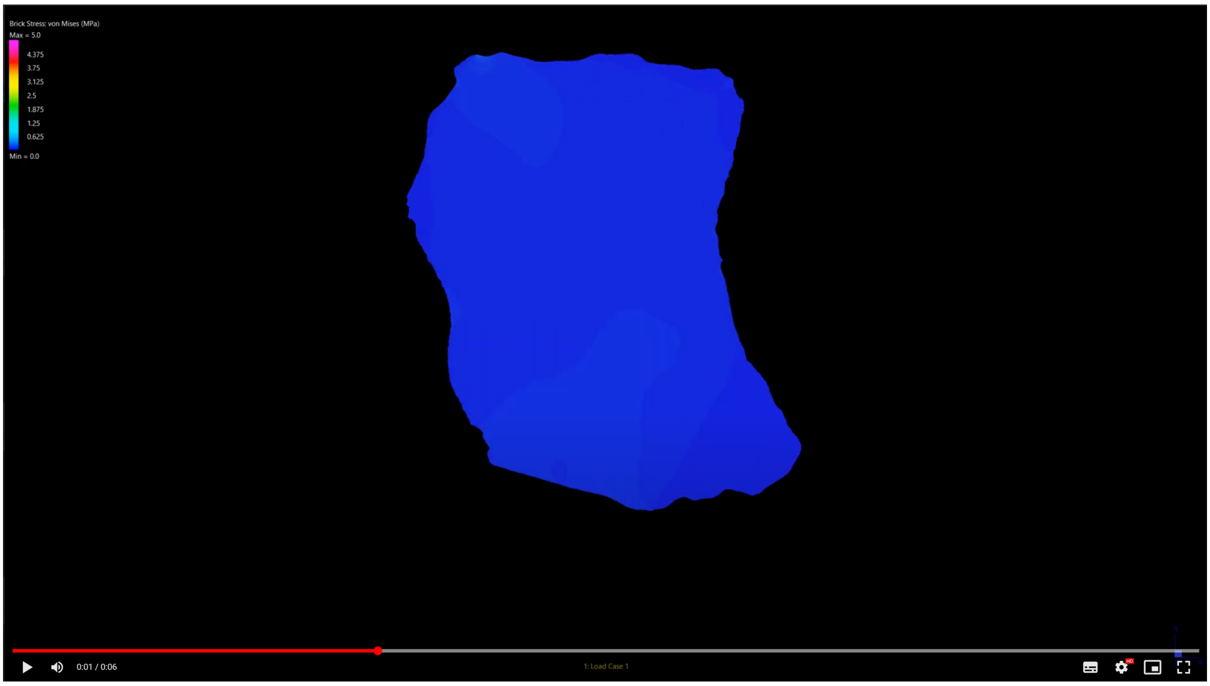

Supplementary Movie 9

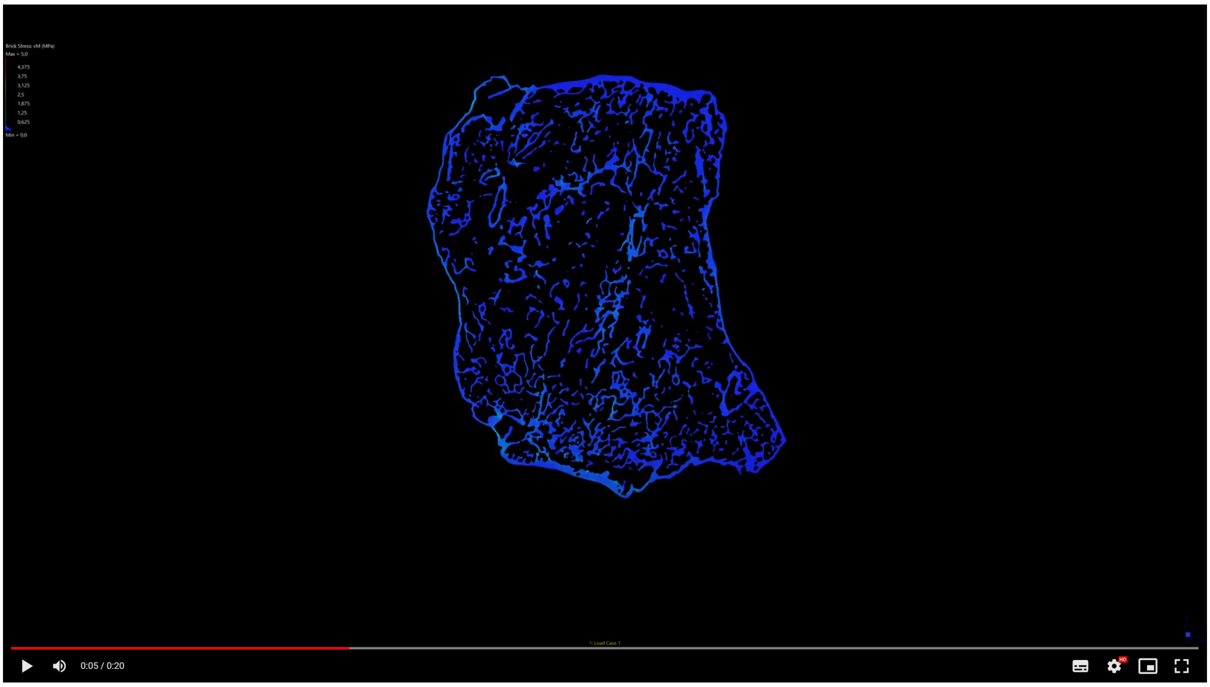


Supplementary Movie 10

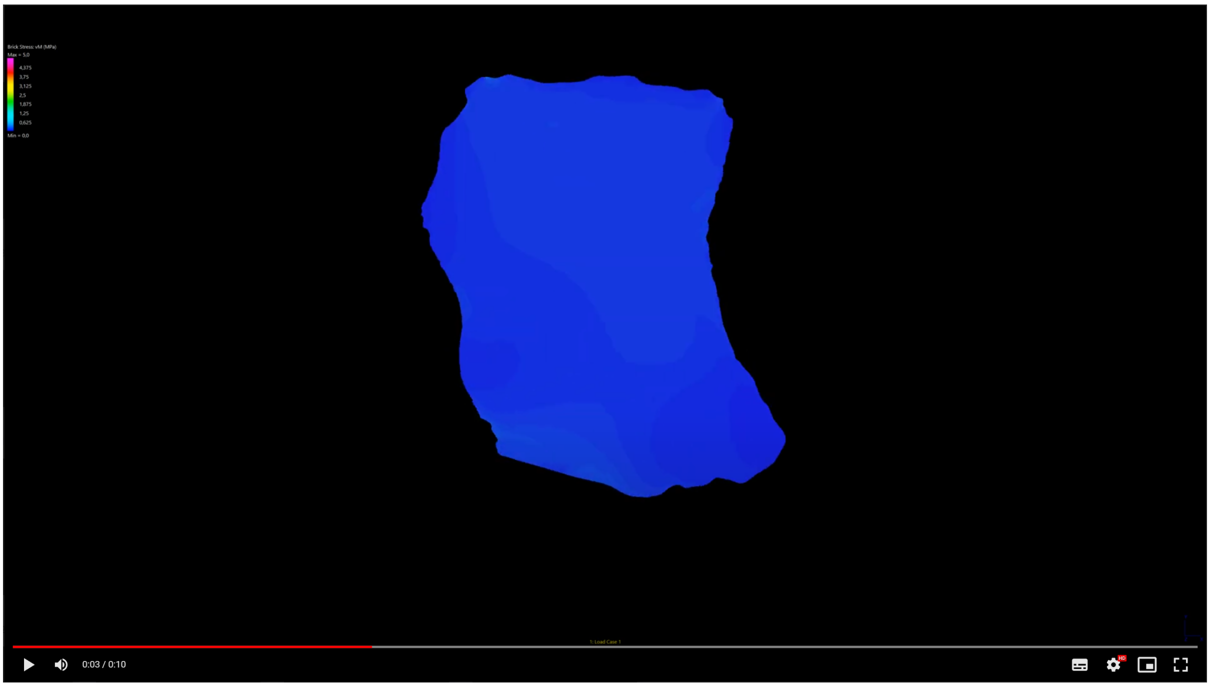

Supplementary Movie 11

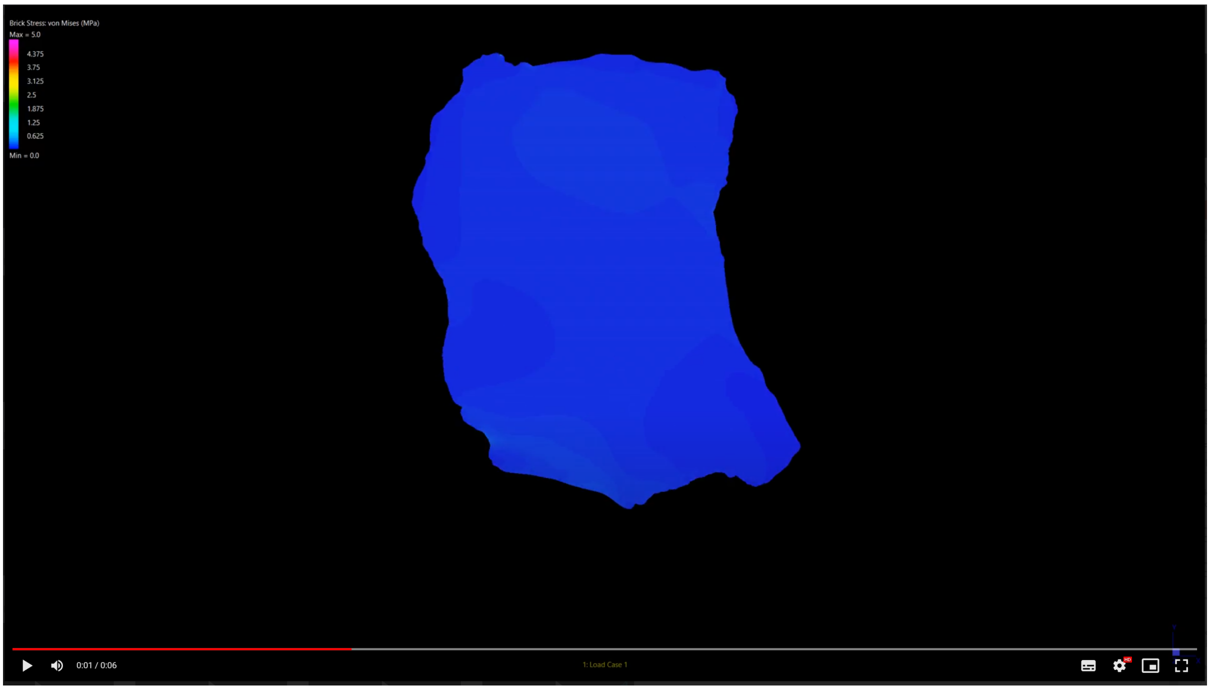


Supplementary Movie 12

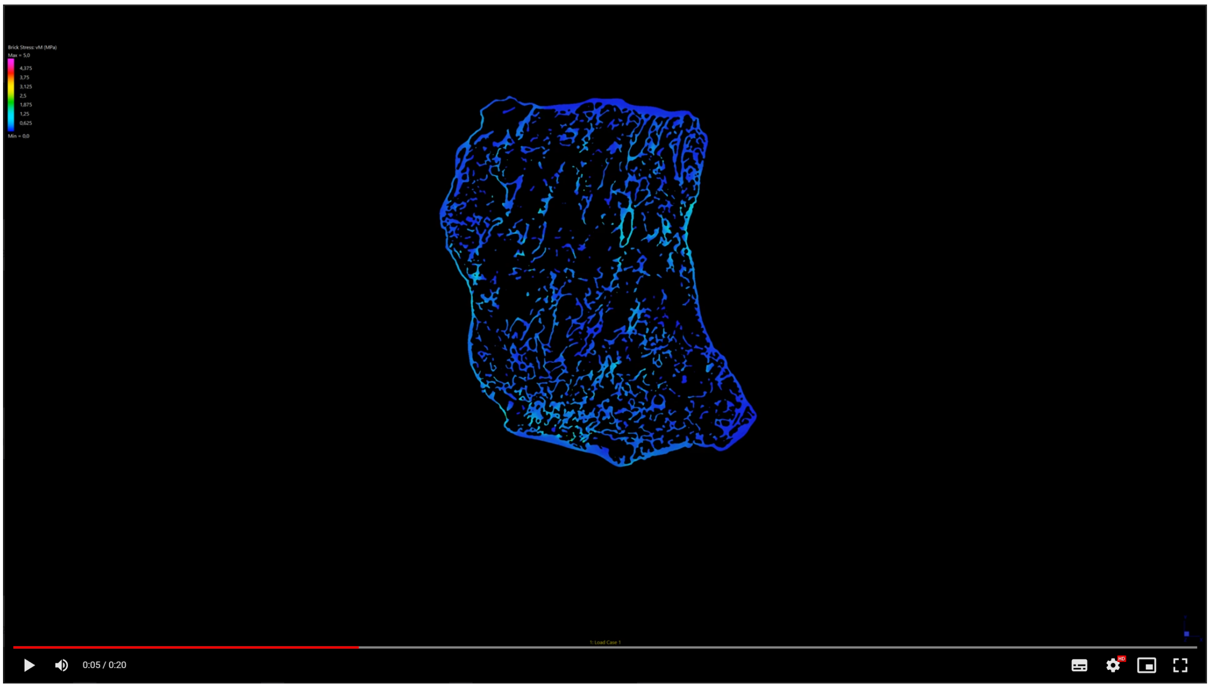

Supplementary Movie 13

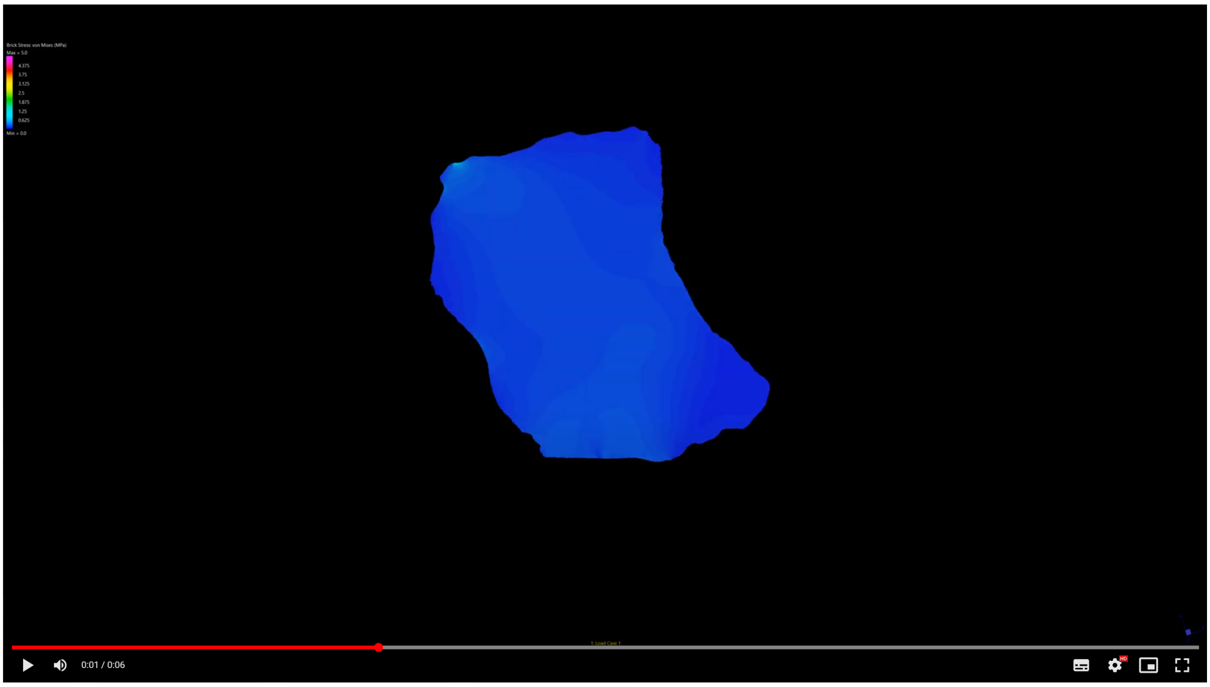


Supplementary Movie 14

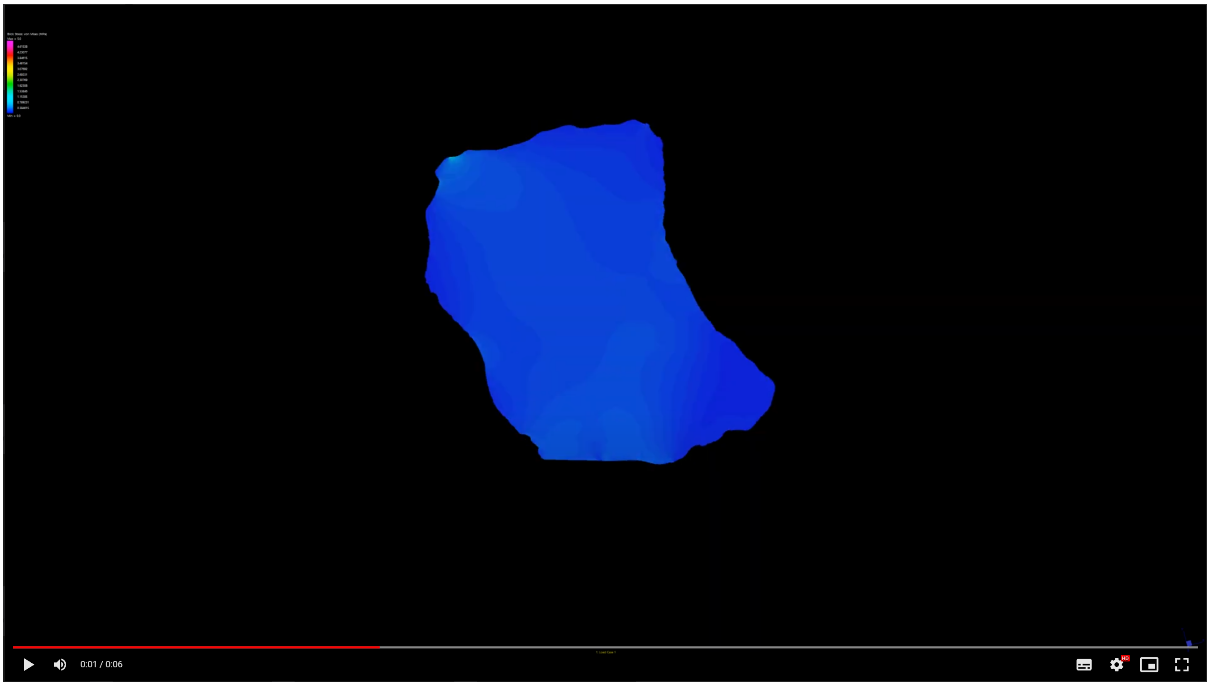

Supplementary Movie 15

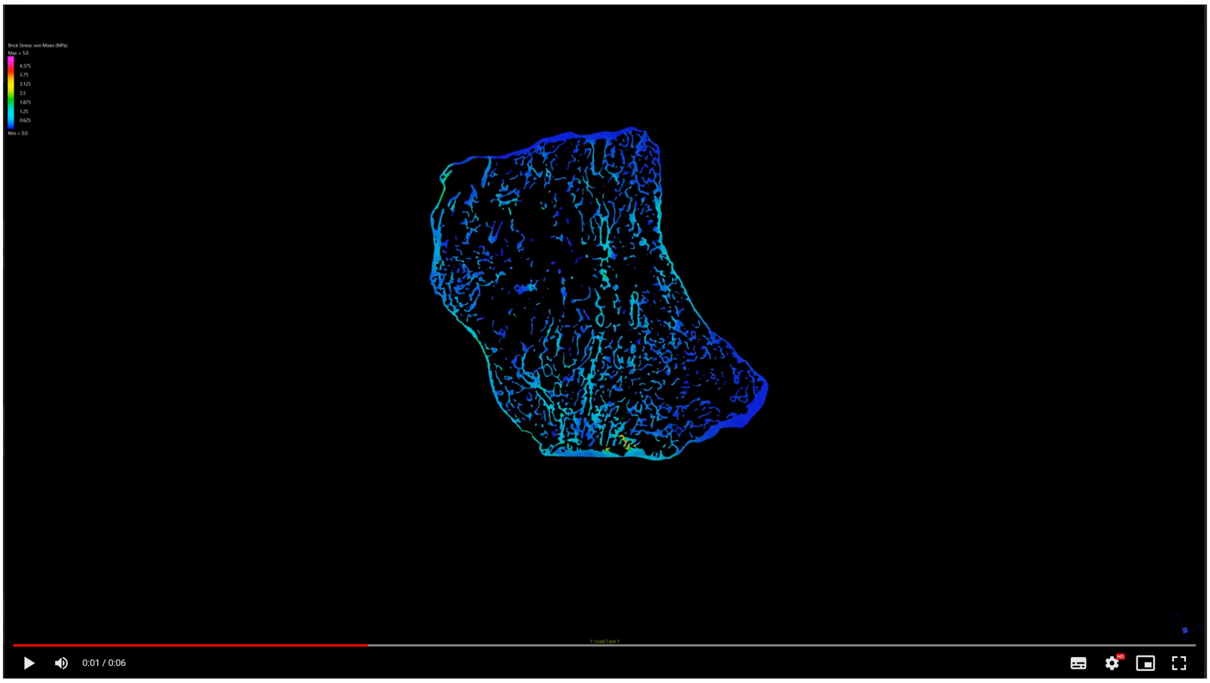


Supplementary Movie 16

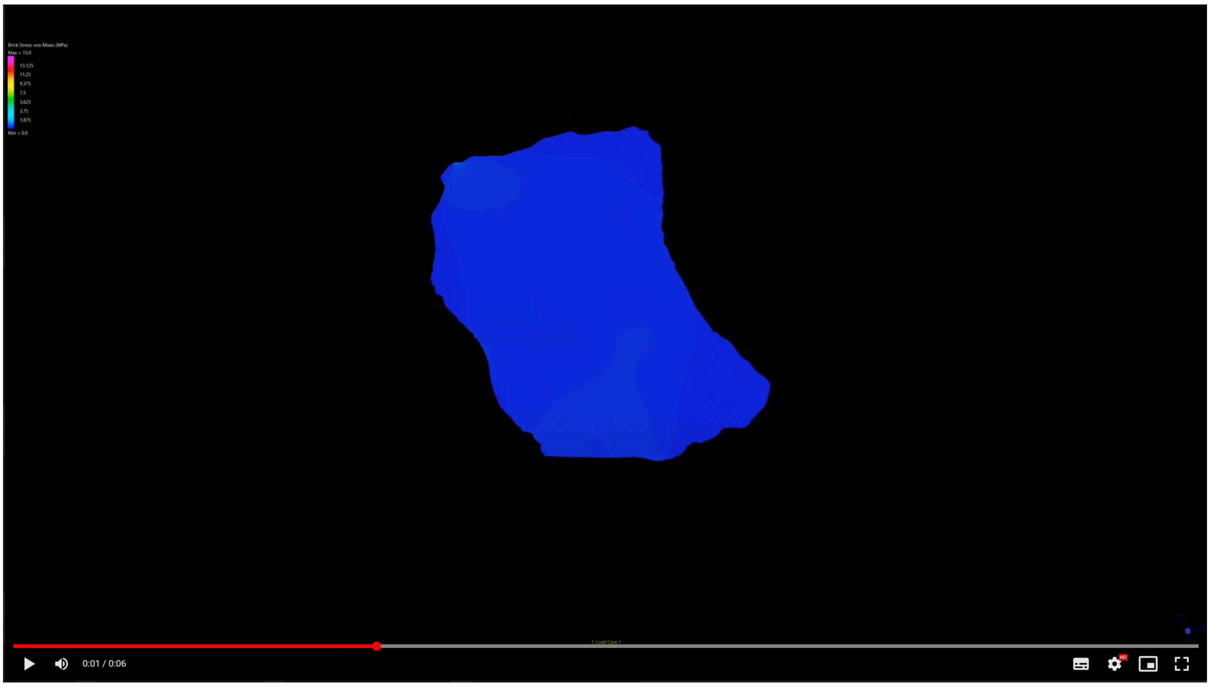

Supplementary Movie 17

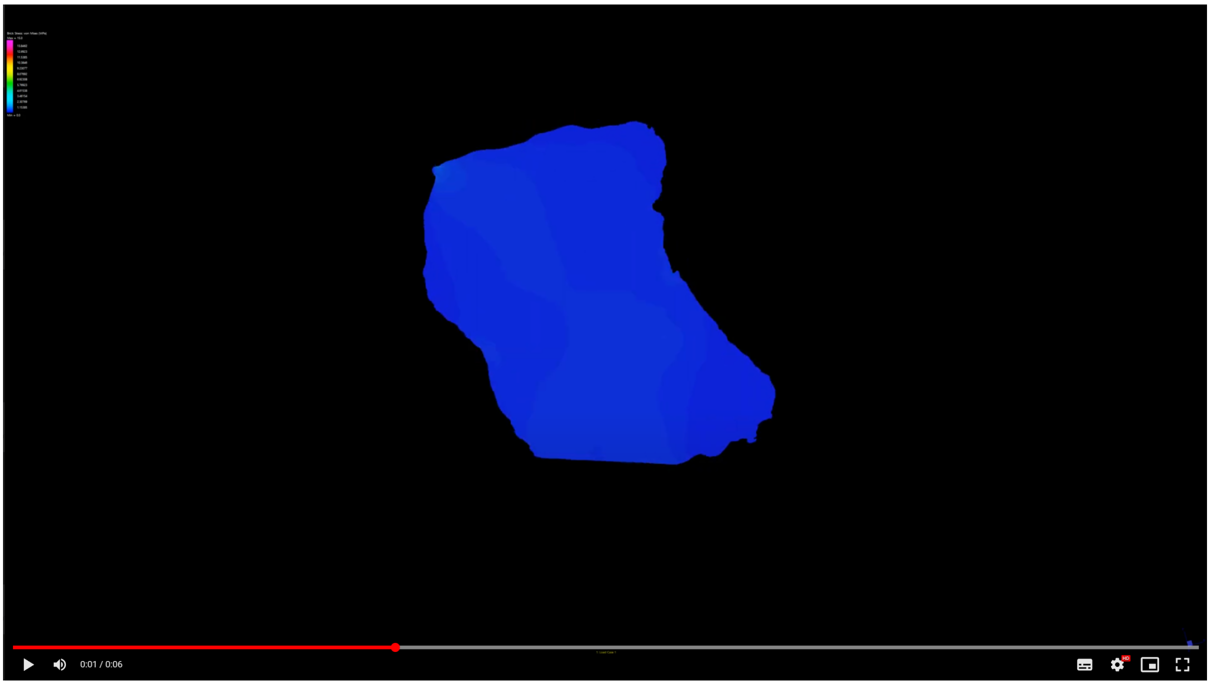


Supplementary Movie 18

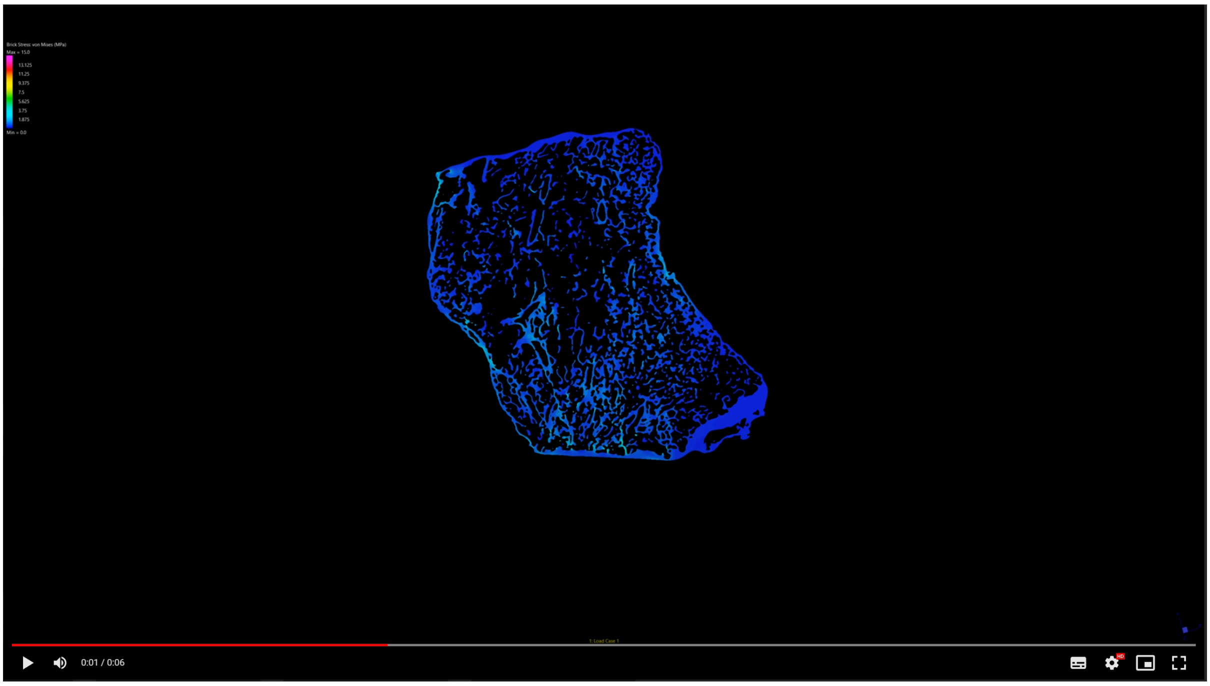

Supplementary Movie 19

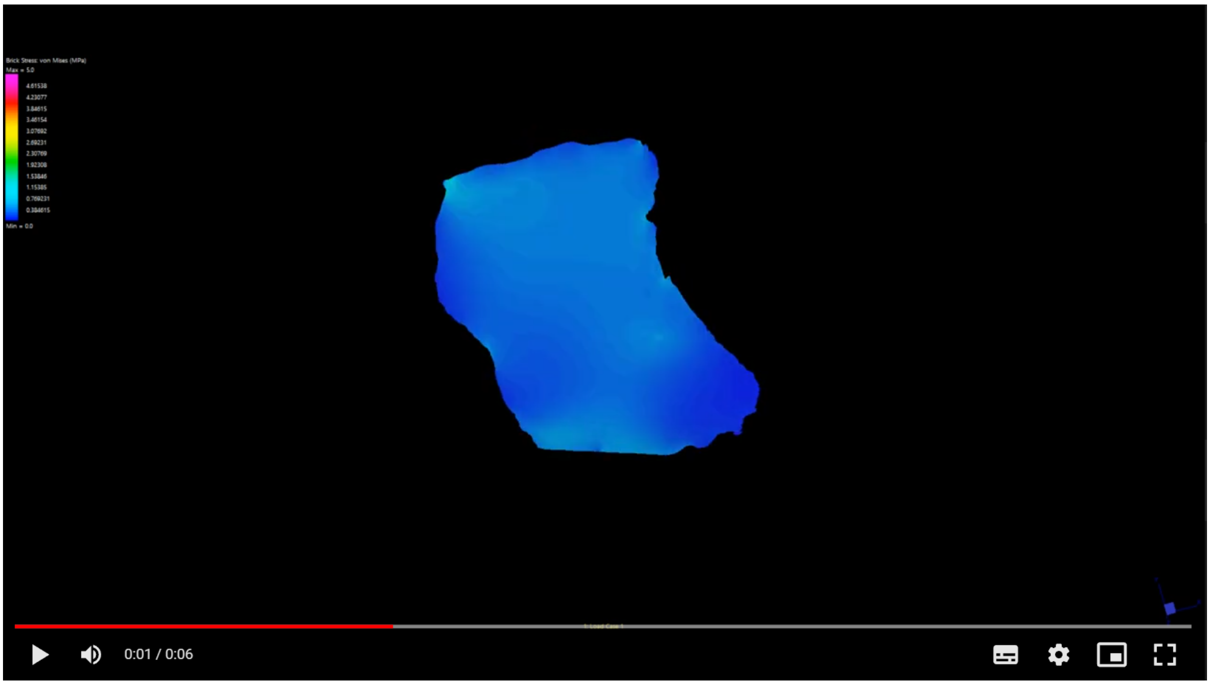


Supplementary Movie 20

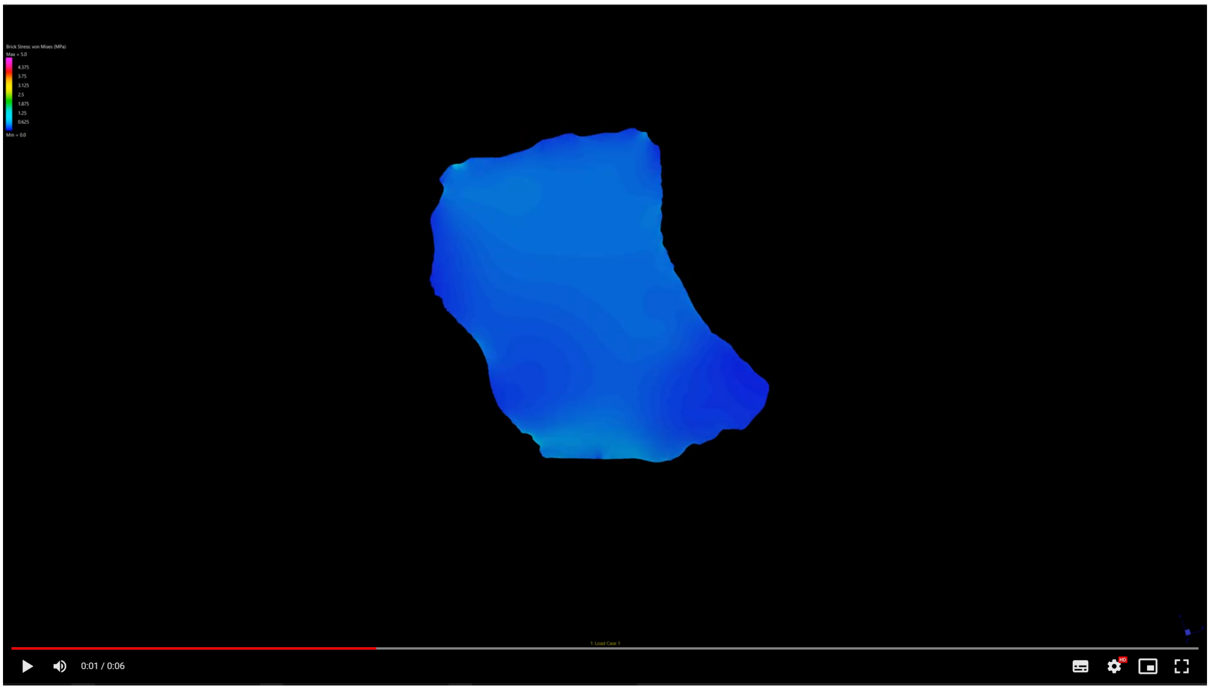

Supplementary Movie 21

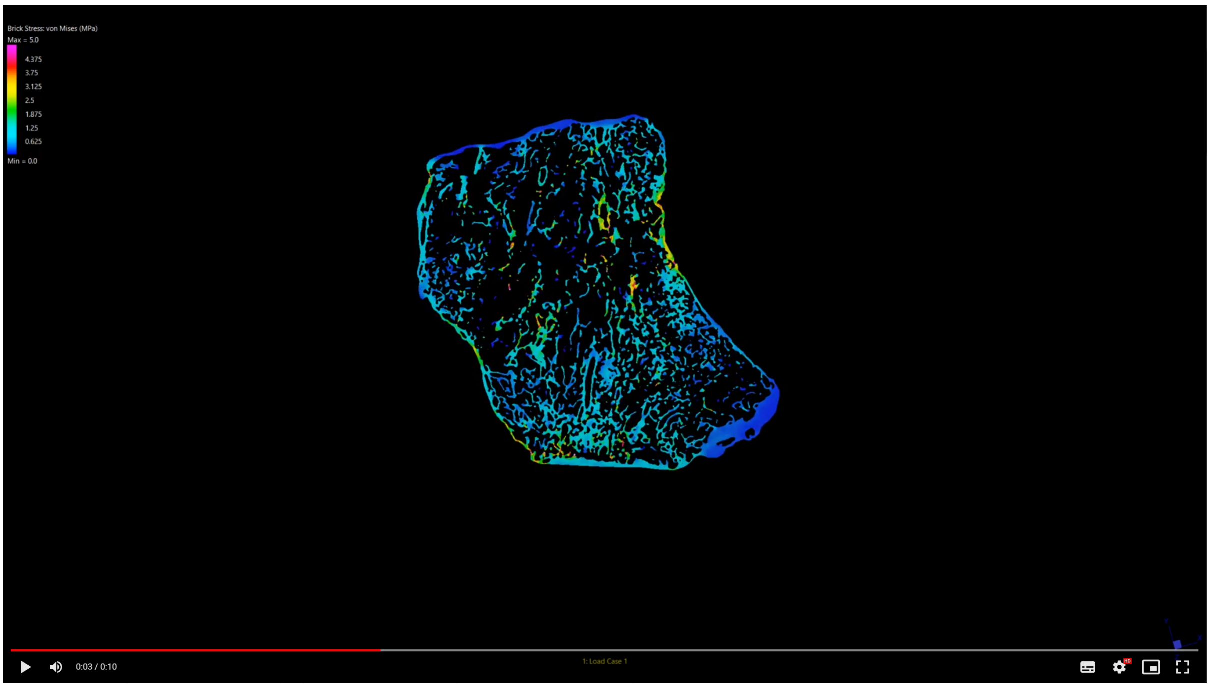


Supplementary Movie 22

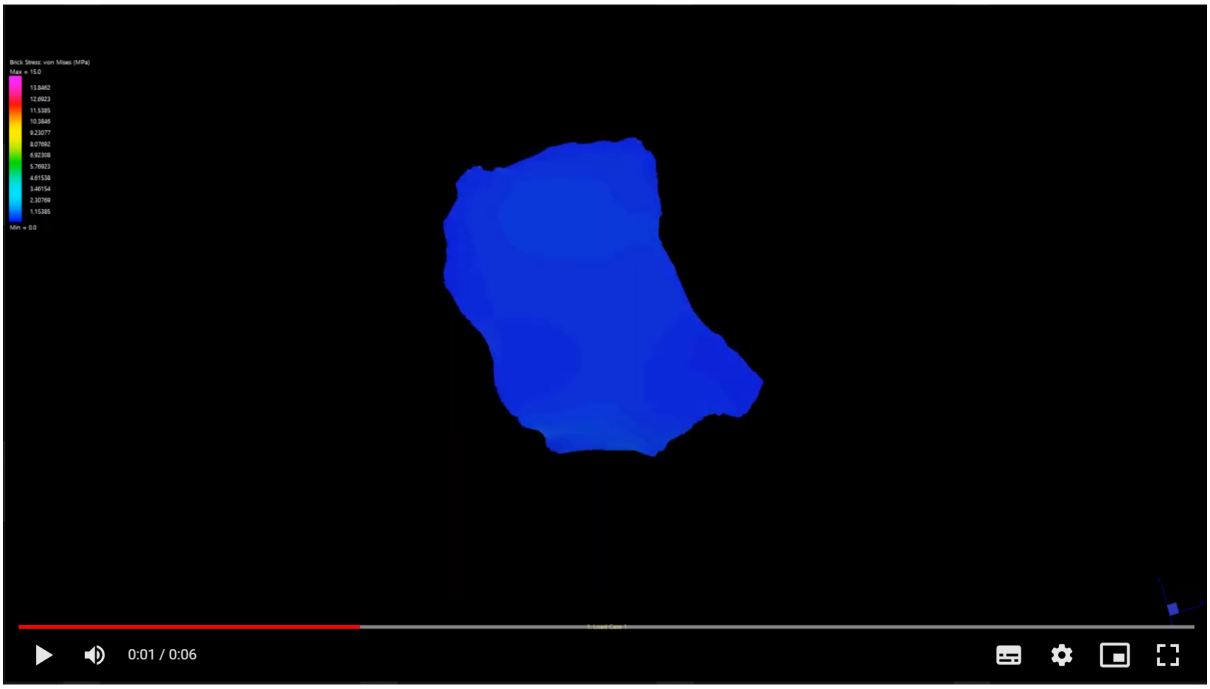

Supplementary Movie 23

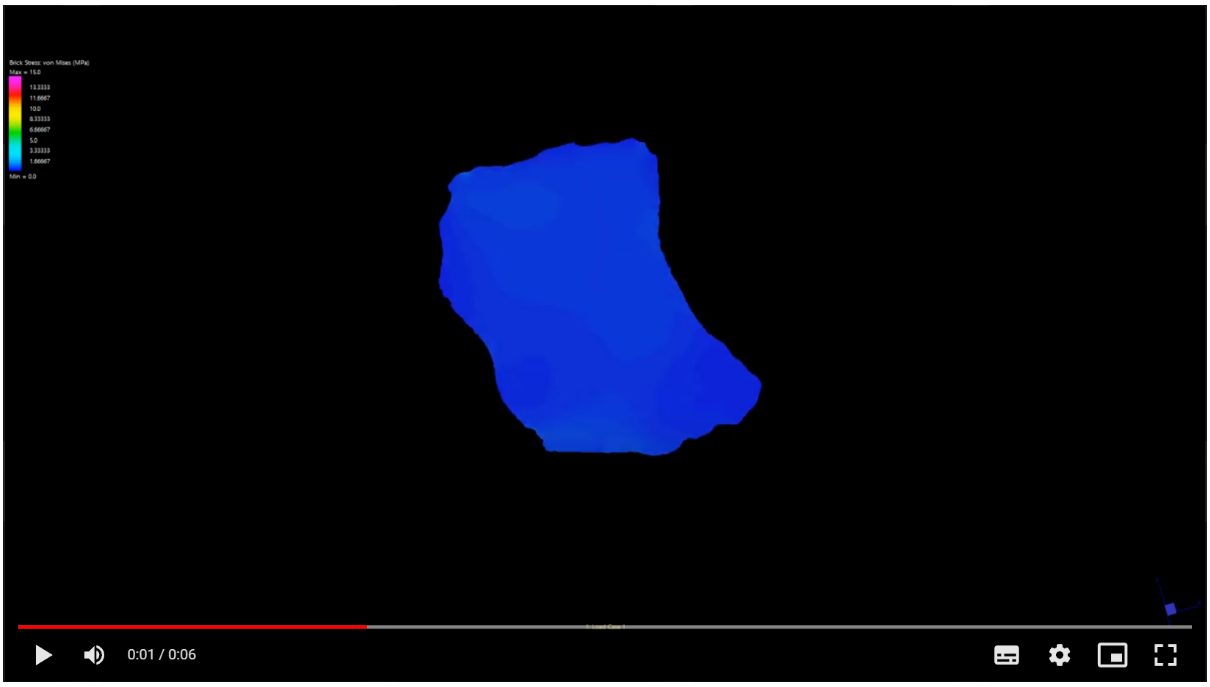


Supplementary Movie 24

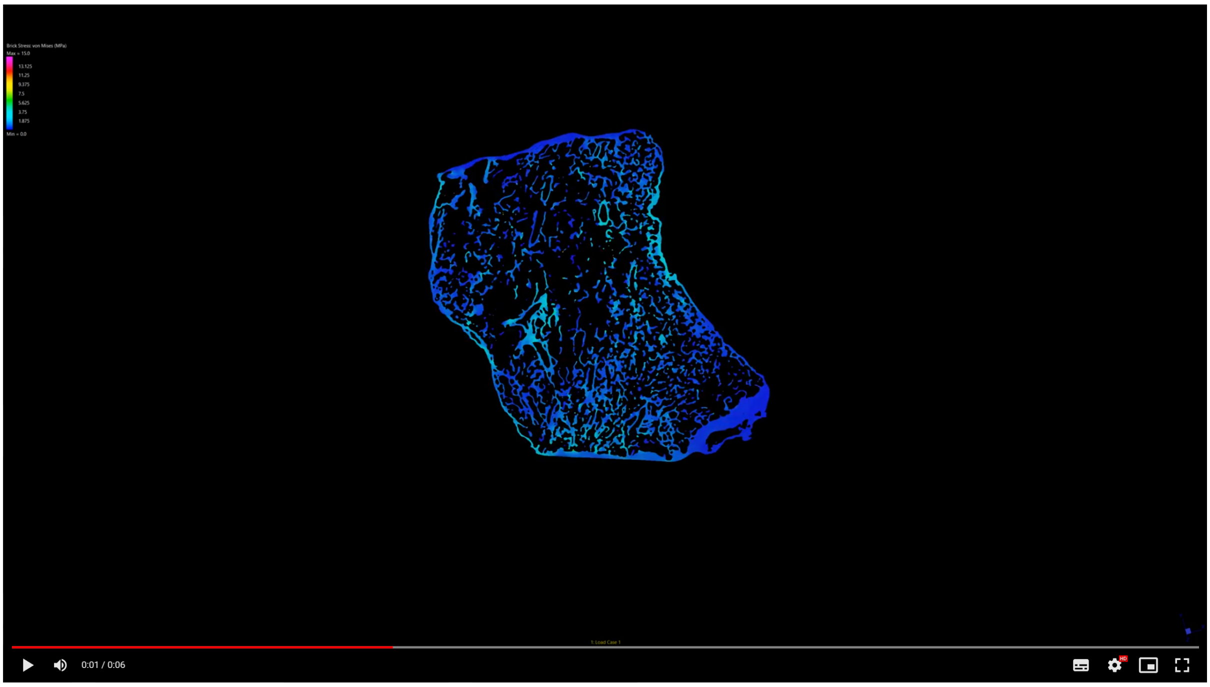

Supplementary Movie 25

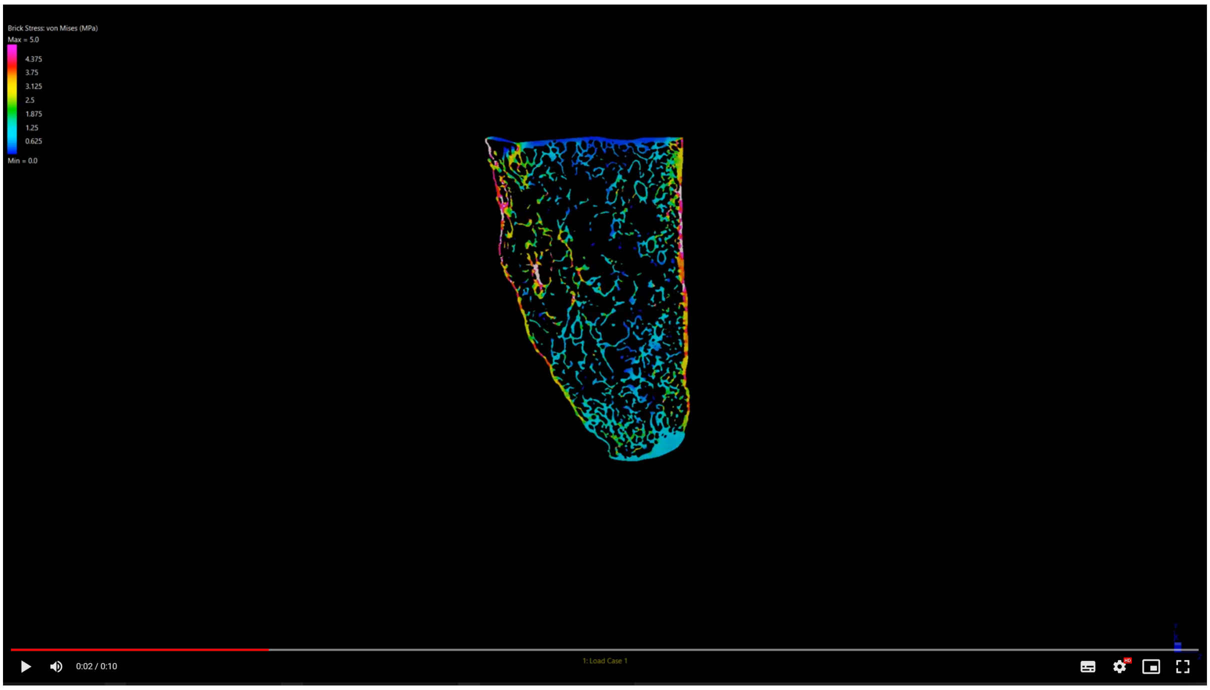
