## Supplementary figures and images for "Tetrapod terrestrialisation: a weight-bearing potential already present in the humerus of the stem-tetrapod fish *Eusthenopteron foordi*"

### Supplementary Fig S1

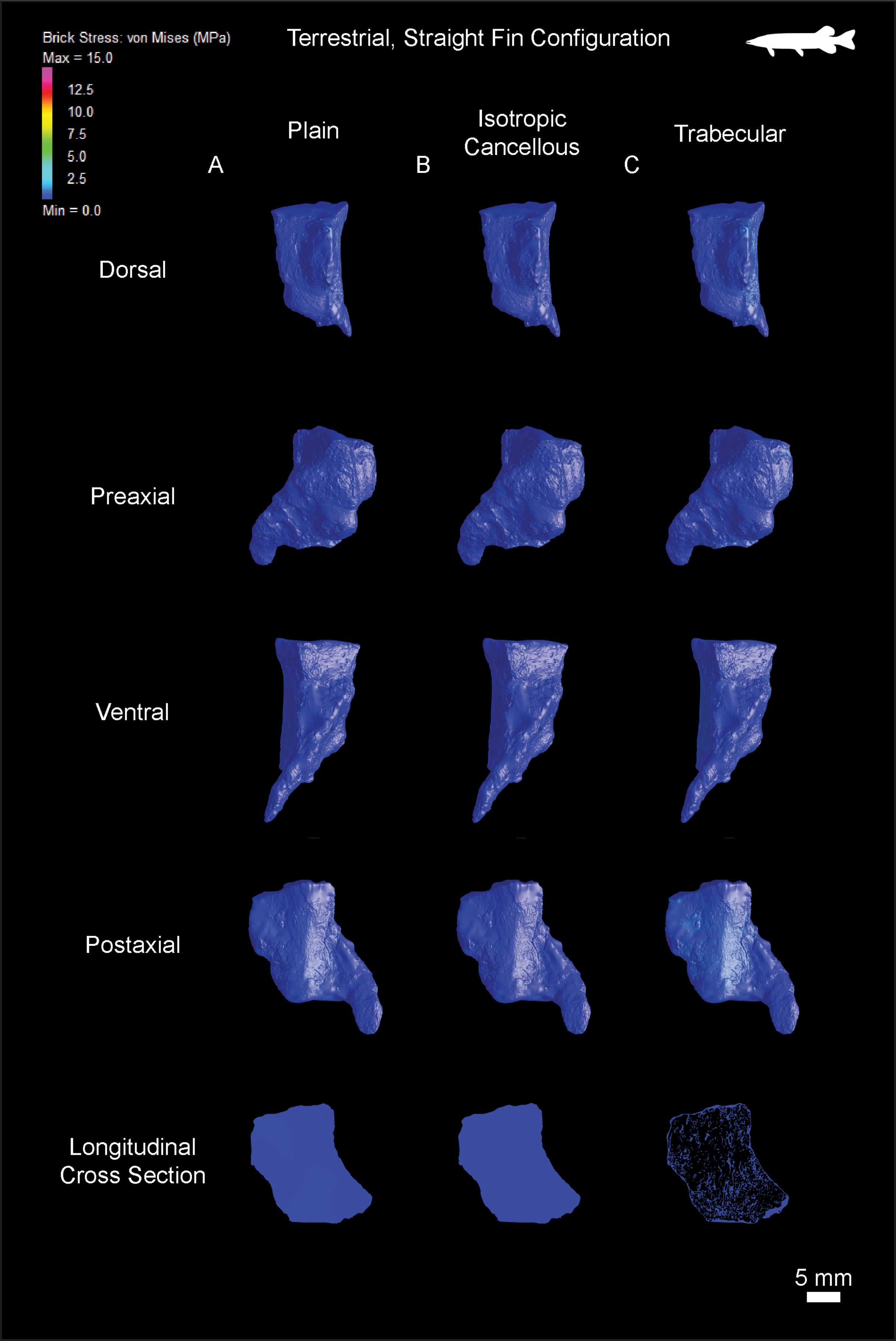

### Supplementary Fig S2

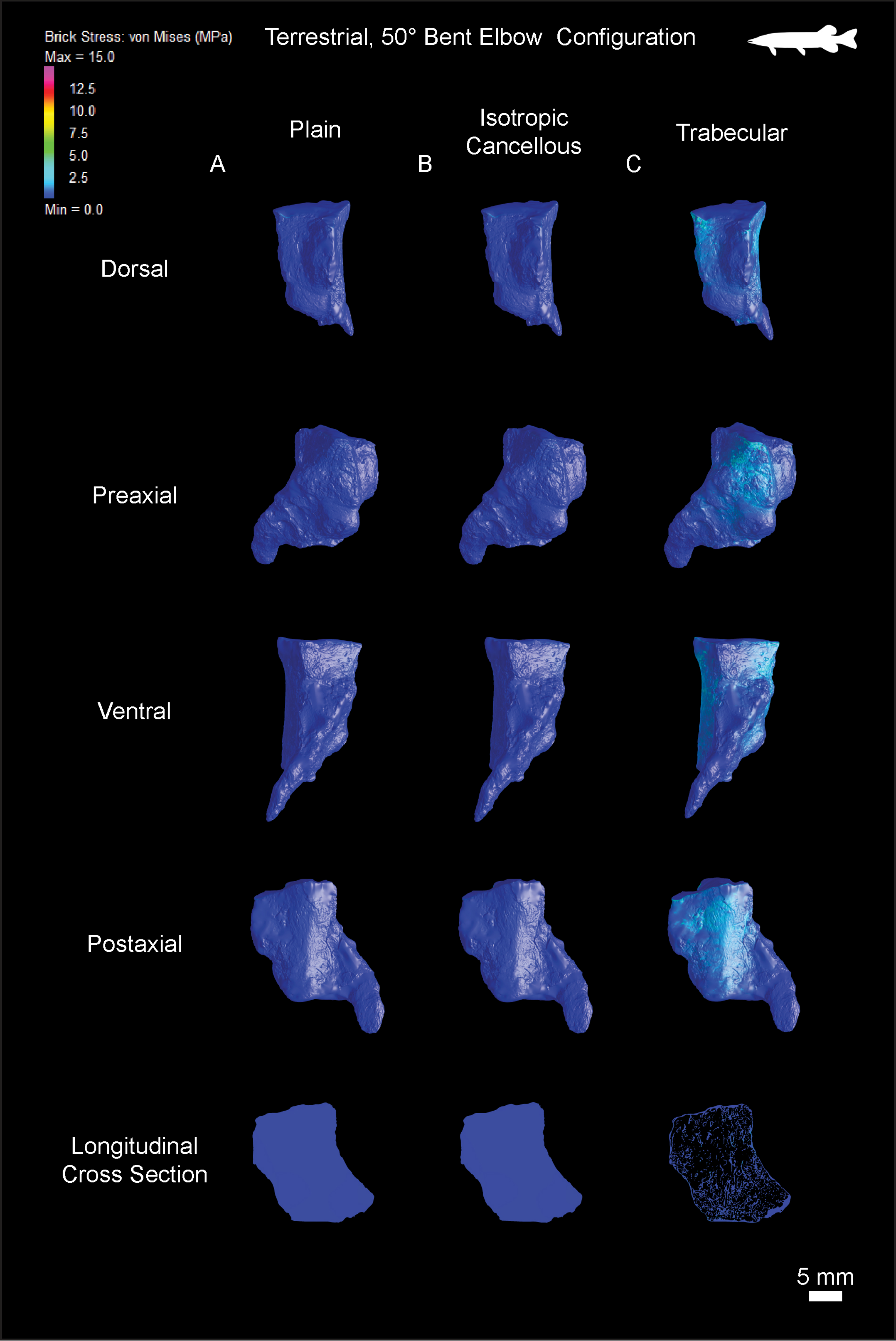

### Supplementary Fig S3

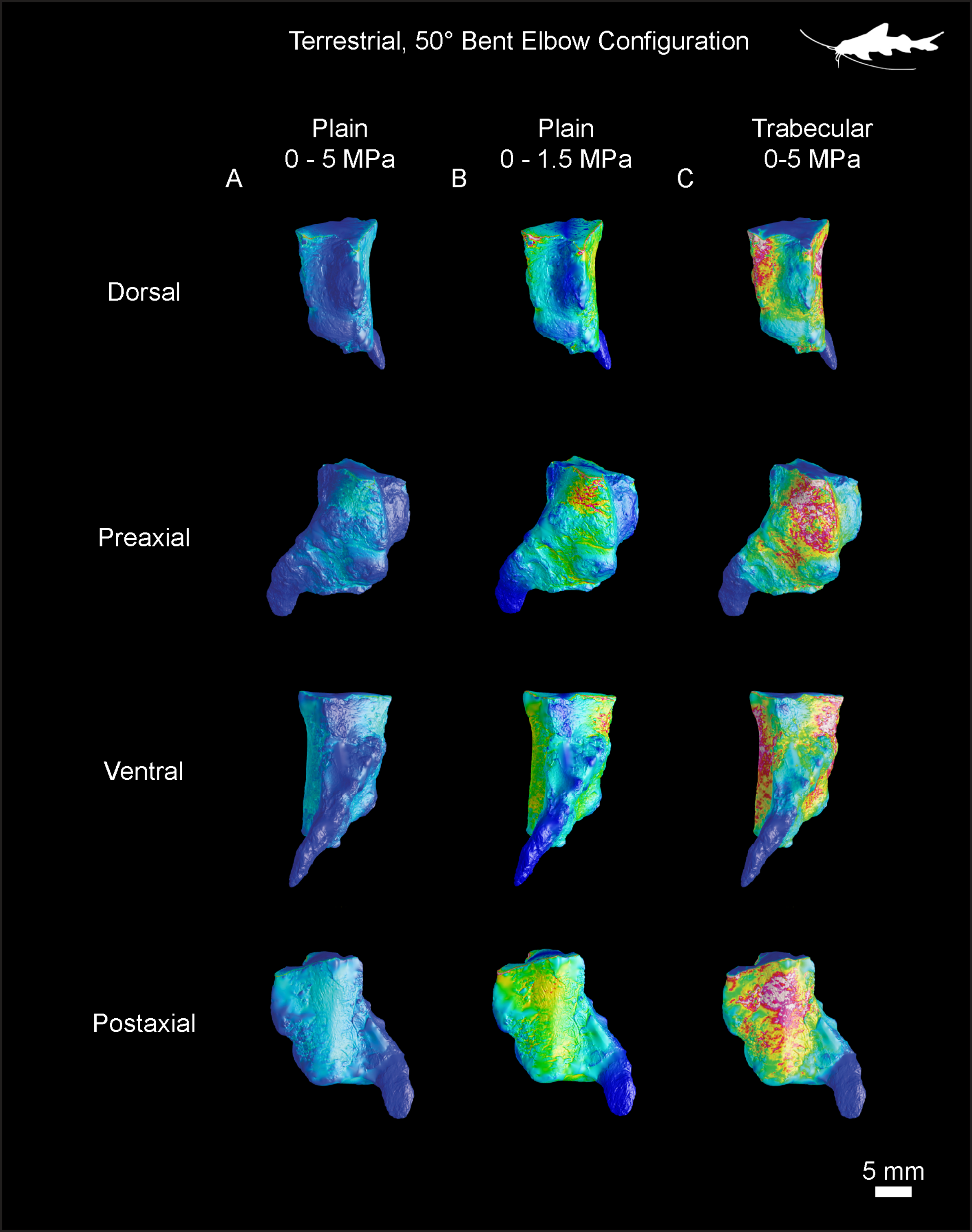
